# Brain-wide mapping of neural activity mediating collicular-dependent behaviors

**DOI:** 10.1101/2020.08.09.242875

**Authors:** Arnau Sans-Dublanc, Anna Chrzanowska, Katja Reinhard, Dani Lemmon, Gabriel Montaldo, Alan Urban, Karl Farrow

**Author notes:** These authors contributed equally to this work (Arnau Sans-Dublanc, Anna Chrzanowska).

## Abstract

Neuronal cell-types are arranged in brain-wide circuits to guide behavior. In mice, the superior colliculus is comprised of a set of cell-types that each innervate distinct downstream targets. Here we reveal the brain-wide networks downstream of four collicular cell-types by combining functional ultrasound imaging (fUSi) with optogenetics to monitor neural activity at a resolution of ~100 μm. Each neuronal group triggered different behaviors, and activated distinct, partially overlapping sets of brain nuclei. This included regions not previously thought to mediate defensive behaviors, e.g. the posterior paralaminar nuclei of the thalamus (PPnT), that we show to play a role in suppressing habituation. Electrophysiological recordings support the fUSi findings and show that neurons in the downstream nuclei preferentially respond to innately threatening visual stimuli. This work provides insight into the functional organization of the networks governing defensive behaviors and demonstrates an experimental approach to explore the whole-brain neuronal activity downstream of targeted cell-types.

## Introduction

Different behavioral tasks rely on distinct networks of neurons distributed across the brain. Insights into how specific cell-types are linked to sensation and behavior have seen some great advances through the application of molecular technologies, providing a list of critical circuit elements ^1^. On the other hand, computational understanding have been gained by comparing large-scale measurements of brain-wide activity with sensory inputs and behavior ^2–5^. However, in mammals the link between individual cell-types, large scale neuronal activity and behavior remains unclear. In the superior colliculus (SC), there is evidence for a strong relationship between individual cell-types and behavior ^6–10^. Here, we use this relationship to delineate the cell-type specific brain-wide functional networks that lie downstream of the SC.

In mice, the SC is a major hub of visual processing, where the superficial layers of the SC receive direct sensory inputs from >85% of the retinal output neurons ^11^. The retino-recipient neurons in the SC contain at least six sets of genetically identified cell-types with distinct anatomy and visual response properties ^12–16^. Different cell-types project to different sets of targets, including nuclei of the thalamus and midbrain, thereby forming a putative structural basis for the relationship between cell-types and distinct behavioral properties ^6,7,9,12,13,17,18^.

Optogenetic activation of cell-types in the retino-recipient layers of the colliculus has provided insight into the relationship between output circuits of the SC and behavior. For instance, activation of neurons that project to the pulvinar (LP) has been shown to induce arrest behavior in mice ^7,17^, while activation of neurons projecting to the parabigeminal nucleus (PBG) leads to flight behavior ^7^. These optogenetically induced behaviors resemble the reactions of mice to visual stimuli that mimic avian predators ^19,20^. These experiments suggest that activation of neurons early in the visuo-motor circuits of the SC leads to downstream activity and behaviors that are comparable to the network activity and behaviors triggered by natural visual stimuli. But our view of these circuits remains limited to the specific circuits that have been investigated to date.

In rodents, combinations of cell-type specific stimulation and whole brain recordings using functional magnetic resonance imaging (fMRI) have provided insights into the relationship between cell-types and brain-wide network activity ^21,22^. In relation to the SC, fMRI studies in humans have provided evidence that the pathway linking the SC to the amygdala (AMG) via LP is involved in the processing of visual threats ^23–25^. However, recording techniques such as fMRI suffer from limited resolution, which makes it difficult to clearly assign activity to small brain nuclei, in particular in small mammals ^21,26^. Functional ultrasound imaging (fUSi) has been developed to study brain-wide activation patterns at a spatial and temporal resolution in awake mice that makes it practical to follow neural activity in most nuclei of the brain at a resolution of ~100 μm ^27–32^. In addition, its compact size allows for parallel interventions such as optogenetic activation or local neuronal recordings in awake behaving animals ^31,33^.

By combining fUSi with optogenetics (opto-fUSi), we reveal in this study the neural networks through which information is routed after activation of different cell-types in the SC. We unravel the differences in the spatial and temporal organization of network activation depending on the cell-type and link these to differences in evoked behaviors. Opto-fUSi allows us to identify new brain areas that link sensory inputs to behavioral output, and we demonstrate that these brain areas are also activated by threat-like visual stimuli. Finally, chemogenetic manipulations unravel the potential function of one newly identified group of nuclei, the posterior paralaminar nuclei of the thalamus (PPnT), in visually triggered aversive behaviors.

## Results

### Different collicular cell-types trigger different defensive behaviors

To understand the contributions of different collicular cell classes to defensive behaviors, we optogenetically manipulated the activity of four genetically defined cell populations: 1) a population of excitatory neurons expressing CAMKII, referred to as CAMKII; 2) NTSR neurons that project to the LP, referred to as NTSR ^13^; 3) the population of parvalbumin expressing neurons (PV) that consists of local interneurons and excitatory projections to LP, PBG and pontine gray (PG); 4) a set of inhibitory neurons (GAD2) that innervates the lateral geniculate nucleus (LGN) and PBG ^13^. We restricted the expression of light-sensitive channelrhodopsin2 (ChR2) to the distinct cell classes in two ways. First, to express ChR2 in NTSR, PV and GAD2 neurons, we crossed Cre-expressing transgenic mouse lines (NTSR-GN209-Cre, PV-Cre and GAD2-Cre) with a ChR2-reporter mouse line, Ai32 ^34–37^. Second, CAMKII neurons were labeled with an adeno-associated virus (AAV) that carried ChR2 under the CAMKII promotor ^17^ (see Methods). Control experiments were carried out with Cre-negative litter mates. Histological analysis confirmed the layer-specific expression of ChR2 in different cell-types (Figure 1B), which was consistent with previous reports ^6–8,12,13,17^.

**Figure 1.**
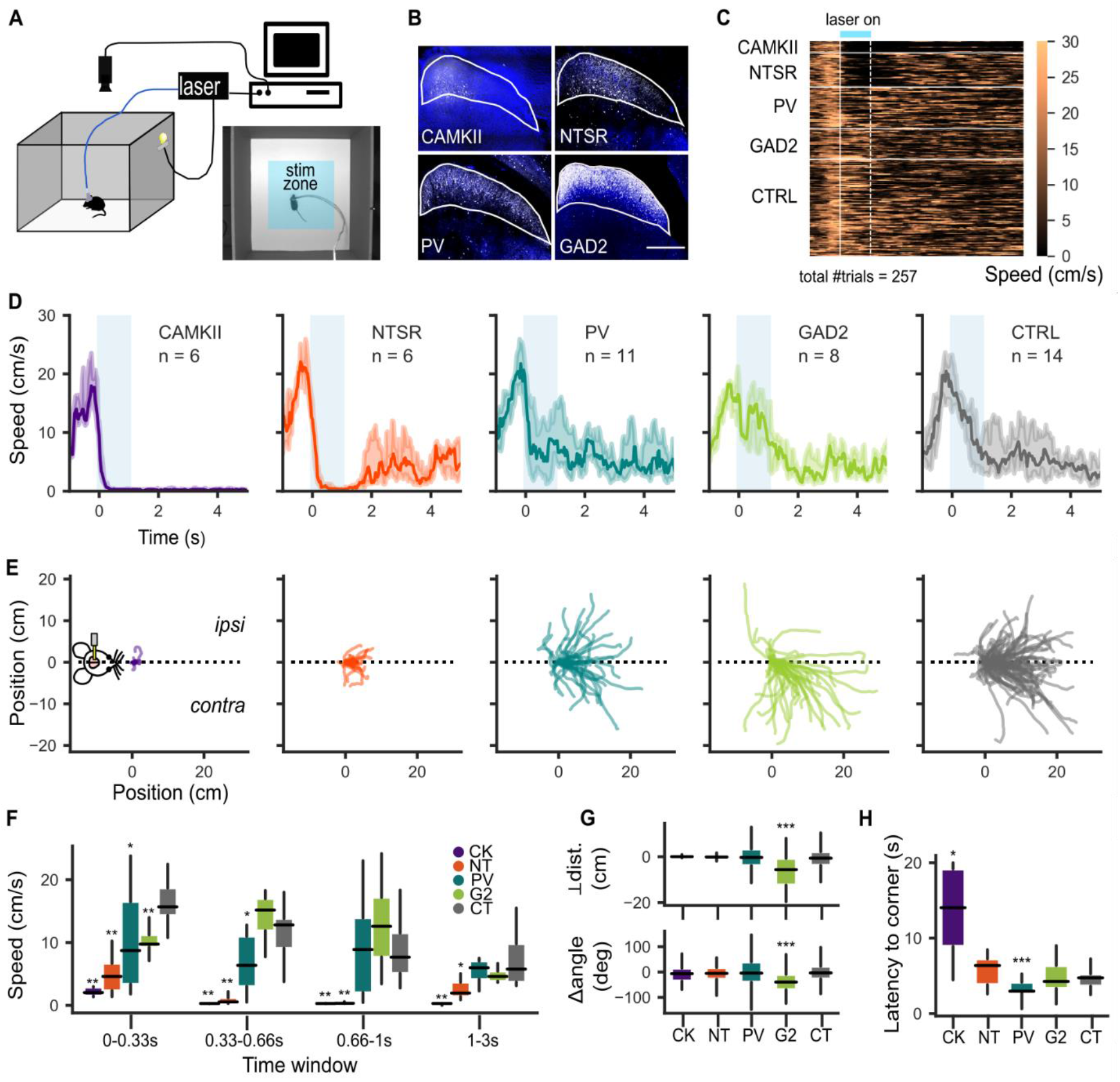
Different SC cell-types trigger different defensive behaviors. **A.** Schematic diagram of the open field setup for optogenetics and a video frame of mouse entering the stimulation zone in the center of the box (bottom right, stimulation zone is marked with blue rectangular shading). **B.** Coronal section showing expression of ChR2 in distinct cell lines. Scale bar, 500 μm. **C.** Heatmap of mice speeds during optogenetic stimulation trials. Values were obtained from the first experimental session for each animal. Horizontal white lines separate different mouse groups. Vertical solid and dashed white lines mark stimulus onset and offset, respectively. Light blue bar on the top marks the stimulus duration. **D.** Speed profiles. Each trace represents the median speed obtained from each mouse line. Shaded area represents the interquartile range. **E.** Mice trajectories during the stimulus duration (1s). Traces were aligned and rotated by the initial body position angle. CAMKII: n=6, 14 trials, NTSR: n=6, 41 trials, PV: n=11, 50 trials; GAD2: n=8, 43 trials; CTRL n=8, 77 trials. **F.** Speed quantification during chosen time windows. (0-0.33s: CAMKII p=0.0003, NTSR p=0.0003, PV p=0.026, GAD2 p=0.004; 0.33-0.66s: CAMKII p=0.0003, NTSR p=0.0004, PV p=0.03, GAD2 p=0.10; 0.66-1s: CAMKII p=0.003, NTSR p=0.0004, PV p=0.34, GAD2 p=0.10; 1-3s: CAMKII p=0.0003, NTSR p=0.007, PV p=0.38, GAD2 p=0.18) **G.** Quantification of preferred body position at the stimulation offset, represented as a change of angle (bottom; CAMKII p=0.26; NTSR p=0.31; PV p=0.45; GAD2 p=0.00002) and perpendicular distance (top; CAMKII p=0.27; NTSR p=0.29; PV p=0.38; GAD2 p=0.00002), both in reference to X axis (dashed line in E). **H.** Quantification of latency to a corner (CAMKII p=0.003; NTSR p=0.15; PV p=0.013; GAD2 p=0.49). All data points are averaged over mice, except in G where the data points are averaged over trials. Significance between control and each mouse line was tested using Mann-Whitney U-test (alpha = 0.05). Box- and-whisker plots for F-H show median, interquartile range and range. * p <0.05, ** p <0.01, *** p <0.001.

To optically stimulate the colliculus, we stereotaxically placed an optical fiber over the medial portion of the superficial layers (Figure S1F). We performed the behavioral experiments in an open field setup (50 cm x 50 cm box). At the beginning of each session mice were given a minimum of 2 minutes to freely explore the box before the experiments began (Figure 1A). Typically, animals were found to actively explore the box soon after being released into the arena, regularly moving from one corner to another (Movie S1; all the movies are listed an described in Table S1). Each trial was initiated when an animal entered the center of the arena, at which point we manually triggered the optogenetic stimulation. The stimulus consisted of blue light pulses (473 nm, 2 ms pulse width, ~9.5 – 12.5 mW/mm^2^) of either 1 s duration at 20 or 50 Hz or 4 s duration at 5 Hz. CAMKII and NTSR neurons were stimulated at 20 Hz, whereas GAD2 and PV neurons at 50 Hz. We chose the stimulation frequency based on our preliminary behavioral observations (see Methods), and the documented firing rates recorded in response to natural visual stimuli ^6–8,13,17,38^. We obtained similar results using either a 20 or 50 Hz (Figure 1) or 5 Hz (Figure S1A-F) stimulus. In the following paragraphs we focus on data obtained using the 20 or 50 Hz 1 s stimulus.

Activation of each neural population led to distinct behavioral responses, that ranged from stopping (CAMKII and NTSR neurons) to directed movement (PV and GAD2 neurons; Figure 1). To capture differences in the triggered behavior, we first looked at the speed dynamics (Figure 1C-D and Figure 1F). All experimental groups responded with a drop in speed during the first 333 ms following the start of the stimulation (Figure 1F; Mann-Whitney U-test compared with control, CAMKII: p=0.0003; NTSR1: p=0.0003; PV: p=0.026; GAD2: p=0.004), whereas control mice did not show any identifiable change in behavior (Movie S2). An animal was determined to have stopped if its speed dropped below 1.5 cm/s for at least 0.5 s. Activation of CAMKII neurons resulted in particularly long stopping events that lasted for up to 19.8 s (Figure S1H, Movie S3; median stopping duration: 9.21 s, IQR=[8.02, 12.65]), whereas stimulation of NTSR neurons resulted in mice stopping during the 1 s stimulus and resuming locomotion shortly after stimulus offset (median stopping duration 1.69 s, IQR=[1.42, 1.95]; Figure S1H, Movie S4). Activation of PV cells caused mice to slow down, but rarely led to a full stop (Figure S1I). Instead, their behavior was characterized by active movement towards one of the corners (Figure 1H, Movie S5; median latency: 2.96 s, IQR=[2.7, 4.16]; Mann-Whitney U-test compared with control, p=0.01). Animals with ChR2 expression in GAD2 neurons showed a tendency to increase their speed during the stimulation (Figure 1F; Mann-Whitney U-test compared with control, GAD2: p=0.10). Interestingly, this was accompanied by movement contralateral to the stimulated hemisphere (Figure 1F-G) that manifested itself as turning (Movie S6, Figure 1G Bottom, median Δ angle: −39°, IQR=[−70.01, −9.36]; Mann-Whitney U-test compared with control, GAD2: p=0.00002), or a whole-body drift quantified as the perpendicular to the distance traveled in the first second relative to the axis of motion before the stimulus (Movie S7, Figure 1G Top; median perpendicular distance: −5.66 cm, IQR=[−12.36,-0.72]; Mann-Whitney U-test compared with control, GAD2: p=0.00002). Taken together, these findings suggest that each collicular cell-type makes a different contribution to behavior that can broadly be characterized as defensive or orienting.

### Brain-wide functional ultrasound during optogenetic stimulation in mice

In order to assess the brain-wide neural activity that drives the different behaviors observed above, we developed a chronic preparation that allowed us to combine functional ultrasound brain imaging (fUSi) and optogenetics, in awake head fixed animals (Figure 2, A-C). fUSi reports neuronal activity indirectly by measuring changes in blood volume of the microvasculature ^32,39,40^. To accommodate the optogenetic fiber and image as much of the brain as possible, a large cranial window that spanned a single hemisphere was implanted (AP +2 to −6.5; L +1.25 to −4.5). An optic fiber was pointed at the surface of the ipsilateral SC near the midline at an angle of 56°, approached from the contralateral side (Figure 2B; Methods). All animals included in the fUSi experiments were tested for behavior before each imaging session.

**Figure 2.**
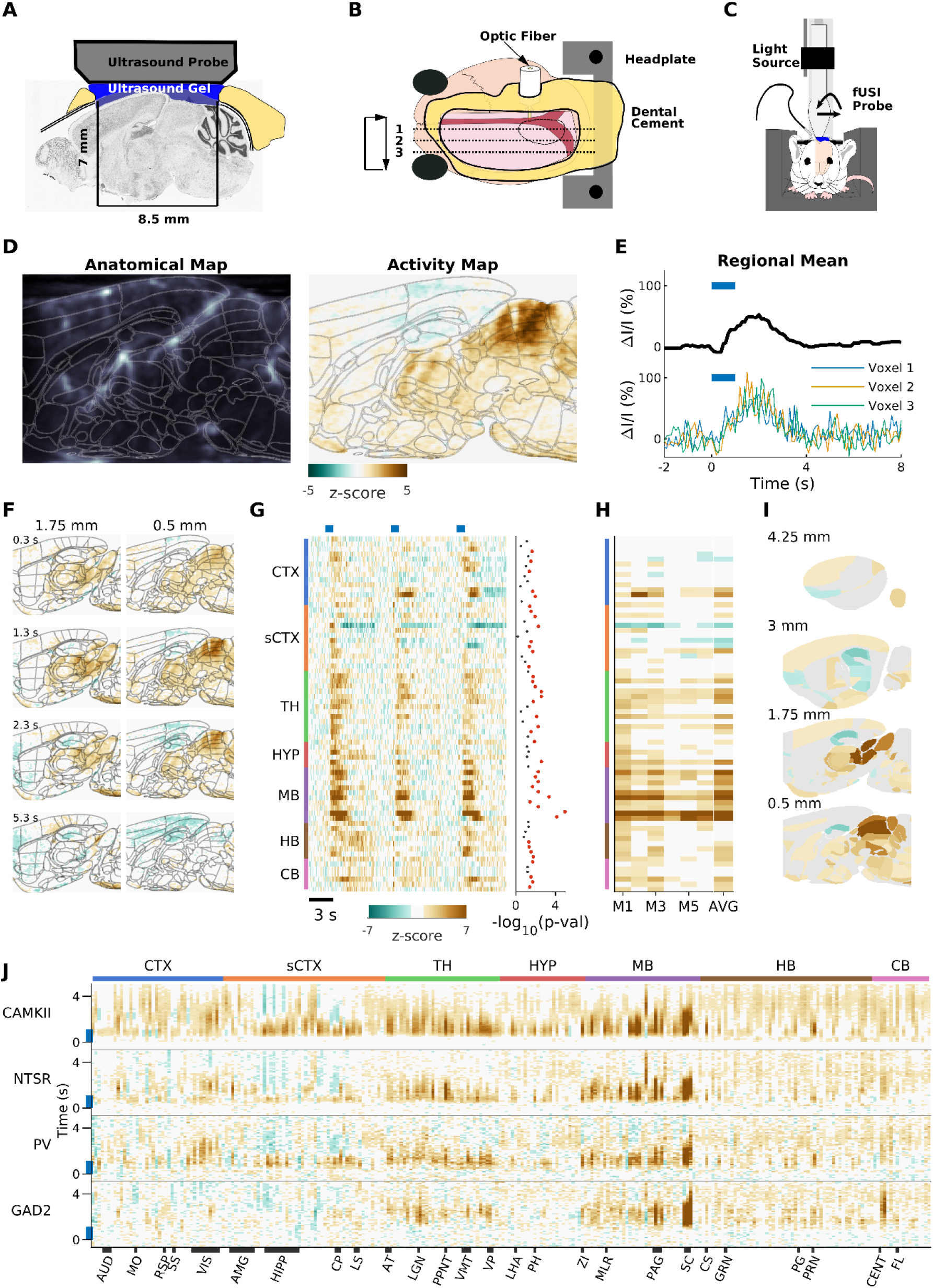
Functional ultrasound imaging of awake mice during optogenetic stimulation. **A.** Scheme of a sagittal cross-section of the chronic preparation. **B.** Top-view scheme of a chronic cranial window with implanted optic-fiber cannula inserted at 56 °. **C.** Schematic of experimental set-up for awake imaging. **D.** Left. Example sagittal section of a blood volume map registered to the Allen Mouse brain reference atlas (thin gray lines). Right. Voxel to voxel normalized response to optogenetic stimulation of plane shown in left panel, registered to the Allen Mouse brain reference atlas (thin gray lines). **E.** Bottom: relative hemodynamic response curves to the optogenetic stimulation of three example voxels in the intermediate superior colliculus. Top: mean response of the intermediate superior colliculus. Blue lines indicate duration of optogenetic stimulation. **F.** Two example sagittal planes from the activity maps of a single animal. **G.** Left: Standardized responses of a selection of 72/264 segmented areas. Mean responses are shown for 3 different mice. Response for each mouse is an average of 6 trials. Blue lines indicate duration of optogenetic stimulation. black thick line indicates optogenetic stimulation. Right: Inactive (gray) and active (red) areas colored based on significance threshold corrected for multiple comparisons (p<0.05). **H.** Mean response of each segmented area shown in G during the 2 s after the start of the stimulus for 6 different NTSR mice and the average across all mice. Areas considered not significant (p>0.05) are set to zero in the average. **I.** Projection of the average activity vector from H onto a map of the mouse brain. **J.** Average time course of each of the 264 segmented areas for each stimulated cell population. Black bars along the bottom indicate span of the labeled brain regions. CTX: cortex, sCTX: cortical subplate, TH: thalamus, HYP: hypothalamus, MB: midbrain, HB: hindbrain, CB: cerebellum AUD: Auditory cortex, MO: Motor cortex, RSP: Retrosplenial cortex, SS: Somatosensory cortex, VIS: Visual cortex, AMG: Amygdala complex, HIPP: Hippocampus, CP: Caudatoputamen, LS: Lateral septum, AT: Anterior thalamus, LGN: Lateral geniculate nucleus, VMT: Ventromedial thalamus, VP: Ventral posterior thalamus, LHA: Lateral hypothalamic area, PH: Posterior hypothalamic area, ZI: Zona incerta, MLR: Mesencephalic locomotor region, CS: Superior central nucleus raphe, GRN: Gigantocellular reticular nucleus, PG: Pontine gray, PRN: Pontine reticular nuclei, CENT: Cerebellar lobuli, FL: Flocculus

Neural activity was monitored with a fUSi probe positioned over the craniotomy parallel to the long axis of the animal. Sagittal planes were imaged sequentially, each spanning the entire depth of the brain, where the probe was stepped (250 um) along the medial-lateral axis (Figure 2A-C). While imaging each plane, the colliculus was optically stimulated. In each experiment, each plane was imaged for two 20 s periods, when either a 1 s (20 Hz or 50 Hz), or 4 s (5 Hz) light stimulation was delivered via the implanted optic fiber 10 s after the imaging started. The parameters of the light stimulation were the same as those used during the behavioral experiments. Each mouse was imaged in 3-5 sessions that were separated by 48-72 hours. Each voxel was assigned to an individual brain region by performing a 3D rigid registration of the series of sagittal images, obtained in the absence of visual stimulation (125 μm steps), to the Allen Mouse Brain Common Coordinate Framework version 3 (CCF v3) ^41^ (Figure 2D). We used a modified version that is comprised of 264 brain areas in one hemisphere of the brain (Table S2).

To build a spatial map of brain activity, we compared, voxel by voxel, the hemodynamic signals (DI/I, referred to as “activity”) obtained during and after the optogenetic stimulus to a 10 s period before the light stimulus (Figure 2D-F; Movie S8). The hemodynamic activity of all voxels within each area were averaged to estimate the response for that region (Figure 1E and G). Temporal traces were obtained for each mouse and compared (t-test corrected for false discovery rate) to identify the areas that displayed a response (Figure 1G, H-I). Average responses for each segmented area in each mouse line are shown in Figure 1J. The same analysis was applied to low-frequency stimulation data (Figure S3).

### Distribution of temporal response properties

We began our analysis of how distinct cell-types of the SC distribute information across the brain by looking at the temporal structure of the hemodynamic changes induced by the optical stimulation (Figure 3). We found that our 1 s optical stimulus caused a reliable set of temporal responses that could be grouped into 4 broad categories (see Methods). These four response types could be broadly described as: Fast, Delayed, Slow and Inhibitory (Figure 3A and B). The Fast responses were characterized by a relatively fast rise time (1.27 +/- 0.42 s), resulting in a transient response. The Delayed responses showed a clear delay with time to peak of 3.3 +/- 0.79 s. The Slow responding areas started their responses early but took longer to reach their peak (2.1 s +/- 0.70 s) and showed a more sustained response (1.78 +/- 1.30 s). Finally, a set of responses that we will refer to as Inhibitory, showed a negative response. Inhibitory responses were commonly preceded by a very transient early positive response (time-to-peak = 0.91 +/- 0.65 s: decay time = 0.52 +/- 0.18 s) in each cell-class except the GAD2 (Figure 3A and 3B).

**Figure 3.**
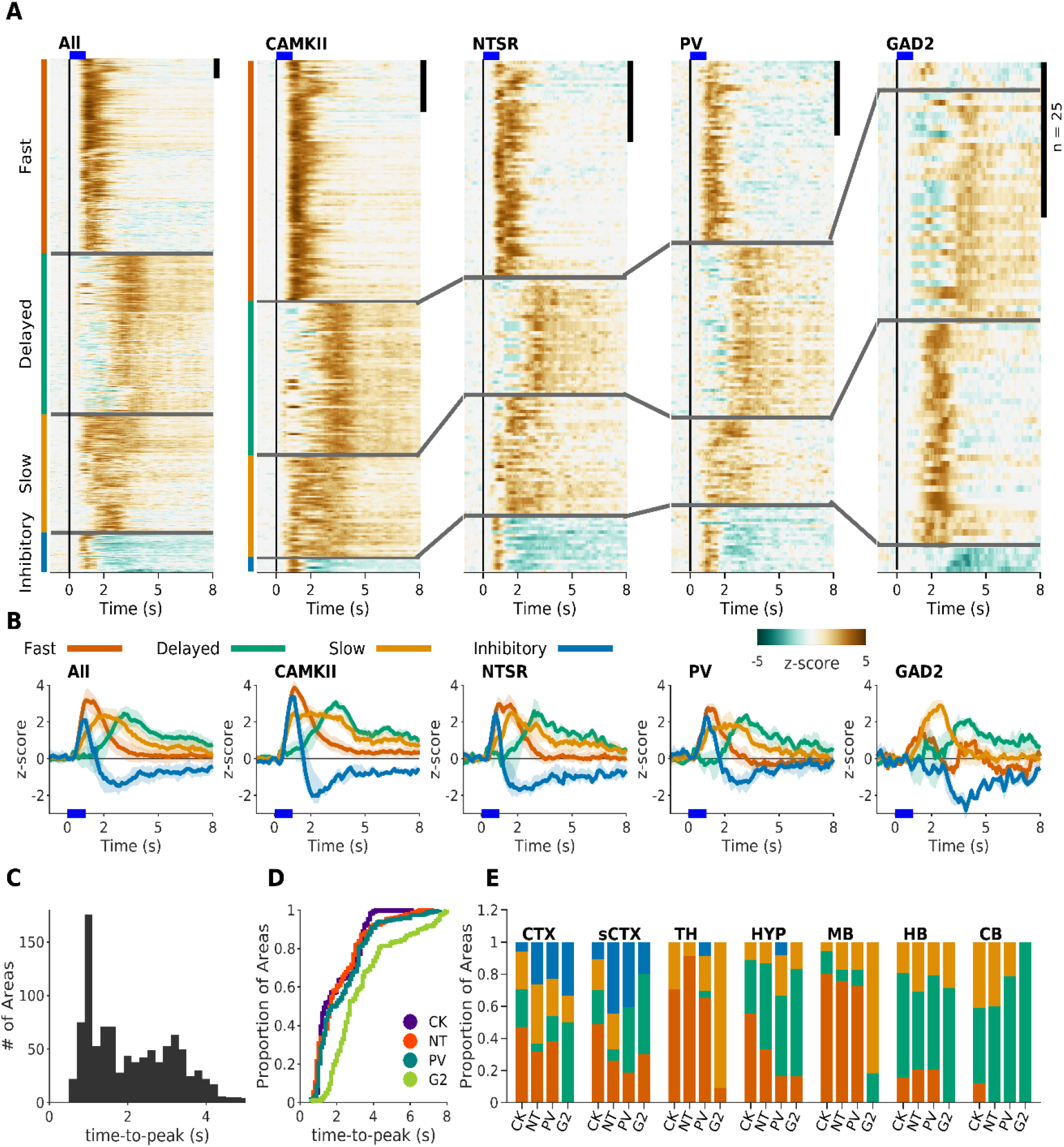
Distribution of temporal responses dynamics. **A.** Normalized responses to optic stimulation. Responses are organized into their respective clusters: Fast (orange), Delayed (green), Slow (yellow) and Inhibitory (blue). Black scale bar at top right of each panel represents 25 areas. Blue line represents the 1 s optical stimulus. Left Panel. Responses of all areas that had a statistically significant response across all cell populations (n = 659). Other panels are the active areas in each mouse line (CK = 246, NT = 157, PV = 170, G2 = 82). **B.** Average response of each of clustered responses. **C.** Histogram of the time-to-peak of each active area in all mouse lines. **D.** Cumulative histogram of time-to-peak in each mouse line. **E.** Proportion of each response type sorted by brain area and mouse line.

The distribution of the different response types varied among the mouse lines. We found that the Fast responses were more common in CAMKII and NTSR mice (CK 47%; NT 42%; PV 35%; G2 5%). PV mice had a similar proportion of fast (35%) and delayed (34%) responses, while GAD2 mice had predominantly delayed (45%) and slow (44%) responses. In addition, inhibitory response types were more common in NTSR and PV mice, as compared to CAMKII and GAD2 (CK 3%; NT 11%; PV 13%; G2 5%). The almost complete absence of Fast responses in the GAD2 mice is evident in the distribution of the response latencies, estimated as the time-to-peak (Figure 3C-D). The distribution of latencies formed two broad groups, those responding within the first 2 s, and those responding after 2 s (Figure 3C). While the distribution of latencies is similar for CAMKII, NTSR and PV cell populations, and spanned the entire range of times, activation of inhibitory GAD2 neurons did not cause any early responses (Figure 3A and 3D).

Stimulating the SC neurons at a lower frequency (5 Hz for 4 s), generated similar temporal dynamics triggered by each neuronal population (fast, delayed, slow, inhibitory), but exacerbated the differences between them (Figure S3). Additionally, the change in frequency corresponded with a reversal in the sign of the response in some areas such as the visual cortex. The visual cortex had a fast positive response during high-frequency stimulus, but an inhibitory response during the low-frequency for each cell population (Figure 3 and Figure S3). The distribution of the different response types across the major brain structures was relatively consistent between the different cell populations, except for slow responses (Figure 3E). Fast responses occurred mainly in the midbrain and thalamus, delayed activations took place in hypothalamus, hindbrain and cerebellum, and inhibitory responses in the cortex and cortical subplate. Slow responses were more homogeneously distributed. For example, CAMKII and NTSR had the largest proportion of slow responding areas in the cerebellum (23 and 24%, respectively), while GAD2 had none in that structure.

### Different brain-wide activity patterns are triggered by each class of cells

To understand how the activation of each neuronal population triggers distinct brain-wide networks, we compared the distribution of brain areas that had increased or decreased hemodynamic responses for each cell-type. Based on the temporal dynamics of the responses observed (Figure 3C), we performed this comparison in two distinct time windows, an early (0-2 s) and a late (3-8 s) phase (Figure 4A). Complete lists of the responsive areas to high- and low-frequency stimuli are provided in the supplementary materials (Table S3-S4). 3D movies of the activated brain areas can be found in the supplementary material for the early (Movie S9-12) and late phase (Movie S13-16).

**Figure 4.**
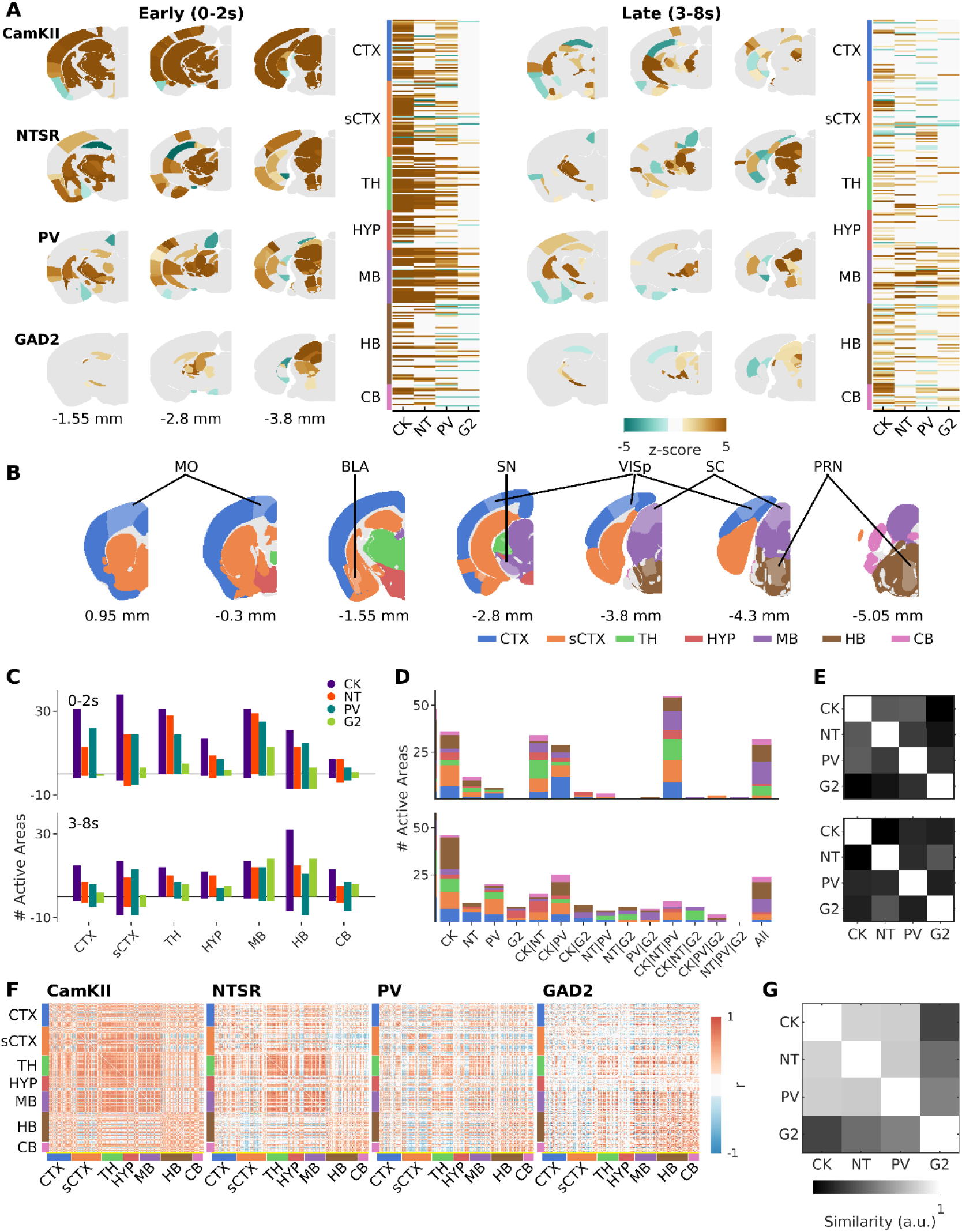
Cell-type specific activation of downstream pathways of the superior colliculus. **A.** Activation maps during early (Left, 0-2 sec) and late (Right, 3-8 sec) time windows. Three example coronal slices are shown for each mouse line. Active areas are shown in each plane with the mean z-score across mice (CK=6; NT=6; PV=13; G2=6). Next to the activation maps, is the peak response of all areas active in at least one mouse line. **B.** Summary of the extent of the imaging locations and corresponding names. Imaging was done in 264 brain regions. Here we delineate the major brain areas and highlight a few single regions for orientation. A complete list of the segmented brain regions and abbreviations are presented in Table S2. **C.** Distribution of active areas during early and late phases. **D.** Quantification of shared areas across mouse lines during early (Top) and late (Bottom) phases. Areas included in one group are excluded from the others. **E.** Similarity matrix (between cell populations) of maximum activity during early (Top) and late (Bottom) phases. **F.** Pairwise Pearson correlation coefficients between the mean response traces of the 264 segmented areas during the 8 s after stimulus onset. **G.** Similarity between the correlated hemodynamic responses in F.

The distribution of responsive areas across the brain in the early and late phases followed different patterns for each of the cell classes (Figure 4A and C). Broadly, stimulating CAMKII, NTSR and PV cell-types at high-frequency, resulted in more responses during the early phase (CK=193; NT=138; PV=129), as compared to the late phase (CK=142; NT=82; PV=97). In contrast, stimulating GAD2 cells resulted in 41 areas responding during the first two seconds, followed by an additional 68 areas responding during the late phase. When we compared the distributions of the active areas (Figure 4C), we found that stimulation of CAMKII evoked responses in large portions of the CTX (80%, 33/41 areas), sCTX (80%, 41/51 areas), TH (80%, 31/36 areas) and MB (89%, 33/37 areas) in the early phase, but was dominated by HB (72%, 39/54 areas) and CB (79%, 15/19 areas) in the late phase. Stimulating the NTSR population activated large portions of the MB (78%), and TH (78%), during the early phase (other structures ranged from 34%-58%) and had less but more distributed activity during the late phase (CTX: 24%, sCTX: 27%, TH: 28%, HY: 38%, MB: 41%, HB: 28%, CB: 42%). PV neurons preferentially modulated the MB (MB: 70%; others: 31-59%) in the early phase, and the HB (HB: 74%; others: 22-43%) in the late phase. In GAD2 mice, most areas were activated in the MB in the early phase (MB: 38%; others: 2-19%). During the late phase, GAD2 activated more areas across the whole brain, particularly in the MB (early/late; 38% / 54%), the HB (early/late; 19% / 33%) and CB (early/late; 16% / 32%). Low-frequency stimulation did not change the overall distribution of responsive areas in early and late phases of the different cell-types (Figure S4A). However, compared to high-frequency, PV and GAD2 mice had a noticeable decrease of activated areas in both the early (high-frequency/low-frequency; PV: 129/72; GAD2: 41/14) and late (high-frequency/low-frequency; PV: 97/73; GAD2: 68/9) phases. Differently, NTSR mice had more responsive areas in both early and late phases, most noticeable in the HB during the late phase (high-frequency/low-frequency; HB: 28% / 93%). Of note, the early phase of NTSR mice had a large increase of negatively modulated areas of the cortex by low-frequency stimulation, which had positive responses upon high-frequency stimulation (Figure S4A-B).

To gain insight into the different downstream networks, we next looked at the overlap between the areas modulated by the different cell classes (Figure 4D-E). We found that in the early phase up to 93 areas had shared activity between at least three of the neuronal populations, 71 where shared by only two, and 54 areas were unique. Consistent with the fact that the CAMKII population likely includes the other two excitatory cell-types, when two areas were shared, in most cases it was between CAMKII and either PV (29) or NTSR (34) mice. During the late phase, the specificity increased and only 47 areas where shared by three or more cell-types, compared to the ones shared by two (70) or uniquely modulated (84). To measure how similar the activated networks were from each other, we calculated the similarity between cell-lines of the maximum activity during early and late phases (Figure 4E). We found that during the early phase, the greatest similarity was between PV and NTSR with CAMKII (Figure 4E). GAD2 showed the least similarity with the other cell lines. During the late phase, GAD2 and NTSR mice showed the highest similarity towards each other (Figure 4E Bottom), and all other pairings showed very low similarity. Low-frequency stimulation was characterized by a large increase in the number of areas solely activated by NTSR neurons (Figure S4E). Similarity analysis showed that, in the early phase, CAMKII and PV networks were the most similar (Figure S4D Top). In the late phase, the similarity pattern was conserved, with GAD2 and NTSR networks being the most similar (Figure S4D Bottom).

Finally, in order to compare the activated networks from a holistic point of view, we generated functional connectivity maps of the relationship between areas across the whole brain (Figure 4F). To do this, we first quantified the pairwise correlation across all active areas of each neuronal population. Then, we compared the resulting matrices to each other (Figure 4G). Broadly, correlations across the brain upon high-frequency stimulation followed similar patterns in CAMKII, NTSR and PV, and were clearly different from GAD2. More concretely, CAMKII, NTSR and PV mice all had a marked high level of correlation between areas of the MB and TH. In GAD2 the highest correlations where between the MB, the HB and the CB. Under low-frequency stimulation, CAMKII and NTSR mice continued to have similar correlation patterns across the brain (Figure S4E-F), but there was a pronounced shift of the brain-wide correlations that showed increased correlations of the MB with the HY, HB and CB and a decrease with CTX and sCTX. The change in frequency did not affect the brain-wide correlations of PV mice, while the correlations for GAD2 became sparser and more localized within the different structures, making it the most differentiated cell-type (Figure S4F). Taken together, these results indicate that each collicular cell-type modulates a distinct brain-wide network.

### Defensive and fear related networks are differentially modulated by each cell class

To understand how the activation of each neuronal population triggers distinct aversive behaviors, we compared the activity patterns of each cell-type within a list of 30 areas that have been previously shown to mediate or modulate defensive behaviors (Figure 5A). This comparison showed that each neuronal population activated a different subset of areas or modulated the same areas in a different manner. For example, the central amygdala (CEA), the posterior medial and paraventricular hypothalamic areas (PMH, PVH), and the ventral tegmental area (VTA) were shared uniquely by cell classes that elicited freezing-like behaviors (CK, NT). On the contrary, the cuneiform (CUN), or the superior central nucleus (CS) were activated by all mouse lines but exhibited different temporal dynamics. The CUN had fast transient responses in CAMKII and NTSR and slower sustained responses in PV and GAD2 mice, whereas the CS had sustained responses in CAMKII and GAD2 but transient in NTSR and PV. Interestingly, there were also cases where different cell-types activated the same areas but in opposite directions. For example, areas of the ventral midline thalamus (RE and Xi), cingulate cortex (ACAd), and subthalamic nucleus (STN), had positive responses to CAMKII and NTSR types, but were dominated by negative responses in PV. Finally, a few areas were similarly activated by all mouse lines, namely the motor layers of the colliculus (SCi), the dorsal periaqueductal gray (PAGd) and the zona incerta (ZI), all with similar fast positive responses. When we compared the correlated activity across this group of areas (Figure 5B), and the similarity of the traces (Figure 5C) across the different mouse lines, it confirmed that CAMKII and NTSR evoked the most similar responses compared to PV, and GAD2. Principal component analysis of the trajectories followed by the responses (Figure 5D), showed that CAMKII and NTSR evoked almost identical responses during the first 2 seconds after the stimulus onset, but then diverged into different paths. This was likely due to the more sustained activity evoked by the excitation of CAMKII neurons (Figure 5E). These results are consistent with the different collicular cell-types activating distinct behaviors through parallel functional networks.

**Figure 5.**
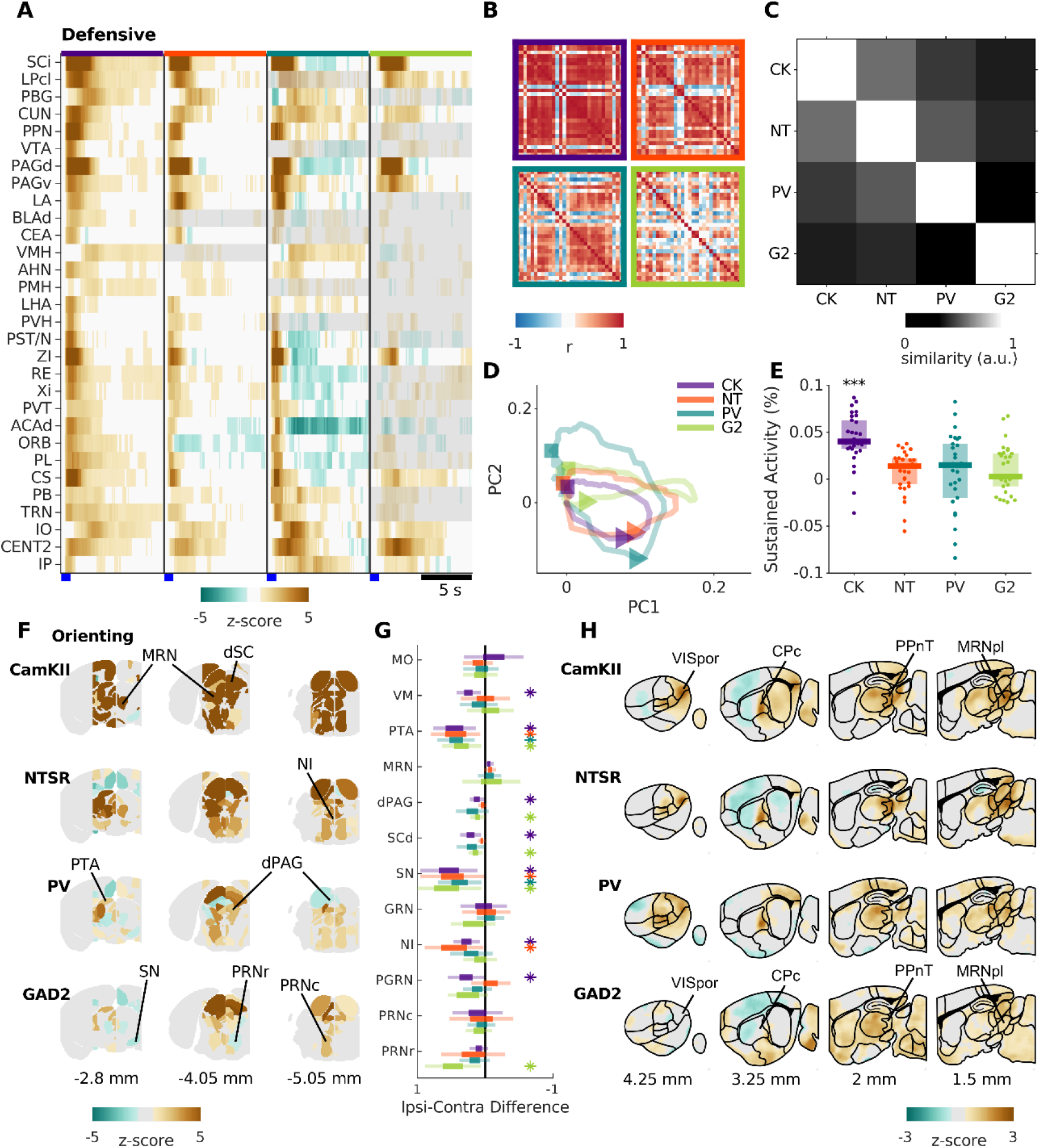
Activity in behaviorally defined networks. **A.** Heatmap of the average responses of 30 nuclei commonly associated with defensive behaviors triggered the different cell populations. **B.** Correlation matrix of the responses in each cell populations. **C.** Similarity of the response properties of these 30 areas across cell populations. **D.** 2D Trajectories of the neuronal activity in these 30 brain areas. Triangle and square are the points on the trajectories 1.5 s and 5 s after the stimulus, respectively. **E.** The average sustained activity for each of the 30 areas after high-frequency stimulus of each cell population. Each dot represents a brain area. **F.** Comparison of the activity in the nuclei that lie within 2 mm of the midline, both ipsi- and contralateral to the optogenetic fiber stimulus. Each ipsi-contra pair is shown when the difference is the highest within a 4 s window after stimulus onset **G.** The percent difference in the fUSi signal between the ipsi- and contralateral brain areas. * represent differences that were statistically significant (p < 0.05 permutation test, after correction for multiple comparisons). Light color lines represent the estimated 95% confidence intervals. Dark portions represent the interquartile range. **H.** Average pixel-pixel maps of four sagittal sections from each neuronal population that highlight brain areas not commonly reported to mediate visually guided defensive behaviors.

### Asymmetric activity originating from the medial part of the superior colliculus

Orienting behaviors, including eye, head and body movements can be controlled via contralateral projections of the SC to the medial pontomedullary reticular formation (MPRF) ^42,43^. In mice the MPRF is comprised of a set of nuclei that includes the pontine gray and pontine reticular nuclei (PRN) that we were able monitor on both sides of the brain as they lie close to the midline. On the contrary, defensive behaviors are thought to be mediated mainly by ipsilateral pathways originating in the medial part of the SC ^44^. Consistent with our stimulations targeting the medial part of the colliculus, we found that all cell-lines evoked asymmetric activations that where preferentially ipsilateral (Figure 5F-G). Also, activating GAD2 neurons generated the greatest number of asymmetries, including pontine areas such as the rostral part of the PRN, which is in line with that cell line being the only one triggering a change in orientation.

### Novel areas involved in collicular-driven aversive behaviors

Visual inspection of brain-wide activity maps (Figure 5H), revealed a few highly responsive nodes in areas that have not been previously studied in the context of collicular-driven defensive behaviors. The four areas that were most salient were the the caudoputamen, especially its caudal part (CPc); the postrhinal visual area (VISpor); the posterior lateral part of the midbrain reticular formation (MRNpl) and a group of thalamic areas surrounding the medial geniculate complex referred to here as the posterior paralaminar nuclei of the thalamus (PPnT) ^45^. The VISpor and CPc are known di-synaptic targets of the colliculus, via the pulvinar, but have not been implicated in guiding defensive behaviors ^46–49^. The MRNpl and PPnT have not been previously described to receive mono- or di-synaptic inputs from retino-recipient neurons of the SC ^14^.

### Correspondence between fUSi and neuronal activity

To test to what degree fUSi signals correlate with the underlying spiking activity, we used Neuropixels probes to record from different parts of the brain and compare them to the fUSi responses observed for the same stimulus. We focused on NTSR cell population in Ntsr-Cre x Chr2 mice. Animals were head-fixed on a treadmill or floating ball and neural activity was recorded while either optogenetically activating NTSR neurons with repeated trials of 1 s 20 Hz stimulation, or while viewing visual stimuli on a screen (Figure 6A). The recording probes were coated with a fluorescent dye (DiI) to visualize the recording locations post-hoc (Figure 6B). On some electrodes, we found spiking activity that was triggered by each of the 20 light pulses (Figure 6C Top). On other electrodes, while the response to the first light pulse was often strong, the responses to the subsequent pulses were weak or absent (Figure 6C middle and Bottom). A raster plot of all 384 recording electrodes for one trial of 20 light pulses are shown for a penetration (from Figure 6B) that passed through the cortex, the SC and periaqueductal gray (Figure 6D). Spikes were defined as local peaks of 4 times the standard deviation of the average activity before stimulation. Detected maxima during the 1 ms pulse itself were excluded since they were contaminated by electrical artifacts (see Methods). Clear responses to the 20 light pulses can be seen on the patch located in the superficial and deep colliculus (approximately electrodes 150 to 250, spanning ~1000 um in depth). We aligned the histological slices including the probe tract with the Allen Brain Atlas, which allowed us to overlay the probe location with the fUSi data of the same coronal slice (Figure 6E). The spiking activity is indicated as a color-coded bar on top of the fUSi data. We averaged and normalized the fUSi data pooled from 6 brains along the Neuropixels probe track and compared it to the spiking activity. In this recording, we found a correlation coefficient between the fUSi and spiking signal of 0.83 (Figure 6E Right). We generally found a stronger correlation between the spiking activity and the fUSi signal in fUSi experiments with stronger responses (Figure 6F; correlation coefficient r = 0.69; n = 26 probe recordings).

**Figure 6.**
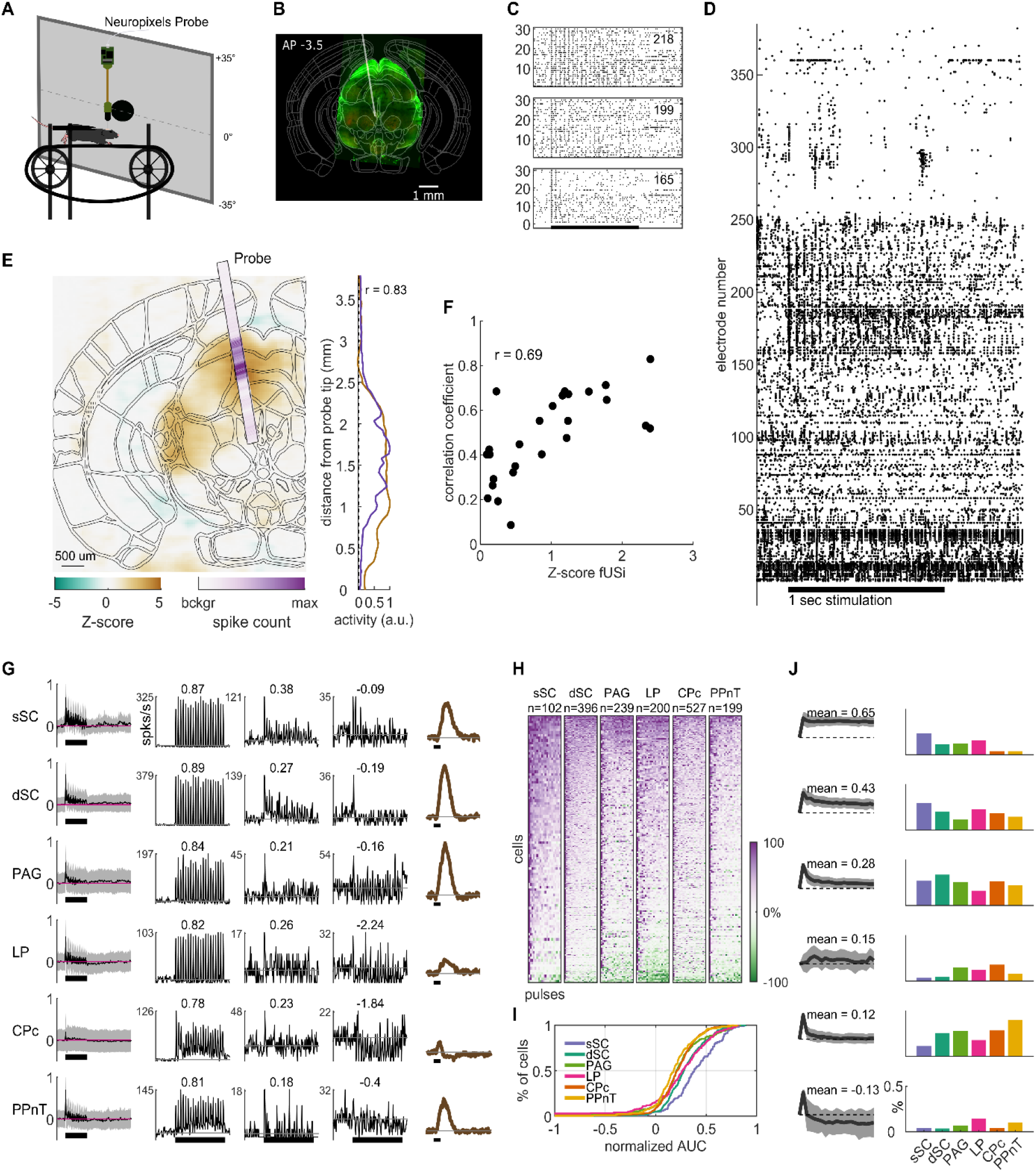
Correspondence of fUSi and spiking activity. **A.** Setup for Neuropixels recordings in awake, head-fixed mice. **B.** Example histological section with Ntsr-Chr2 positive neurons (green) and the probe location indicated with a gray line. **C.** Raster of spikes for 30 repetitions of the optogenetic stimulus for three example electrodes. Stimulus time (1 s, 20 pulses) is indicated with a black bar. **D.** Raw spiking data on all 384 electrodes of the probe shown in B during a 20 Hz optogenetic stimulation of NTSR neurons. **E.** Overlay of fUSi and spiking activity for a colliculus recording. r indicates correlation coefficient of the average activity on the probe and the corresponding pixels of the fUSi data. **F.** Dependence of correlation between fUSi and probe recordings on overall fUSi response strength. **G.** Average response (averaged across all optogenetic stimuli, background-subtracted and normalized) of sorted units in six selected areas (Left; mean ± std). Responses to optogenetic stimulation of 3 example cells for each area. The cell with the highest area under the curve (AUC) (second column), the medium AUC (third column) and the lowest AUC (fourth column) as well as the temporal fUSi response (last column). Numbers indicate AUC. **H.** Response strength to each optogenetic pulse sorted by AUC. 0% is background activity. **I.** Cumulative distribution of AUCs for each area. **J.** Responses to the 20 optogenetic pulses were clustered into 6 types. Average and STD of the normalized response strength for each cluster as well as average AUC (left) and % of cells for each response type and area (Right). sSC - superficial superior colliculus; dSC - deep superior colliculus; PAG - periaqueductal gray; LP - lateral posterior nucleus of the thalamus (pulvinar); CPc - caudate putamen; PPnT - posterior paralaminar nuclei of the thalamus.

### Optogenetic response patterns are different in the colliculus and downstream targets

Next, we asked how the optogenetic activation of neurons in the SC propagates through its direct and indirect output circuit elements. To this end we analyzed optogenetic responses in 6 different brain areas (Figure S5): 1) the superficial SC (sSC), 2) the deep SC (dSC), 3) The periaqueductal gray (PAG) which has previously been linked to aversive behaviors ^50–52^, 4) the pulvinar (LP), which is a direct target of NTSR neurons, 5) The caudal caudoputamen (CPc), and 6) the posterior paralaminar nuclei of the thalamus (PPnT), which have not previously been linked to innate aversive behaviors but were strongly activated during NTSR stimulation (Figure 5H).

In all six areas, we found that the population of single neurons responded well to the optogenetic stimulation (Figure 6G first column). Similar to the raw spiking analysis, each area contained single neurons that responded well to all 20 pulses throughout the 1 s stimulation (Figure 6G second column) as well as cells that responded to the first stimulation only and others that were inhibited by further pulses (Figure 6G fourth column). We found that both the amplitude and the temporal changes in the spiking activity corresponded well to the fUSi signal recorded in the same brain areas (Figure 6G last column). When accounting for the delayed and slower blood response, the temporal profile of the fUSi signal and the probe recordings were similar for each of the tested areas. Areas with a stronger signal in the fUSi experiments showed a corresponding stronger spiking response (Figure 6G), and areas that showed a decrease in blood flow also showed a decrease in firing rate (Figure S6B and S6C).

To quantify the neural responses during the 20 optogenetic pulses, we calculated the mean, background-subtracted response of each responding neuron to the 40 ms after each pulse, normalized these 20 measurements to its maximal response and calculated the area under the curve (AUC) of those 20 values. An AUC of 1 indicates a cell that responds equally well to all 20 pulses, whereas negative AUC values indicate more inhibition than excitation. The resulting activity maps sorted by AUC indicate a different distribution of optogenetic responses in the different areas (Figure 6H; sSC n = 101 units, dSC n = 392, PAG n = 225, LP n = 199, CPc n = 517, PPnT n = 196). We found a higher proportion of sustained responses (high AUC) in the superficial SC, more transient responses in the PPnT, CPc and periaqueductal gray, and a higher percentage of inhibited neurons in the pulvinar (Figure 6I). Subsequently, we then clustered the optogenetic responses into 6 types (Figure 6J). Neurons from the superficial SC were mostly in the more sustained clusters 1-3. Late-onset neurons (cluster 4) were almost absent in the SC but found in the other four multi-synaptic targets. Transient responses (cluster 5) are the dominant response type in the PPnT and inhibition was found in the pulvinar and PPnT (cluster 6). Taken together, these results show that optogenetic stimulation could be traced from the SC across several synapses. Response patterns were different at different stages downstream of NTSR neurons and the temporal profile of area-specific activity measured using spike recordings or fUSi corresponded well with each other.

### Visual responses downstream of NTSR neurons

Activation of pulvinar-projecting neurons has been shown to induce arrest behavior ^7,17^ and we found arrest-like behavior when activating NTSR neurons (Figure 1). In addition, NTSR neurons respond well to visual stimuli mimicking attacking and over-head flying predators ^38^ that induce aversive behaviors ^19,20^. We thus tested whether neurons at different stages in the NTSR output circuitry that respond to optogenetic activation of NTSR neurons would also respond to behaviorally relevant visual stimuli. We found responses to a looming stimulus mimicking an attacking predator in optogenetically activated neurons in all tested brain areas (Figure 7A; sSC: 31 out of 57 optogenetically activated neurons, dSC: 76/139, PAG: 58/60, LP: 25/44, CPc: 59/109, PPnT: 14/50). Neurons showed different response properties including early and late onset responses as well as transient and continuous activity, and inhibition to a looming visual stimulus. These activation patterns were distributed differently in the six tested brain areas (Figure 7B). It is known that stimuli with similar properties as looming, but without ecological relevance, e.g. a dimming stimulus, do not elicit aversive behaviors ^19^. In accordance with these behavioral findings, we could not detect responses to the dimming stimulus in optogenetically activated neurons of the sSC (Figure 7C). Neurons in further downstream areas sometimes responded to dimming stimuli at similar strength as for looming (Figure 7D and 7E). However, in all areas, fewer neurons responded to dimming stimuli as compared to looming stimuli (sSC: 0 out of 6 optogenetically responding cells, dSC: 8/44, PAG: 3/59, LP: 12/44, CPc: 2/7, PPnT: 4/50). We found no dimming responses in the sSC even when including units that did not respond to optogenetic stimulation and similar higher percentages of dimming responses in the LP, CPc, and PPnT (Figure S6). These data show that ecologically relevant visual information is present throughout the multi-synaptic downstream networks of the colliculus that is revealed during opto-fUSi imaging.

**Figure 7.**
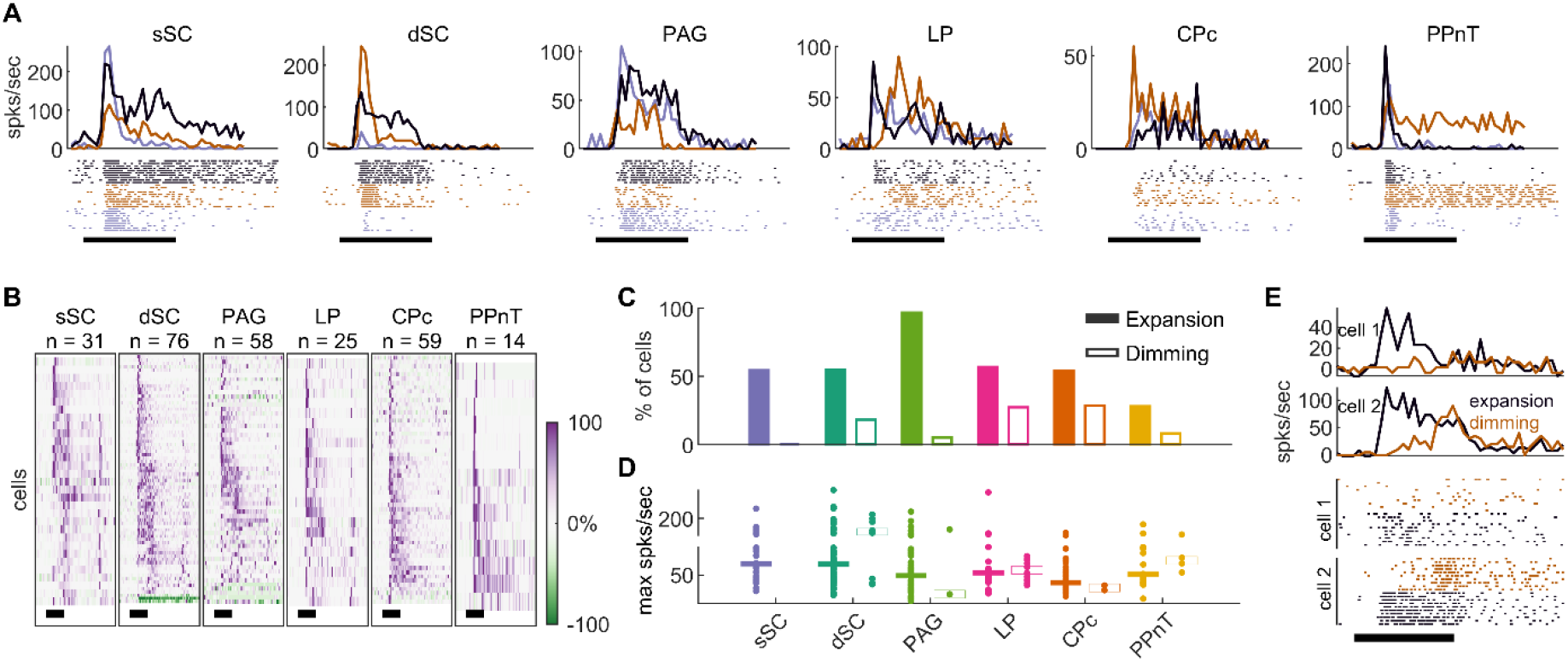
Visual responses in optogenetically activated cells of the NTSR-circuit. **A.** Example responses (raster and spikes/s) to a black looming stimulus for 3 cells with median response strength for each area. Black bar indicates time of looming (0.315 s). **B.** Normalized responses to a black looming disk of all optogenetically activated cells with looming responses in each area. **C.** Percentage of optogenetically activated cells with looming or dimming responses. 100% for looming/dimming for sSC n = 57/6, sSC n = 139/44, PAG n = 60/59, LP n = 44/44, CPc n = 109/7, PPnT n = 50/50. **D.** Maximal response strength to expansion and dimming stimuli for each responding cell and their median per area. **E.** Example of cell with looming, but without dimming response (cell 1), and an example cell responding to both (cell 2).

### Inhibition of PPnT facilitates habituation to repeated stimulation of NTSR neurons

The PPnT has not been previously shown to participate in collicular driven behaviors. Our fUSi data showed that it is consistently activated in response to the stimulation of CAMKII, NTSR and PV neurons of the colliculus (Figure 5). We corroborated that neurons in the PPnT respond to both optogenetic stimulation of NTSR neurons and visual stimuli (Figure 6 and 7). To investigate its role in defensive behaviors, we chemogenetically suppressed activity of its neurons while optogenetically stimulating NTSR neurons in the SC. We injected an AAV coding for the inhibitory DREADDs hM4D(qi) under human synuclein 1 promoter into the PPnT of NTSRxChR2 mice (Figure 8A-B). The same optogenetic stimulation protocol as in previous experiments was used to activate ChR2 in NTSR neurons. We tested mice in the open field arena (Figure 1A) and optogenetically stimulated (20 Hz for 1 s) as the mouse crossed the center of the arena. We conducted five experimental sessions, separated by at least 2 days (Figure 8A). To inhibit the PPnT, clozapine N-oxide (CNO) was injected intra-peritoneally 20-30 minutes before the beginning of the second session.

**Figure 8.**
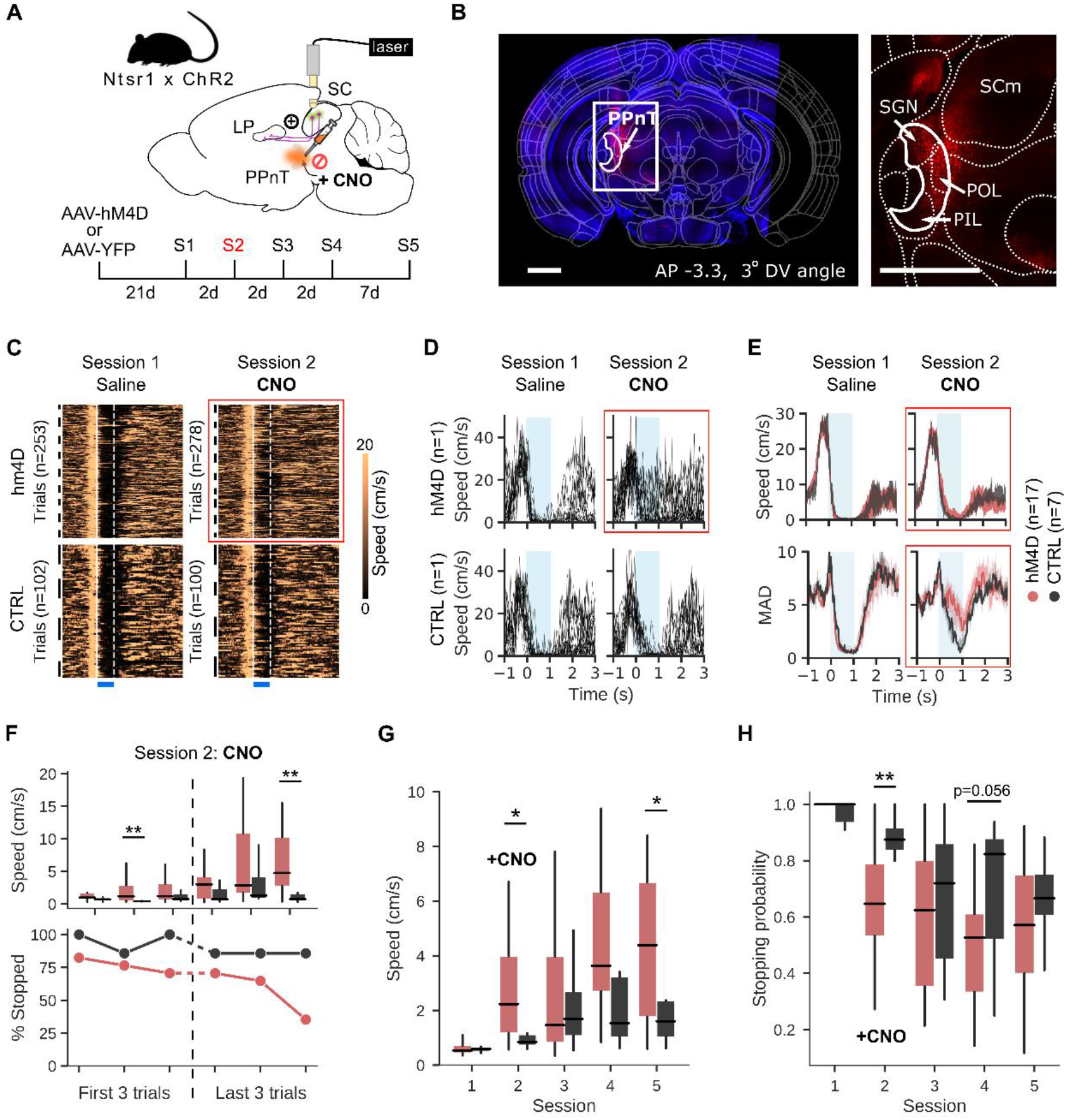
PPnT inhibition facilitates habituation to repeated optogenetic stimulation. **A.** Experimental paradigm **B.** Expression of AAV2-hSyn-hM4D(Gi)-mCherry (red) in the PPnT of a NTSR x Chr2 mouse. Left, coronal brain section aligned to Allen Mouse brain atlas. White rectangle indicates injection site and corresponds to right, zoomed in area. Scale bar 1 mm. **C.** Speed traces of NTSR x Chr2 mice injected with AAV-hM4D (Top row, n=17) and controls (n=7, Bottom row), injected with AAV-YFP for session 1 (saline) and session 2 (CNO). Black lines delineate trials belonging to single animals. White lines delineate onset and offset of optogenetic stimulus. **D.** Speed traces from all trials in session 1 (Left) and 2 (Right) from one example animal from each group. **E.** Top row: median speed traces from all hM4D (red) and CTRL (gray) mice during sessions 1 and 2. Bottom row: mean absolute deviation of the speed for hM4D (red) and CTRL (gray) mice. **F.** Median speed (Top row) and stopping probability (Bottom row) during the one second stimulation of hM4D (red) and CTRL (gray) mice, within the session where CNO was administered. **G.** Median speed of hM4D (red) and CTRL (gray) mice, during the 1 s stimulations across sessions **H.** Stopping probability of hM4D (red) and CTRL (gray) mice, during the 1 s stimulations across sessions. Box plots indicate median, interquartile range, and 5th to 95th percentiles of the distribution. *p < 0.05; **p < 0.01.

We found that inhibition of PPnT increased the variability in the responsiveness of mice to the optical stimulation (Figure 8C-E). This variability manifested as a decrease in the probability that arrest would be triggered by the optogenetic stimulus after a CNO injection (Figure 8F, Movie S17 and Movie S18). During subsequent sessions mice did not regain the lost behavioral response (Figure 8G) and tended to maintain a higher speed than controls during the stimulation periods (Figure 8H).

To investigate the relationship between stopping behavior and the specific location of hM4D expression in the PPnT, we examined the correlation between stopping probability in Session 2 (when CNO was first administered) and the coordinates of the center of expression (Figure S7A-C), or the antero-posterior spread of the expression (Figure S7D). Linear regression revealed weak correlations with the AP, ML, DV planes and extent of expression (Pearson coefficient r=0.112, 0.245, −0.293 and −0.297 respectively). Additionally, we compared the stopping probability of animals with and without DREADD expression in the different areas included in the PPnT (Figure S7E). All the examined mice had expression in the POL. The PoT was present in most animals (n=13/17), but the presence or absence of expression in this area did not change the effect on stopping probability (Mann-Whitney U-test; p=0.198). Interestingly, expression in the PIL and SGN was observed in approximately 50% of the animals (PIL: n=9/17; SGN: n=8/17) and rather than causing the behavioral attenuation, expression in these areas seemed to interfere with the effect, increasing stopping probability (Mann-Whitney U-test; PIL: p=0.037; SGN: p=0.042). We also observed viral expression in the most caudal part of the LP in several animals (n=8/17), but its presence did not correlate with a reduced stopping probability (Mann-Whitney U-test; p=0.168, Figure S8F). Overall, these results suggest that the inhibition of the PPnT, most likely through the POL, facilitates habituation to the repeated activation of collicular NTSR neurons.

## Discussion

In this study, we combined fUSi with optogenetics to reveal the whole-brain neuronal networks that link individual cell-types of the SC with a triggered behavior. We show here that the mouse colliculus distributes information encoded in specific cell-types through distinct networks that share a set of common nodes. Three principals have emerged from this work. First, the spatial and temporal activity patterns evoked by each cell-type are distinct from each other. Second, the observed differences (or similarities) in behavior could not be explained by the activity in any single brain region but appear to be the consequence of distributed activity across many, predominately, subcortical brain areas. Third, while fUSi imaging revealed activation of known downstream targets of each cell-type, it also revealed activity in a set of areas previously not considered as part of these behavioral networks. This allowed us to uncover a putative role of one of these novel targets, the PPnT, in habituation. Direct measurements of neural activity using silicon probes demonstrate a strong correspondence in both space and time between the fUSi signal and mean firing rate in each brain region. Using probe recordings, we also found responses to ecologically relevant visual stimuli in brain regions identified as part of the functional network. Taken together, these results support the notion that in the SC individual cell-types trigger distinct behaviors, not via single dedicated pathways, but instead via distinct brain-wide networks that share a common set of nodes.

### Collicular cell-types activate different, partially overlapping downstream networks

A variety of previous lines of evidence are consistent with our finding that activation of different cell-types of the SC and their output pathways leads to the broad yet restricted propagation of information across the brain. We found that each collicular cell-type relayed information through a different downstream network, that converge in a few key nodes (Figure 4 and 5). Our fUSi experiments show that activation of each cell-type modulated the neural activity of at least 68 and up to 193 brain areas. Among the pathways activated were a set of areas that are consistent with known output pathways of the SC and that have been identified to trigger freezing and escape behaviors ^6,7,10,12–14,50^. Here we demonstrate that activation of the same neural populations results in neural activity in a much larger than expected set of downstream areas. This extensive dissemination of information is likely due in part to recurrent connectivity within the SC ^12,16^, as well as recurrent feedback loops with, for example, deeper layers of the SC, PAG, thalamus and PBG ^14^. While we do observe the previously reported activity in specific nuclei, brain-wide fUSi allowed us to observe functional networks downstream of cell-types in the SC across the most of the brain.

Conversely, the brain-wide activity we observed shows a higher degree of specificity than we would predict from meso-scale maps of area-to-area connectivity ^53^. If, for example, we assume each brain area, like the retinal recipient layers of the SC, projects to at least 6 downstream structures we only need 3 synapses to modulate 216 brain areas. In many cases this underestimates the number of projections a brain area makes. The retina sends projections to approximately 40 targets ^54^, while the primary visual cortex innervates at least 18 cortical and subcortical areas ^55^. We saw our maximal spread of activity when stimulating CAMKII neurons, which modulated 246 areas during the early phase, while NTSR neurons modulated 146 areas. This restriction in the extent of dissemination is likely due to the cell-type specific connection made in the SC and other brain regions. In the SC, distinct output pathways are known to selectively sample retinal inputs and project to selected downstream areas ^13,56^. In addition, outputs of the SC to the LP have been shown to be relayed to a narrow set of downstream targets in the visual cortex and amygdala ^17,48^. This cell-type and pathway specific relay of information is a common feature of several brain structures investigated, including the visual cortex, amygdala and VTA ^55,57,58^.

### Optogenetically triggered behavior can be explained by network activity

How the SC routes the cell-type specific information to evoke different behaviors corresponded well with the observed similarities and differences in the network activity. We observed that activation of CAMKII and NTSR neurons each resulted in an interruption of locomotor activity, which was reflected in the similarity in the brain-wide activity evoked by each cell population (Figure 4). While activating CAMKII and NTSR neurons each interrupted locomotor activity, activating CAMKII neurons resulted in prolonged periods of immobility as compared to NTSR neurons (Figure 1), which corresponded well with the observed differences in the temporal response profiles (Figure 5). For example, in comparison to NTSR, stimulation of CAMKII neurons evoked prolonged activity in areas including the premammillary nucleus of the hypothalamus (PMH) and the superior central nucleus raphe (CS). In addition, we demonstrated that activation of PV and GAD2 neurons facilitated active avoidance and orienting movements, respectively. In comparison to CAMKII and NTSR, activation of PV and GAD2 neurons was characterized by increases in the activity of cerebellar areas like CENT2 and IP. To further discuss how different behaviors are mediated, we focus below on three brain areas: the subthalamic nucleus (STN), the cuneiform nucleus (CUN) and midline thalamus nuclei.

First, activating either CAMKII or NTSR neurons resulted in increase in the activity of the subthalamic nucleus (STN) – a region involved in the interruption of ongoing behaviors ^59,60^. In contrast, the activity of STN was suppressed or not detectable after stimulation of PV and GAD2 cell-types, where locomotion was not interrupted. This suggests that the STN is either activated to pause ongoing behaviors or silenced to promote escape strategies. Second, we found distinct temporal responses in the CUN across mouse lines. The CUN has been shown to trigger freezing and escape ^44^, participate in the initiation and control of locomotion ^61,62^ and modulate cardiovascular response ^63^. We observed fast, transient responses in animals with freezing-like behaviors (CAMKII, NTSR) and slow, more sustained responses, in animals with continuous locomotion (PV, GAD2). It is therefore plausible that different neuronal subpopulations of the CUN are functionally connected with different collicular cell-types to play various roles in defensive behaviors. Third, we demonstrated that the nuclei of ventral midline thalamus (RE and Xi), which play a role in decision making when exposed to threat ^64^, are modulated by both PV and CAMKII neurons but in opposite direction, with sustained activation and inhibition, respectively (Figure 5). Taken together our results suggest that opto-fUSi is a reliable method for studying how the cell-type specific information is disseminated from the SC across the brain to trigger behavior.

### Opto-fUSi reveals new players in aversive behavior driven by the SC

In our experiments, we observed areas consistently activated that have not previously been reported to be involved in mediating visually guided aversive behaviors. Precise circuit dissections have highlighted three major pathways that pass information about visual threat from the SC to downstream areas. These cell-type specific pathways include projections to the amygdala through the LP or the PBG ^67,17^; and projections to the dPAG ^50^. These dissections have led to an atomistic understanding of how the SC mediates aversive behaviors. Here we observed activity in many areas across the brain and four of them captured our attention, namely: the caudal part of caudoputamen (CPc), postrhinal visual area (VISpor), posterior lateral part of the midbrain reticular formation (MRNpl) and PPnT (Figure 5). Some of these areas have been implicated in modulation of visual behavior ^65–68^. The PPnT stood out during visual inspection of activity maps of the brain, as it was both reliably activated upon optogenetic stimulation of NTSR neurons of the SC and during the presentation of visual stimuli. The PPnT is known for its role in associative learning during auditory fear conditioning and in mediating fear discrimination and extinction ^69,70^. Our experiments revealed that this group of areas play a similar role in behaviors triggered by the SC by acting to suppress habituation. Our results are consistent with the proposed role of PPnT in fear extinction and suggest that PPnT is part of the pathway downstream of the SC and is involved in mediating behaviors triggered from the SC. These results highlight the power of combining optogenetic manipulations with brain-wide observation of neuronal activity, which provides a method to identify the brain-wide networks involved and thus the design experiments that allow us to build a more complete picture of how defensive behaviors are controlled.

### Optogenetics and the relationship between fUSi and neural activity

One of the key questions for fUSi imaging is the degree to which it faithfully represents the underlying neural activity. In the context of fMRI, while it is generally accepted that the measured blood oxygen-level dependent (BOLD) signal changes are associated with neuronal activity ^71^; how and where the BOLD signal reliably represents the spiking activity of individual neurons still remains an open question ^72^. Like fMRI, fUSi also relies on the indirect measurement of neural activity through hemodynamic changes, in this case, in cerebral blood volume. Recently, it has been shown that fUS signals, can reliably represent both increases and decreases in local neuronal activity ^28,33^. We provide additional evidence that the changes in blood volume detected by the fUSi are consistent with the local changes in neuronal activity in a number of different brain regions including parts of the cortex, striatum, hippocampus, thalamus and midbrain (Figure 6 and Figure S6).

Light delivery into the brain, similar to our optogenetic activation, has been reported to cause local temperature changes and arterial vasodilation in naïve mice and rats that can cause artefactual signals both in opto-fMRI and opto-fUSi ^73,74^. To minimize such potential effects, in our experiments, we used shorter and lower light intensity stimuli (0.3-0.4 mW, 2ms pulses, 20-50 Hz, 1 s) than the energy threshold calculated (<1 mW, 20 ms pulses, 20 Hz, 2 s) by Rungta et al. 2017. We did not observe any hemodynamic signal in control experiments using naïve mice in these conditions, indicating that the results reported in this study are driven by the activation of collicular cell-types. In addition, we found that the magnitude, sign, and time course of the fUSi signal corresponded well with the local spiking activity measured in the same area of the brain and in response to the same stimuli.

### Brain-wide mapping of function using Opto-fUSi

Understanding the neural basis of defensive behaviors is relevant not only because these behaviors are important for survival, but also because their dysregulation may contribute to anxiety and post-traumatic disorders. To accomplish this, it is necessary to have a holistic understanding of how the nervous system integrates information to guide appropriate behaviors, as well as to know how molecularly defined components of the nervous system contribute to this neuronal activity. Here, we have presented a set of experiments that highlight how combining optogenetics with fUSi can bridge this divide, enabling both the manipulation of targeted components of the nervous system and the simultaneous monitoring of brain-wide activity.

Various methods enable us to look at activity across large parts of the brain, including fMRI, wide-field calcium imaging, large-scale electrophysiology, and genetic markers of neural activity. However, fMRI suffers from low temporal and spatial resolution ^75,76^. Cortical wide calcium imaging and large-scale probe recordings have excellent temporal and spatial resolution but only allow for measurements that are limited to cortex, in the case of calcium imaging, or in thin columns near the electrode tract for silicon probe recordings ^3,77,78^. Finally, genetic markers that putatively report elevated levels of neural activity (e.g. cfos or CaMPARI) provide single cell resolution across the whole brain but have poor temporal resolution and their activation is difficult to interpret ^79,80^.

fUSi enables access to the whole brain at a spatial and temporal resolution (~100 μm^3^, 0.1 s) that allowed us to map the neural network activated by defined cell populations of the SC across a larger portion of the brain. To our knowledge, this is the first time a comprehensive brain-wide mapping has been done at this spatiotemporal resolution of the circuits involved in innate defensive behaviors of mice. Previous experiments in humans and primates have provided evidence of the involvement of structures such as the LC, the PAG, the SC, the visual thalamus, the amygdala, the insular cortex and PFC ^23–25,81–83^. In rodents, molecular (cfos) functional brain maps have implicated a few additional thalamic, hypothalamic and cerebellar areas in various fearful conditions ^64,84–88^. We analyzed the activity of 264 areas across the mouse brain. Our results indicate that the neural pathways involved in mediating and modulating the behavioral responses to activation of the SC stimuli is far more complex that previously reported. We believe combining fUSi with targeted cell-type manipulations and natural stimuli will allow us to understand how different brain regions act in concert to guide defensive behaviors under a variety of conditions.

## ACKNOWLEDGMENTS

This work was supported by the FWO (G094616N to KF, G091719N to KF and AU, MEDI-RESCU2-AKUL/17/049 and 1197818N to AU, 1197818N/1197820N to ASD, 11C5119N/11C5121N to AC and 12S7920N to KR); The Leducq Foundation (15CVD02 to AU); the European Union’s Horizon 2020 research and innovation programme under the Marie Skłodowska-Curie grant agreement No 665501 (12S7917N to KR); A Master Mind Scholarship (F200075 to DL).

## AUTHOR CONTRIBUTIONS

Arnau Sans Dublanc, Conceptualization, Data curation, Formal analysis, Funding acquisition, Validation, Investigation, Visualization, Methodology, Writing—original draft, Writing—review and editing; Anna Chrzanowska, Conceptualization, Data curation, Funding acquisition, Investigation, Visualization, Methodology, Writing—original draft, Writing—review and editing; Katja Reinhard, Conceptualization, Data curation, Supervision, Funding acquisition, Investigation, Visualization, Methodology, Writing—original draft, Writing—review and editing; Dani Lemmon, Data curation, Funding acquisition, Investigation, Visualization; Gabriel Montaldo, Conceptualization, Methodology, Software, Supervision, Writing—review and editing; Alan Urban, Conceptualization, Funding acquisition, Methodology, Software, Supervision, Writing—review and editing; Karl Farrow, Conceptualization, Software, Formal analysis, Supervision, Funding acquisition, Investigation, Visualization, Methodology, Writing— original draft, Project administration, Writing—review and editing.

## Methods

### Animals

All experimental procedures were approved by the Ethical Committee for Animal Experimentation (ECD) of the KU Leuven and followed the European Communities Guidelines on the Care and Use of Laboratory Animals (004–2014/EEC, 240–2013/EEC, 252– 2015/EEC). Male and female adult (2-4 months old) transgenic mice were used in our experiments including, *Ntsr1-GN209Cre, Ai9, Thy1-STOP-YFP, Ai32 x Ntsr1-GN209Cre, Ai32 x PvalbCre and Ai32 x Gad2Cre. Ntsr1-GN209Cre* mice (Genset: 030780-UCD) express Cre recombinase in *Ntsr1-GN209* expressing neurons. *Ai9* (JAX: 007909) and *Thy1-STOP-YFP* (JAX: 005630) are reporter lines that express tdTomato and YFP fluorescent proteins respectively, when in presence of the Cre recombinase. *Ai32* (JAX: 012569) is a reporter line that expresses Channelrhodopsin2 in presence of Cre recombinase. *PvalbCre* mice express Cre recombinase in parvalbumin-expressing neurons. *Gad2Cre* mice express Cre recombinase in *Gad2-*expressing neurons. Mice were kept on a 12:12 h light:dark cycle and sterilized food pellets and water were provided *ad libitum*. Experiments were performed during the light phase.

### General surgical procedures

Anesthesia was induced at the beginning with an intraperitoneal injection of Ketamine (100 mg/kg) and Medetomidine (1 mg/Kg). Before starting any surgical procedure, the paw of the animal was pinched to check for the absence of pedal reflex. After deep anesthesia was achieved, mice were placed in a stereotaxic workstation (Narishige, SR-5N), on a homeothermic blanket to keep a stable body temperature. Eye ointment was applied to protect the eyes from drying and from light (Dura tear, NOVARTIS, 288/28062–7) and Lidocaine (0.5%, 0.007 mg/g body weight) was injected under the skin above the skull. The surgical areas were shaved and the skin was disinfected using iso-betadine. Then, the skin was cut following the midline and retracted to the sides to expose the skull. Anterior-posterior coordinates are measured from Bregma.

### Viral injections

Once the skull was exposed, a hole was performed at the right coordinates by gently rotating a needle against the skull. We used micropipettes (Wiretrol II capillary micropipettes, Drumond Scientific, 5-000-2005) with an open tip of around 30 μm, prepared with a Laser-Based Micropipette Puller (Sutter Instrument, P-2000), and an oil-based hydraulic micromanipulator MO-10 (Narishige) for all stereotactic injections. To trace back the injection sites, we coated the glass pipette tip with DiD (Thermo, D7757).

For optogenetic experiments, we targeted *CamkII-*expressing neurons of the retinorecipient layers of the colliculus by injecting wild type mice with AAV2-CamkII-hChR2(E123T/T159C)-p2A-EYFP-WPRE (UNC vector core, AV5456B). We used *Ai9* and *Thy1-STOP-YFP* mice as wild type mice. We injected 200–300 nl of AAV in 100 nl doses with a waiting time of 5–10 min after each injection. Coordinates for the superficial colliculus were AP: −3.6 to 3.8, ML: −0.2 to −0.3, DV: −1.1 to −1.4. To express ChR2 specifically in *Ntsr1-GN209* expressing neurons of the colliculus, we injected 200–300 nl of AAV2-EF1a-DIO-hChR2(E123T/T159C)-p2A-EYFP-WPRE (UNC vector core, AV5468C) into the superficial colliculus of *Ntsr1-GN209Cre* mice.

For chemogenetic experiments, we injected 300 nl of AAV2-hSyn-hM4D(Gi)-mCherry (Addgene, 50475), or AAV2-hSyn-EYFP (UNC vector core, AV4376E) as control, into the PPnT. PPnT coordinates: AP −3 to −3.4, ML −1.8 to 2, DV: −3.5 to −3.3.

Following injection, the skin was glued with Vetbond tissue adhesive (3M,1469) to close the wound. Next, mice were injected with painkillers (Buprenorphine 0.2 mg/kg I.P.) and antibiotics (Cefazolin 15 mg/kg I.P.) and were allowed to recover on top of a heating pad. After recovery from anesthesia, animals were provided with soft food and water containing antibiotics (emdotrim, ecuphar, BE-V23552) and were monitored for 3 days and administrated Buprenorphine and Cefazolin depending on the condition of the animal. Any following surgery was performed 21 days after injection to allow for proper gene expression.

### Cranial Windows and optic-fiber cannula Implantations

#### opto-fUSi

Once the mouse was anesthetized and the skull was exposed, the lateral and posterior muscles were retracted. Vetbond was applied to open skin and exposed muscle, and a titanium head plate was attached to the skull using dental cement (Superbond C&B, Prestige-dental). Then a cranial window extending over almost the whole extent of the left hemisphere and part of the right hemisphere (AP: +2 to −6.5mm, ML 1.5 to −4.5) was made with a drill. Then, an optic-fiber cannula (Doric Lenses, MF1.25, 200/245-0.37, FLT) was implanted. The entry point of the fiber into the brain was AP: −3.6 to −3.8, ML +1.5 at a 56° angle. The fiber was slowly inserted 1.8 mm into the brain so that the tip would be placed at ML: 0 and DV: −1.1 to −1.2. Next, a ring of dental cement was formed around the craniotomy and the optic-fiber to stabilize the whole preparation. Finally, the cranial window was covered with silicone elastomer for protection and the mouse was allowed to recover on a heating pad. Mice were treated with painkillers (Buprenorphine 0.2 mg/kg I.P.), antibiotics (Cefazolin 15 mg/kg I.P.) and anti-inflammatory (Dexamethasone 0.1 mg/kg) drugs for 5 days.

#### In vivo electrophysiology

Once the mouse was anesthetized and the skull was exposed, Vetbond was applied to open skin and exposed muscle, and a titanium head plate was attached to the skull using dental cement (Superbond C&B, Prestige-dental). Then, cranial windows (~0.5 to 1 mm^2^) were performed over the coordinates of the target regions. The following coordinates were used as the center of craniotomies: SC: AP: −3.7, ML: −0.5; PPnT: AP: −3.3, ML: −1.9; lateral posterior nucleus of the thalamus: AP: −2.1, ML: −1.8; tail of caudate putamen: AP: −1.4, ML: −3.1. An additional hole was made for the implantation of an optic-fiber cannula. The entry point of the fiber into the brain was AP: −3.6 to −3.8, ML: +1.5 at a 56° angle. The fiber was slowly inserted 1.8 mm into the brain so that the tip would be placed at ML: 0 and DV: −1.1 to −1.2.

### Opto-open field test

All behavioral experiments were performed in a custom made square wooden box (W: 50 cm x L: 50 cm x H: 36 cm). Dim ambient light (~50 lux) was provided by a lamp (Paulmann Licht GmbH, PDG09/14) positioned above the arena and oriented away from it, towards a wall. Behavior was recorded at 30 fps using a camera (Point Grey Research, FMVU-03MTM-CS) positioned 53 cm above the center of the arena. For optogenetic activation we used a 473 nm DPSS laser system (Laserglow Technologies, R471003GX) connected to a patchcord with a rotatory joint (Thorlabs, RJPFL2). Optogenetic stimulation was controlled with custom software written in MATLAB. Before every experiment, the output of the laser was measured at 20 Hz or 50 Hz (2 ms pulse width) and set at 0.3-0.4 mW (9.5 – 12.5 mW/mm^2^). For any given mouse line, a high-frequency stimulus was chosen based on preliminary behavioral data. In those mouse lines where 20 Hz stimulation did not evoke any visible response, the following experiments were done at 50 Hz. In the data shown here, CAMKII and NTSR mice were stimulated at 20 Hz whereas PV and GAD2 were stimulated at 50 Hz.

5 days after implantation of the optic fiber, mice were habituated to the handler, patchcord and experimental room for at least 3 days. The day of the test, mice were placed in the center of the arena and were allowed to freely explore for 2 min. After the acclimatization time, when the mice moved away (~10 cm) from the perimeter of the box, towards the center, light stimulation was manually triggered. At any given test, mice where stimulated at high (20 Hz or 50 Hz, 20 or 50 pulses) and low (5 Hz, 20 pulses) frequencies in a pseudo-random manner. Time between stimuli was set to be of at least 30 seconds. A typical experiment lasted 20-40 min.

In tests that combined optogenetics with DREADDs, the experiments where performed as explained above, except that mice were injected with either CNO (2 mg/kg) or saline 30 min prior to the test and where only stimulated at 20Hz.

Repeated tests where always separated by at least 48h.

### Protocol of Functional Ultrasound Imaging

5 days after surgery, mice where habituated to the handler, experimental room and to head-fixation on a platform for 7 days. Then, the awake mouse was head-fixed on the platform and the body movement was partially restrained by a foam shelter. The silicone cap was removed and the cranial window was covered with a 2-3 % agarose layer to reduce brain movement. A 473 nm DPSS laser system was then connected to the optic fiber cannula using a ferrule patch cable (Thorlabs, M83L01). Before every experiment, the output of the laser was measured at 20 Hz or 50 Hz and set at 0.3-0.4 mW (9.5 – 12.5 mW/mm2). Next, acoustic gel (~1 mL, Unigel, Asept) was applied on the agarose for ultrasound coupling and the ultrasound probe (L22-14v, Verasonics) was lowered down to a distance of ~3 mm from the brain. The probe was moved along the lateral axis by a linear microprecision motor (Zaber, X-LRM-DE). At the beginning of each session, a reference anatomical scan was acquired for registration (53 sagittal planes from lateral +1.5 mm to −5 mm, 125 μm steps). Following, we acquired the functional scan (23 sagittal planes, from lateral +1.5 mm to −4.5 mm, 250 μm steps). Two optogenetic stimuli were applied at each plane (high and low frequencies) before moving to the next one. For each stimulus, functional images were acquired for 20 s (10 Hz), and the stimulus was applied after a 10 s baseline. The functional imaging and optogenetic stimulation were controlled and synchronized using custom software written in MATLAB. Optogenetic stimuli consisted of a high (20 Hz, 20 pulses, 2 ms pulse width or 50 Hz, 50 pulses, 2 ms pulse width) and a low (5 Hz, 20 pulses, 2 ms pulse width, ~2 mW/mm^2^) frequency stimulus. The acquisition of the 23 sagittal planes was acquired sequentially starting at lateral −4.5, and the whole craniotomy was imaged 7-12 times per session. Total acquisition time was ~3.5 h.

### Generation of a Functional Ultrasound Image

This procedure was adapted from the sequence for fast, whole-brain functional ultrasound imaging described in Macé et al., 2018. An ultrasound probe containing a linear array of 128 ultrasound emitters/receivers, emitted plane waves (15 MHz, 2 cycles) in five different angles (−6°, −3°, 0°, 3°, 6°). The echoes from each plane wave was acquired with the receivers and adjusted with a time-gain compensation to account for the attenuation of ultrasound signals with depth (exponential amplification of 1 dB/mm). This process generated a single emit-receive image (‘B-mode image’) for each angle and was applied three times for averaging. The 15 individual B-mode images were then combined (~2 ms, 500 Hz), resulting in a higher quality image (’compound B-mode image’).

50 compound B-mode images were acquired every 100 ms (10 Hz) to generate a functional ultrasound image. Blood cells flowing inside the vessels scatter back and shift the frequency of the emitted waves (Doppler effect). Such shifts were measured and extracted in real-time, using singular-value-decomposition-based spatiotemporal filtering, and high-pass temporal filtering (cut-off frequency: 20 Hz). From the filtered data, we calculated the mean intensity of the Doppler signal (Power Doppler) in each voxel. Power Doppler integrates all the Doppler signals in a voxel to obtain an intensity value that is proportional to the amount of blood cells moving in that voxel at a given time. Unlike Color Doppler, it lacks information about velocity or direction of the blood flow but reliably reports hemodynamic changes in blood volume ^27,40,89^. The intensity value of a voxel at a given time was calculated as: *I*(*x, y*) = *A*(*x, y, t*)^2^ where *I* is Power Doppler Intensity, *x*, *y* are the coordinates of a given voxel in a given plane, *A* is the amplitude of the compound B-mode images after filtering, and *t* was time. The resulting functional ultrasound image was 143 x 128 voxels in which each voxel had a size of 52.5 μm x 100 μm x 300 μm ^28^.

### Electrophysiological recordings

12 *NTSR1-GN209-Cre x Chr2* (*Ai32*) mice of either sex at the age of 2.5-3 months were used to record optogenetic and light driven responses in the superior colliculus and PAG (7 recordings), pulvinar (9 recordings), caudateputamen (9 recordings) and posterior paralaminar nuclei of the thalamus (3 recordings).

Two days after performing cranial windows, animals were habituated to the recording set up for 3–4 days. The day of the recording, head-posted animals were fixed on a treadmill or floating ball in front of a screen. Then, a Neuropixels probe phase 3A (Imec, Belgium) ^90^ coated with a fluorescent dye (DiD, Thermofisher) was inserted into the brain with the tip reaching down to 1-1.5 mm below the target area. Once the right depth was reached, it was left to rest for 20-30 min, before starting the recording. Artificial cerebrospinal fluid (150 mM NaCl, 5 mM K, 10 mM D-glucose, 2 mM NaH2PO4, 2.5 mM CaCl2, 1 mM MgCl2, 10 mM HEPES adjusted to pH 7.4 with NaOH) was used to cover the exposed brain and skull.

Neuropixel probes contain 960 recording sites on a single shaft distributed in two rows of 480 electrodes along 9600 μm (16 μm lateral spacing, 20 μm vertical spacing), of which 384 can be recorded simultaneously. In all our experiments, we recorded from the 384 electrodes closest to the tip, spanning 3840 μm. Signals were recorded at 30 KHz using the Neuropixels headstage (Imec), base station (Imec) and a Kintex-7 KC705 FPGA (Xilinx). High-frequencies (>300 Hz) and low-frequencies (<300 Hz) were acquired separately. To select the recording electrodes, adjust gain corrections, observe online recordings, and save data, we used SpikeGLX software. Timings of visual and optogenetic stimulation were recorded simultaneously using digital ports of the base station.

While recording from any given location, first, the superior colliculus of these mice received 30 repetitions of blue light trains (20 Hz, 20 pulses, 1 ms pulse width, 0.2 mW) spaced by 20 seconds intervals. Then, visual stimuli were presented.

### Visual stimuli

Visual stimuli were presented on a 32-inch LCD monitor (Samsung S32E590C, 1920×1080 pixel resolution, 60 Hz refresh rate, average luminance of 2.6 cd/m^2^) positioned perpendicular to the mouse head, at 35 cm from the right eye, so that the screen was covering 90° of azimuth and 70° of altitude of the right visual field. Visual stimuli were presented on a gray background (50% luminance), controlled by Octave (GNU Octave) and Psychtoolbox ^91,92^. The visual stimuli consisted of a small black disc that linearly expanded from 2° to 50° of diameter within 300 ms at the center of the screen and a disk of 50° diameter dimming from background to black within 300 ms. The stimuli were repeated 10 times.

### Immunohistochemistry

Animals were perfused and post-fixed overnight using 4% paraformaldehyde (HistoFix, Roche). Vibratome sections (100-200 μm) were collected in 1x PBS and were incubated in blocking buffer (1x PBS, 0.3% Triton X-100, 10% Donkey serum) at room temperature for 1 hour. Then slices were incubated with primary antibodies in blocking buffer overnight at 4°C. The next day, slices were washed 3 times for 10 min each in 1x PBS with 0.3% TritonX-100 and incubated in secondary antibody solution diluted in blocking buffer overnight at 4°C. We used rabbit anti-GFP (Thermo Fisher, A-11122, 1:500) as a primary antibody to label Chr2-positive cells and anti-mCherry (Novus, NBP2-25158, 1:500) to label hM4D-postive cells. Alexa488 donkey anti-rabbit (Thermo Fisher, A21206, 1:500-1000) and Cy3 donkey anti-chicken (ImmunoJackson, 703-166-155, 1:1000) were used as secondary antibodies. Nuclei were stained with DAPI (Roche, 10236276001, 1:500) together with the secondary antibody solution. Sections were then again washed 3 times for 10 min in 1x PBS with 0.3% TritonX-100 and 1 time in 1x PBS, covered with mounting medium (Dako, C0563) and a glass coverslip. Confocal microscopy was performed on a Zeiss LSM 710 microscope. Images of areas with Chr2- and hMD4-expressing cells, the fiber location and the Neuropixels track labelled with DiD were obtained using a 10x (plan-APOCHROMAT 0.45 NA, Zeiss) objective.

### Analysis of behavioral data

Animal tracking was performed using DeeLabCut software ^93^. Stimulus onsets and offsets were extracted with custom-made Bonsai workflow ^94^. Tracking data were sorted into peri-stimulus trials using custom made Python scripts. Trials where stimulation happened in the periphery of the setup were not included in the analysis, unless explicitly stated. Behavioral parameters were calculated by pooling all trials per mouse and calculating the average, followed by average over mice. Speed was extracted based on positional data of the base of the tail obtained from DeepLabCut. Frames with probability lower than 0.9 were filtered out and linearly interpolated. Position data were smoothed with a median filter of window size = 5. The preferred body angle (Figure 1G) was obtained by first aligning the trajectories to the same initial position and by rotating them to Cartesian X axis by an angel of their body position at the stimulus onset frame. The body position angle is the angle between the line connecting the tail base with the nose and the Cartesian X axis. Next, the preferred angle was calculated between then line of stimulus onset nose position and stimulus offset nose position. Latency to the corner was analyzed as time needed to reach a corner (corner is defined as a square of 10 x 10 cm) after the stimulation onset.

### Analysis of fUSi data

#### Registration

At the beginning of each session, we acquired a reference anatomical map. These anatomical maps were then registered to the Allen Mouse Brain Common Coordinate Framework version 3 (CCFv3) (Allen Brain API; brain-map.org/api/index.html) (Figure 2D). Registration was done semi-automatically based on anatomical landmarks that could be recognized on both the anatomical map and CCFv3 (external outline of the brain, dorsal hippocampus, 3rd and 4th ventricles, cerebellar outline, middle cerebral sinus, colliculus, and corpus callosum). These landmarks were used to readjust the 3D volume of the reference map to the CCFv3 by applying scaling in either of the x, y, z axis and rotations and translations in the coronal, sagittal or axial planes when necessary. Then, we calculated the rotations and translations of the coronal, sagittal and axial planes to create a 3D transformation matrix (from anatomical map to CCFv3).

#### Segmentation

fUSi from each session were automatically registered to the CCFv3 using the 3D transformation matrices obtained in the previous steps. Assignment of voxels to brain areas was based on the CCFv3 segmentation. For our analysis, we excluded fiber tracts, ventricles, unsegmented parts of main brain structures (CTX, CTXsp, TH, HY, MB, HB, CB), merged layers of brain areas, and excluded or merged neighboring areas with volumes < 300 μm^3^. Our final version of the atlas was comprised of 264 brain areas in one hemisphere of the brain (see Table S2).

#### Response Time Traces

The relative hemodynamic changes (ΔI/I) were calculated for each voxel, where I was the baseline (mean of 10 s before stimulus onset) and ΔI was the subtraction of the baseline to the signal at each time point. The traces of the individual areas were obtained for every individual trial by summing all the voxels assigned to each area (Figure 2E).

#### Data filtering and normalization

In order to analyze the response traces of each segmented area, first, we created a dataset with the temporal signals, Ti i=1..Ntime, of each region, trial and session of each animal; T_Animal,repetition_(region,time). In this dataset, all the trials of the different sessions were added. Therefore, the repetitions were Nsessions*Ntrials. The intensity signal obtained from the Power doppler, is susceptible to brain tissue motions caused by the awake animal’s movements. However, motion artifacts can typically be distinguished from hemodynamic changes based on the shape (noise/real; quick spiky/slow curved), and amplitude (noise/real ΔI/I; >100%/1-15%) of the signal. In this study, in order to remove trials affected by motion artifacts we computed the mean temporal signal of each region (Tm) and standard deviation (SD), and then eliminated outlier values where Ti-Tm were 2.5 times higher than the SD. The eliminated values were replaced by the previous non-eliminated value. To eliminate the global variations in the brain (baseline perturbations) we selected the 20% of the regions with lowest response during a 2 seconds time window after the stimulus onset, then averaged these regions to create a baseline signal. This baseline signal was then subtracted from all segmented areas. The resulting normalized temporal traces where used for statistical analysis (Figure 2G).

#### Active Brain Regions

To determine if a brain region was activated by a stimulus, first, we used the normalized temporal traces of every trial to calculate a T-score for each animal, using a general linear model, as commonly used in fMRI ^95^. To take into account the different temporal dynamics present in the responses, the GLM was applied using 1 and 2 seconds stepped (0.5 seconds steps) time windows, starting at the stimulus onset until 7.5 seconds after stimulus. Next, a one-sample two-tailed t test was performed on the n T-scores obtained for the n animals. The region was considered active if the resulting P-value, adjusted for a false discovery rate (Benjamini and Hochberg FDR procedure), was <0.05 (Figure S4D) ^96^. For display (such as in Figure 2J), we quantified the median response time courses across animals, we standardized the responses with regard to the values before stimulus onset (z-score) and we corrected for the relative differences between the stimulation levels of each animal. To do this we calculated a correction factor from normalizing the peak response (A = average of the signal 0.5s around the maximal value within a time window after stimulus onset), across all brain areas where: A_norm(region) = A(region)/sqrt(sum(A(region)^2^), then divided the correction factor across each time point. The Average of all the corrected response traces, for each mouse line, are shown in Figure 2J and S2G and are used in the subsequent figures.

#### Pixel-to-pixel Activity Maps

To visualize pixel-to-pixel activity maps, for each voxel, we quantified the median of the response time courses across trials. We then standardized the responses computing the z-score, using the 10 seconds before stimulus onset and a 2 seconds time window after stimulus onset. Next, we averaged activity maps of animals belonging to the same mouse line applied a median filtering of 4 x 4 pixels on the resulting z-score map. The filtered z-score maps were used for visual inspection of the brain activity and for data visualization (Figure 5H).

#### Clustering of fUSi time courses

Time course of all active brain regions in the four neuronal groups were clustered using hierarchical classification and the e-linkage algorithm, an extensions of Ward’s minimum variance method ^97^. The optimal number of clusters was determined by visual inspection of the mean silhouette value ^98^ and Davies-Bouldin Index ^99^. Clustering was only preformed on the 1056 active areas for the 1s optogenetic stimulus.

### Analysis of Neuropixels recordings

#### Raw spiking activity

To extract spikes from raw Neuropixels data, the average voltage on each electrode in the 0.5 s before onset of the optogenetic stimulation was subtracted from the signal during stimulation. Spikes were identified using the ‘findpeaks’ function in MATLAB with a threshold of 4 standard deviations of the signal before the stimulation. Spikes during the 1 ms of each light pulse were excluded as they could cause artifacts, especially on the electrodes in the superior colliculus.

#### Activity maps

Confocal images of brain slices containing the probe tracks were aligned with the Allen Brain Atlas using the allen CCF tool ^100^. This allowed us to identify the same slice of the fUSi data set. To compare the activity on the probe with the fUSi signal, we averaged the z-scored signal of x-2 to x+2 pixels for each x location of the probe. The raw spiking data on the whole probe was resampled to match the resolution of the fUSi data. We normalized each data set separately to its maximal value, resulting in the plot shown in Figure 6G and analyzed in Figure 6F. Correlation coefficients were calculated using the ‘corrcoef’ function in MATLAB.

#### Spike sorting

The high-pass filtered in-vivo data was automatically sorted into individual units using SpyKING CIRCUS ^101^ with the following parameters: cc_merge = 0.95 (merging if cross-correlation similarity > 0.95), spike_thresh = 6.5 (threshold for spike detection), cut_off = 500 (cut-off frequency for the butterworth filter in Hz). Automated clustering was followed by manual inspection, merging of units if necessary and discarding of noise and multi-units using phy2 (https://github.com/cortex-lab/phy). Units were evaluated based on the average waveform shape and auto-correlogram. Only cells with < 1% of inter-spike intervals of ≤ 1 ms were considered and cross correlograms with nearby neurons were inspected to find spikes from the same neurons ^102^.

#### Detection of responding units

Peri-stimulus histograms (PSTH) were calculated using a bin size of 20 ms. For detection of responding cells and for plotting, the mean spikes/s during 0.3 (looming, dimming) or 0.5 seconds (optogenetic stimulus) before stimulus onset were subtracted from the cell’s activity. We calculated a quality index to capture the reliability of a cell’s response to the 10-30 stimulus repetitions. The quality index was defined as 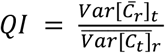 with C being the TxR response matrix, t = time dimension and r = repetition dimension ^103^. Cells were labelled as ‘responding’ if the maximal z-score during the stimulus exceeded 3 and if the quality index was at least 0.15.

#### Transiency measurements

For response transiency when stimulated with 20 light pulses (optogenetics), the mean response during the 40 ms after onset of each pulse was normalized to the absolute maximum of these 20 responses (positive or negative). The transiency of the response was defined as the area under the curve (AUC), i.e. the sum of these 20 values divided by 20. An AUC of 1 means that the cell responded equally well to all 20 pulses, an AUC of −1 means that the cell’s activity was equally suppressed by each pulse.

#### Clustering of optogenetically induced responses

The normalized responses to the 20 light pulses were clustered using k-means (MATLAB) with squared Euclidean distance measurement and 1000 replicates. Based on the Calinski-Harabasz ^104^ and Davies-Bouldin indices ^99^, 7 clusters were chosen where one cluster only contained very few cells with stronger inhibition. These cells were merged with another cluster consisting of inhibited cells, resulting in 6 total clusters.

