## Supplemental Materials for "Brain-wide mapping of neural activity mediating collicular-dependent behaviors"

### The cell-type-specific downstream networks governing collicular behaviors: Supplemental Materials

### Figure S1. Behavior

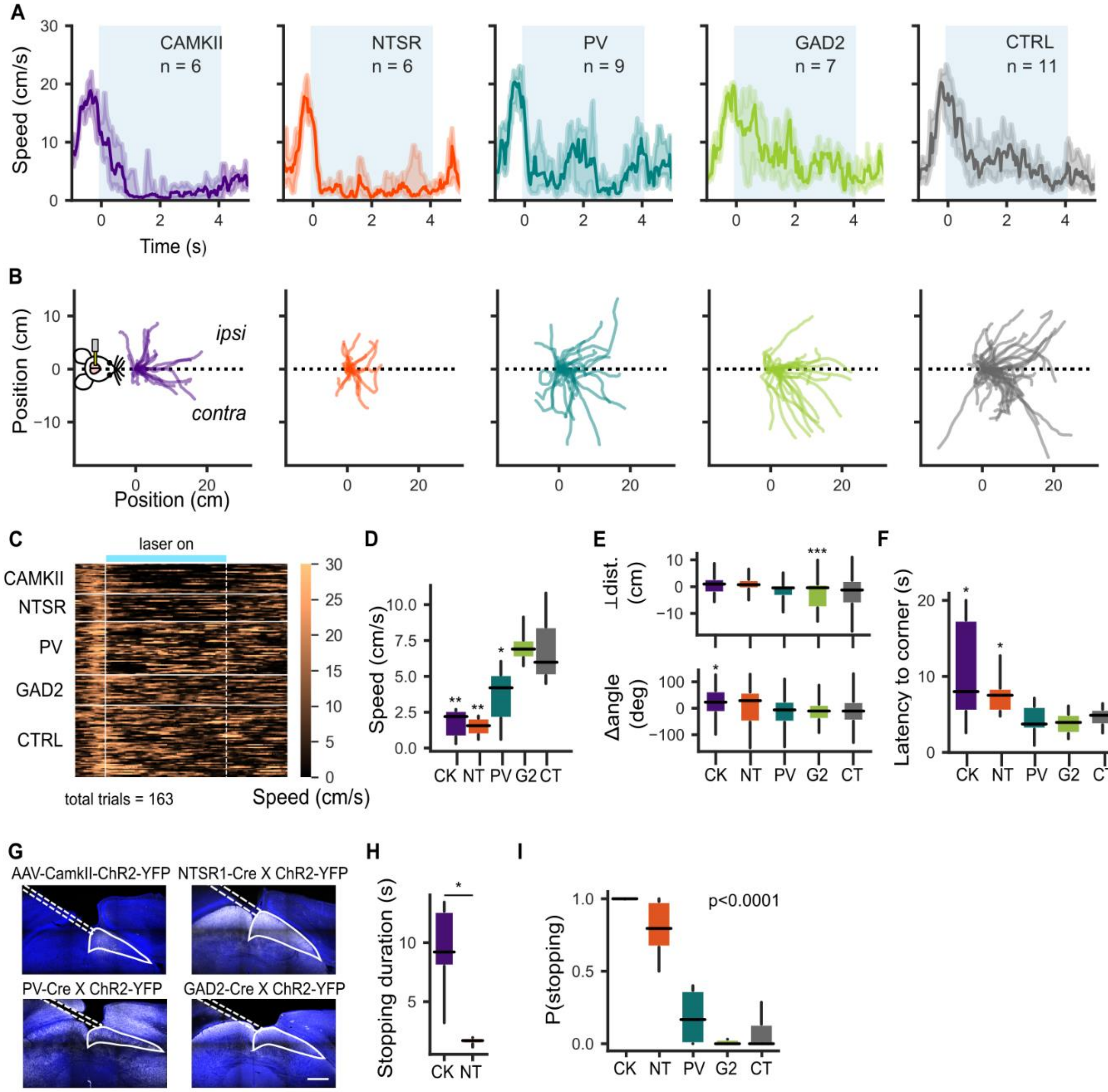


**Figure S1. Behavior. Low frequency (5 Hz) stimulation (A-F) A.** Speed profiles. Each trace represents the median speed obtained from each mouse line. Shaded area represents the interquartile range. **B**. Mice trajectories during the first second of 5 Hz stimulus duration. Traces were aligned and rotated by the initial body position angle. CAMKII: n=6, 23 trials, NTSR: n=6, 21 trials, PV: n=9, 41 trials; GAD2: n=7, 32 trials; CTRL n=11, 46 trials. **C.** Heatmap of mice speeds during optogenetic stimulation trials. Values were obtained from the first experimental session for each animal. Horizontal white lines separate different mouse groups. Vertical solid and dashed white lines mark stimulus onset and offset, respectively. Light blue bar on the top marks the stimulus duration (4s). **D**. Speed quantification during the stimulus duration (CAMKII p=0.009, NTSR p=0.002, PV p=0.02, GAD2 p=0.39). **E**. Quantification of preferred body position at the of the first stimulation second, represented as a change of angle (bottom; CAMKII p=0.02; NTSR p=0.06; PV p=0.49; GAD2 p=0.45) and perpendicular distance (top; CAMKII p=0.27; NTSR1 p=0.29; PV p=0.38; GAD2 p=0.00002), both in reference to X axis (dashed line in B). **F.** Quantification of latency to a corner (CAMKII p=0.03; NTSR p=0.02; PV p=0.5; GAD2 p=0.23). **G.** Coronal section showing expression of ChR2-YFP in distinct cell lines and optic fiber placement. Scale bar: 500 µm. **High frequency (20 or 50 Hz) stimulation (H-I) H**. Stopping duration, p=0.02. **I.** Probability of stopping (CAMKII p=0.0003; NTSR p=0.0003; PV p=0.00001; GAD2 p=0.00008). All data points are averaged over mice, except in E where the data points are averaged over trials.

Significance between control and each mouse line was tested using Mann-Whitney U test (alpha=0.05). Box-and-whisker plots for D-F, H, I show median, interquartile range and range. * P <0.05, ** P<0.01, *** P<0.001.

### Figure S2. fUSI Methods

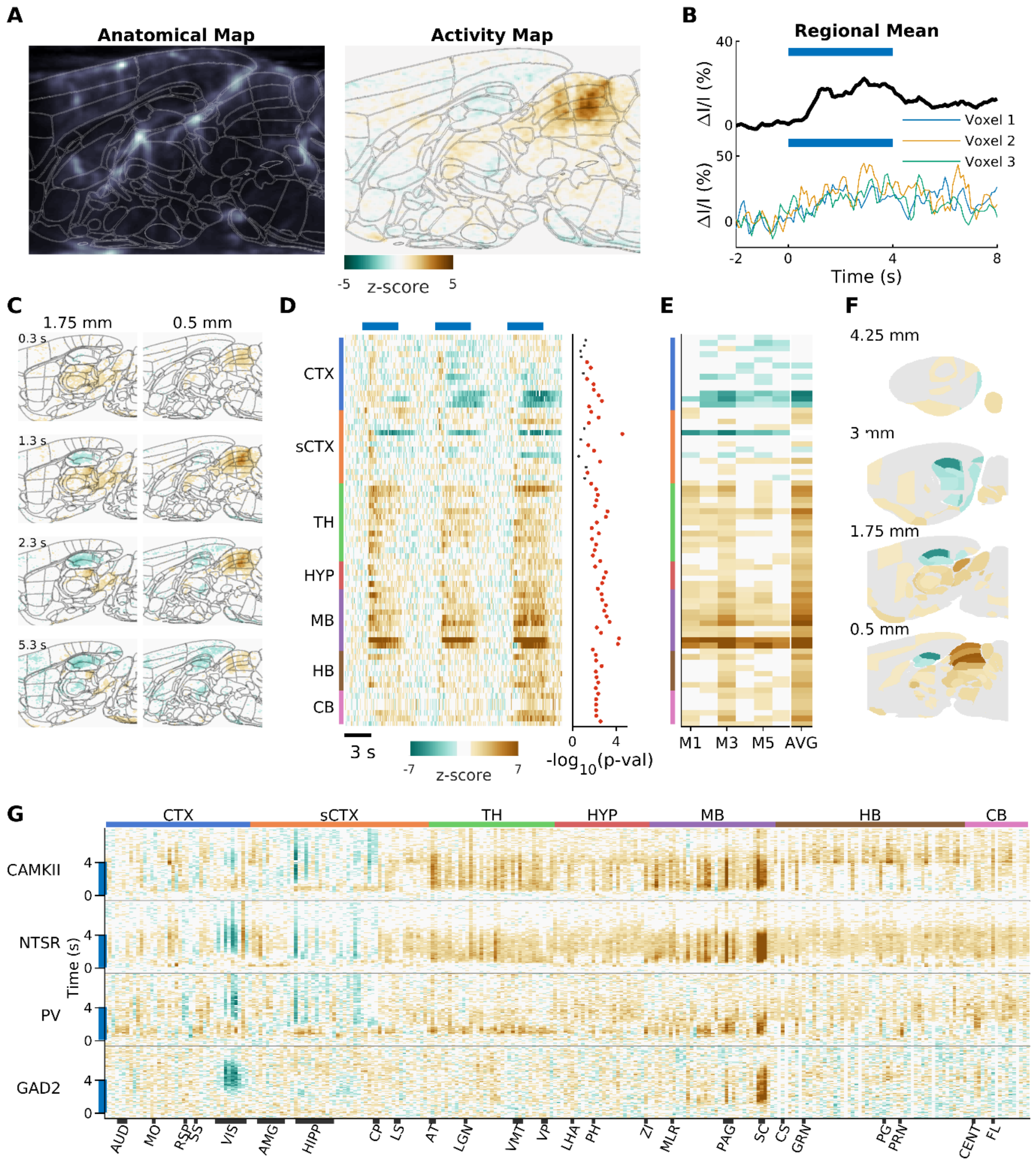


**Figure S2. Functional ultrasound imaging of awake mice during optogenetic stimulation at 5 Hz. A.** Left. Example sagittal section of a blood volume map registered to the Allen Mouse brain reference atlas (thin gray lines). Right. Voxel to voxel normalized response to optogenetic stimulation of plane shown in left panel, registered to the Allen Mouse brain reference atlas (thin gray lines). **B.** Bottom: relative hemodynamic response curves to the optogenetic stimulation of three example voxels in the intermediate superior colliculus. Top: mean response of the intermediate superior colliculus. Blue lines indicate duration of optogenetic stimulation. **C.** Two example sagittal planes from the activity maps of a single animal. **D.** Left: Standardized responses of a selection of 72/264 segmented areas. Mean responses are shown for 3 different mice. Response for each mouse is an average of 6 trials. Blue lines indicate duration of optogenetic stimulation. black thick line indicates optogenetic stimulation. Right: Inactive (gray)

and active (red) areas colored based on significance threshold corrected for multiple comparisons ($p<.05$). **E.** Mean response of each segmented area shown in D during the 2 s after the start of the stimulus for 6 different NTSR mice and the average across all mice. Areas considered not significant ($p > .05$) are set to zero in the average. **F.** Projection of the average activity vector from H onto a map of the mouse brain. **G.** Average time course of each of the 264 segmented areas for each stimulated cell population. Black bars along the bottom indicate span of the labeled brain regions.

CTX: cortex, sCTX: cortical subplate, TH: thalamus, HYP: hypothalamus, MB: midbrain, HB: hindbrain, CB: cerebellum

AUD: Auditory cortex, MO: Motor cortex, RSP: Retrosplenial cortex, SS: Somatosensory cortex, VIS: Visual cortex, AMG: Amygdalar complex, HIPP: Hippocampus, CP: Caudatoputamen, LS: Lateral septum, AT: Anterior thalamus, LGN: Lateral geniculate nucleus, VMT: Ventromedial thalamus, VP: Ventral posterior thalamus, LHA: Lateral hypothalamic area, PH: Posterior hypothalamic area, ZI: Zona incerta, MLR: Mesencephalic locomotor region, CS: Superior central nucleus raphe, GRN: Gigantocellular reticular nucleus, PG: Pontine gray, PRN: Pontine reticular nuclei, CENT: Cerebellar lobuli, FL: Flocculus

#### Figure S3. fUSI Temporal Responses

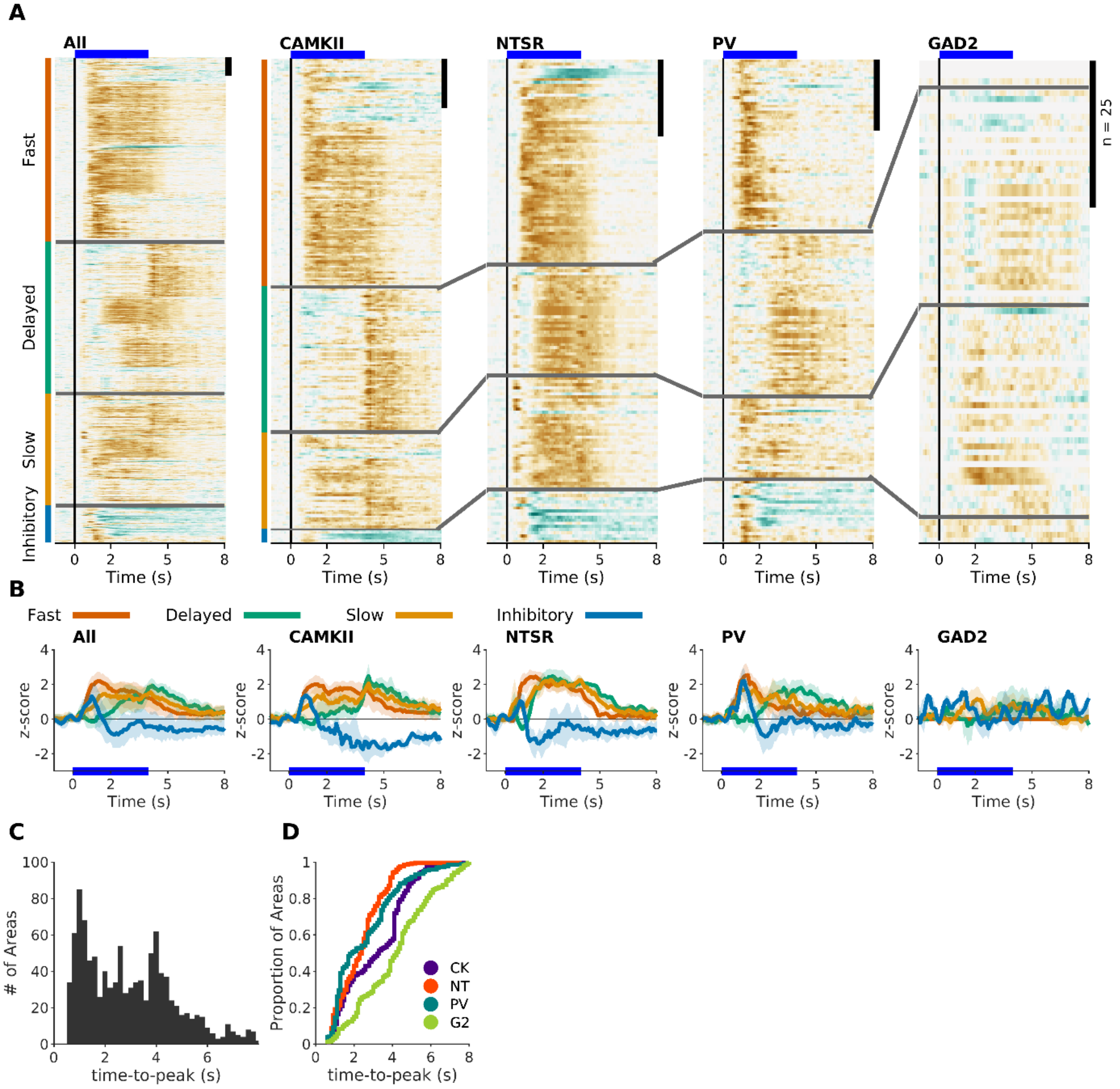


**Figure 3. Distribution of temporal responses dynamics (5 Hz). A.** Normalized responses to optic stimulation. Responses are organized into their respective clusters: Fast (orange), Delayed (green), Slow (yellow) and Inhibitory (blue). Black scale bar at top right of each panel represents 25 areas. Blue line represents the 1 s optical stimulus. Left Panel. Responses of all areas that had a statistically significant response across all cell populations (n = 659). Other panels are the active areas in each mouse line (CK = 246, NT = 157, PV = 170, G2 = 82). **B.** Average response of each of clustered responses. **C.** Histogram of the time to peak of each active area in all mouse lines. **D.** Cumulative histogram of type to peak in each mouse line.

#### Figure S4. fUSI Holistic Activity

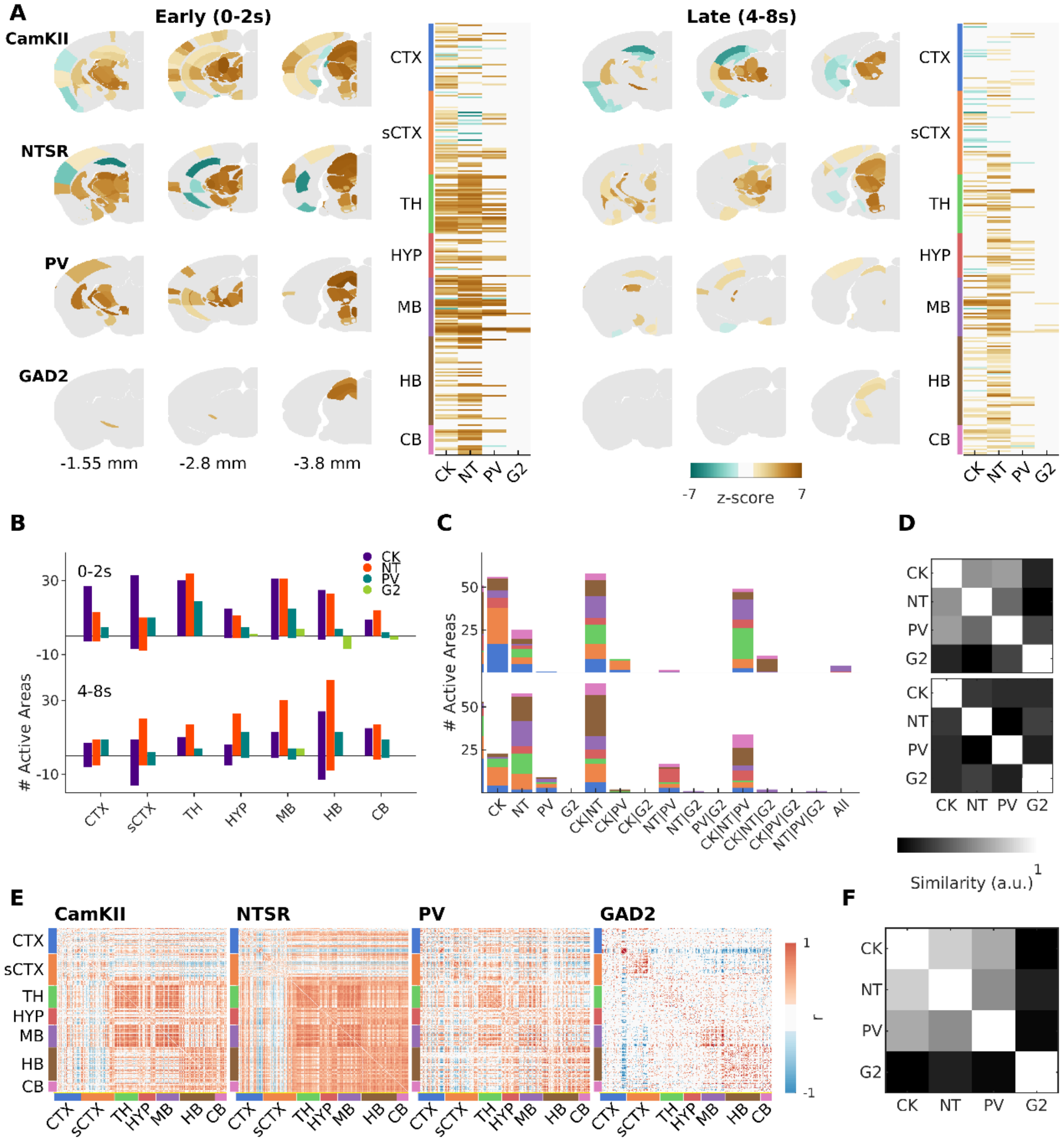


**Figure S4. Cell-type specific activation of downstream pathways of the superior colliculus (5 Hz). A.** Activation maps during early (left, 0-2 sec) and late (right, 4-8 sec) time windows. Three example coronal slices are shown for each mouse line. Active areas are shown in each plane with the mean z-score across mice (CK=2; NT=6; PV=13; G2=6). Next to the activation maps, is the peak response of all areas active in at least one mouse line. **B.** Distribution of active areas during early and late phases. **C.** Quantification of shared areas across mouse lines during early (top) and late (bottom) phases. Areas included in one group are excluded from the others. **D.** Similarity matrix (between cell population) of maximum activity during early (top) and late (bottom) phases. **E.** Pairwise Pearson correlation coefficients between the mean response traces of the 264 segmented areas during the 8 s after stimulus onset. **F.** Similarity between the correlated hemodynamic responses in E.

### Figure S5. Neuropixel Recordings

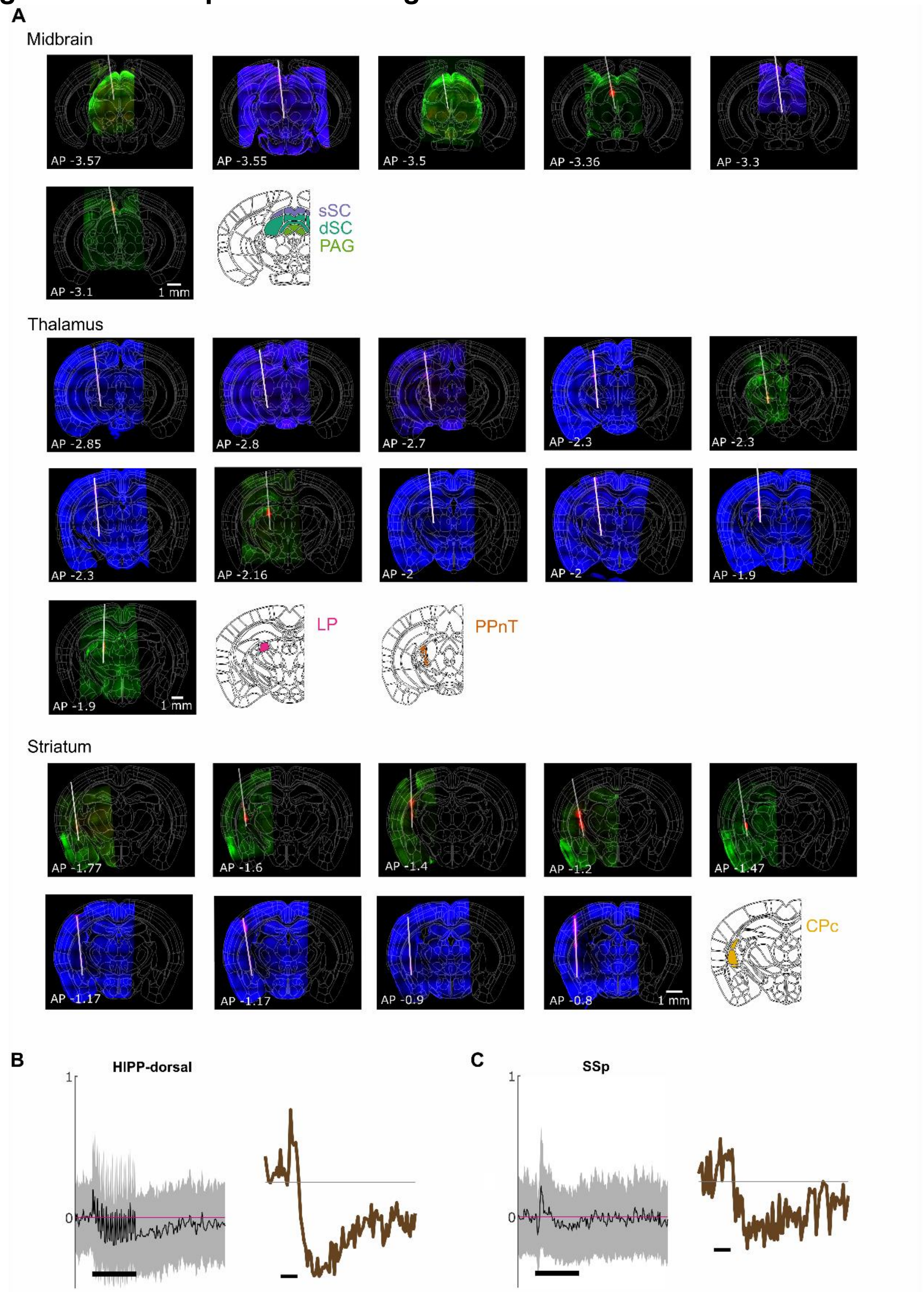


**Figure S5. Probe tracts and temporal response profile of Neuropixel recordings in Ntsr+ mice. A.** One histological slice per recording is shown. The slices were stained for cell nuclei (DAPI, blue) or Chr2-YFP (green). The probe was coated in DiD before recordings (magenta). Probe tracts are represented with gray lines. For each location, an Allen Brain Atlas coronal slice is shown and the analyzed brain areas are highlighted. **B.** Mean ± std of all sorted units (spiking activity) in the dorsal hippocampus recorded during Neuropixels probe recordings (left). Corresponding fUSi signal of the same area (right). **C.** Average spiking activity and corresponding fUSi signal for the somatosensory cortex.

sSC: superficial superior colliculus, dSC: deep superior colliculus, PAG: periaqueductal gray, LP - lateral posterior nucleus of the thalamus (pulvinar); CPc - caudate putamen, caudal part; PPnT - posterior paralaminar nuclei of the thalamus; HIPP-dorsal – dorsal hippocampus; SSp – somatosensory cortex

**Figure S6.**

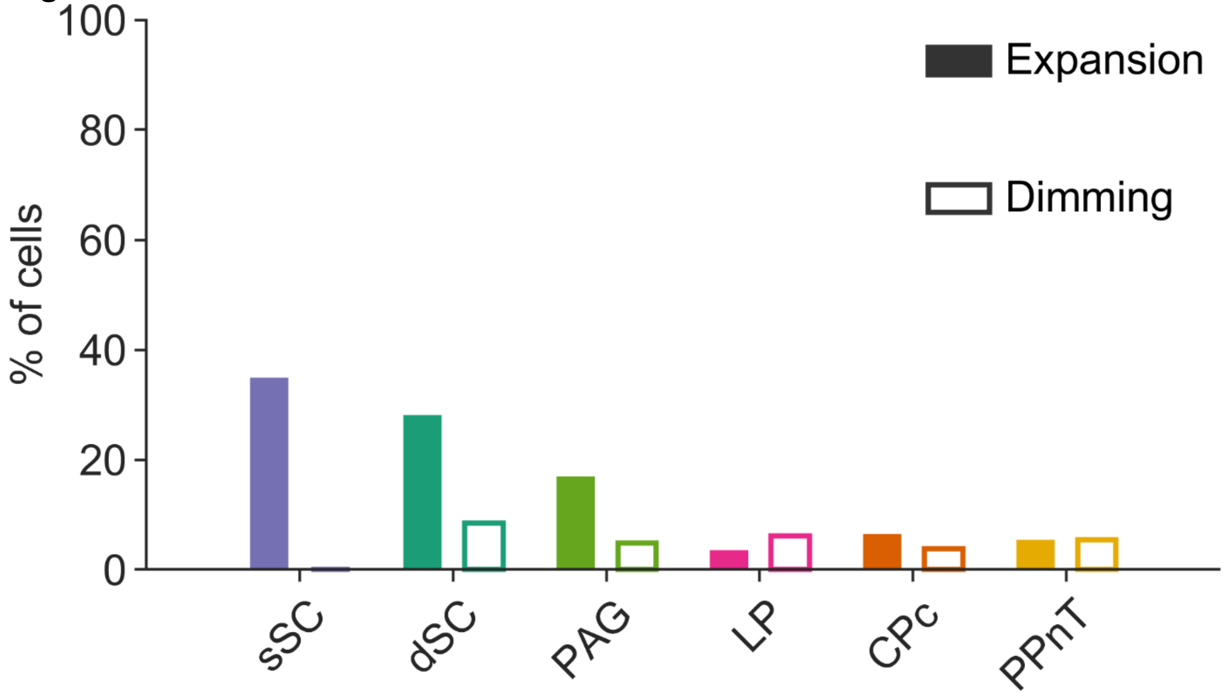


**Figure S6: Responses to expansion and dimming of sorted neurons.** Relates to Figure 7. Instead of only considering optogenetically activated neurons, visual responses of all recorded and sorted units in six brain areas are included. sSC 51/146 (expansion) | 0/21 (dimming), dSC 160/569 | 11/130, PAG 82/483 | 8/163, LP 12/338 | 16/263, CPc 136/2098 | 47/1255, PPnT 16/294, 16/294.

### Figure S7. Chemogenetic Inhibition

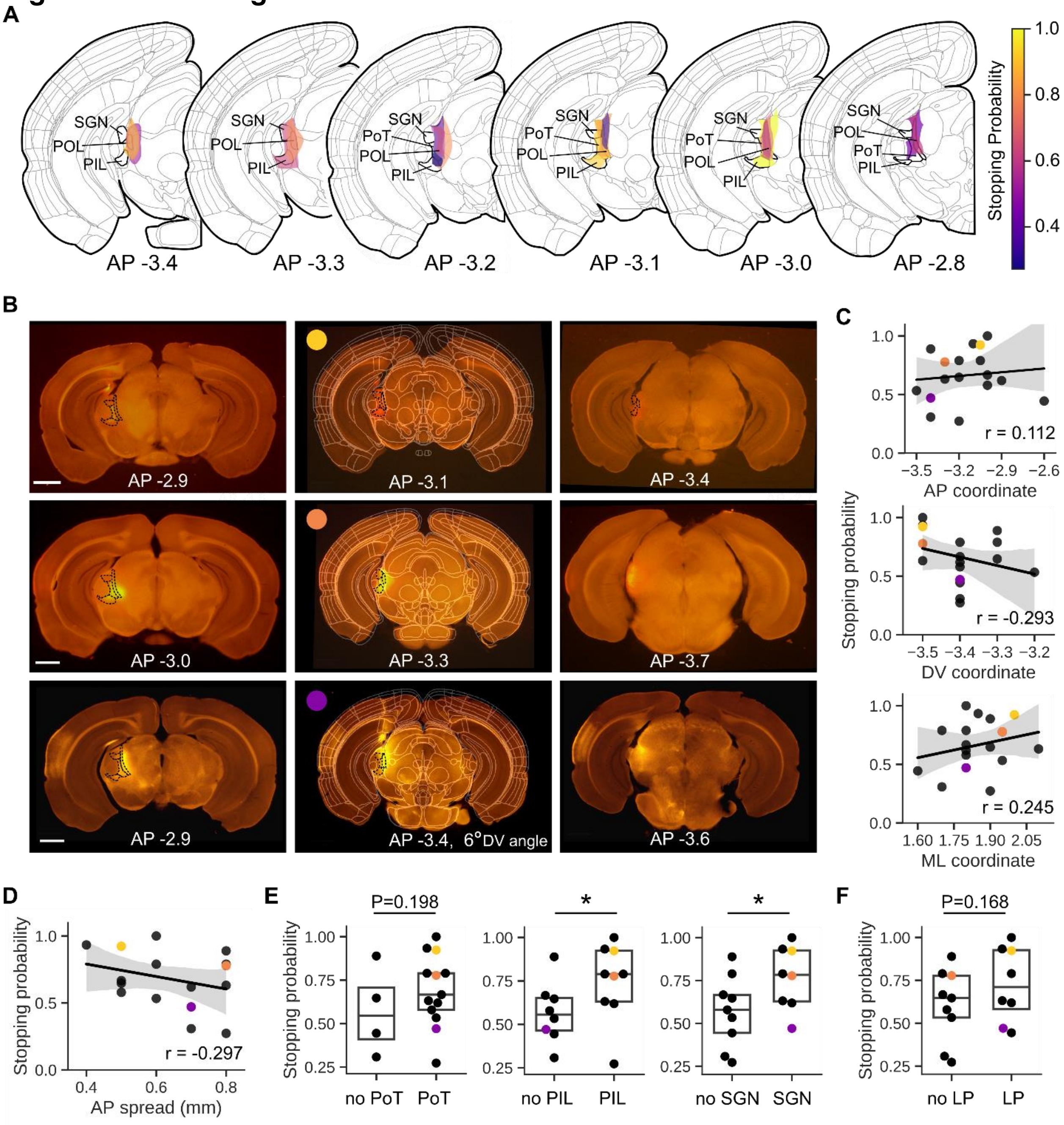


**Figure S7: Histological analysis of the AAV-hM4D expression in the PPnT. A.** Schematic representation of the centers of AAV-hM4D expression for all animals (n=17) grouped on the AP plane. Color represents stopping probability in the CNO-administered. PPnT areas are outlined and labeled in black. **B.** Coronal slices of three example injections aligned to the reference atlas (CCFv3). Each row contains images from a single animal, and in each column is an image of the anterior limit of hM4D expression (left), center of expression (center), and posterior limit of expression (right). The PPnT areas are outlined by dashed black lines. Colored circle in the center column indicates stopping probability of the associated animal based on the colormap in Panel A. Scale = 1mm. **C.** Relationship of Session 2 (CNO) stopping probability of AAV-hM4D mice to injection center coordinates on the AP (top), DV (center), and ML (bottom) planes (n=17). Black line is line of best-fit from linear regression, and shaded area represents the 95% confidence interval. **D.** Relationship of Session 2 (CNO) stopping probability of AAV-hM4D mice to anteroposterior DREADD expression spread **E.** Stopping probability of AAV-hM4D mice with and without expression in the PoT (left), PIL (center), and SGN (right). **F.** Stopping probability of AAV-hM4D mice with and without expression in the LP. Box plots indicate median and IQR. Statistical analysis used Mann-Whitney U test (* = $P < 0.05$). **C-F.** Colored points correspond to histology examples in Panel B.
