## Supplemental Tables S1-S4 for "Brain-wide mapping of neural activity mediating collicular-dependent behaviors"

#### Table S1 | Description of movies

| Movie | Description |
|---|---|
| Related to Figure 1 | |
| Movie S1 | Typical exploratory behavior of a mouse before first optogenetic stimulation trial. |
| Movie S2 | Open field behavioral experiment of a control mouse. Blue light stimulation (20 or 50 Hz for 1 s) does not result in any identifiable behavior. |
| Movie S3 | Open field behavioral experiment of a mouse expressing ChR2 in CAMKII neurons. Blue light stimulation (20 or 50 Hz for 1 s) results in defensive behavior, manifested as long stopping after the stimulation onset. |
| Movie S4 | Open field behavioral experiment of a mouse expressing ChR2 in NTSR neurons. Blue light stimulation (20 or 50 Hz for 1 s) results in defensive behavior, manifested as short stopping after the stimulation onset. Locomotor activity is resumed shortly after the stimulation offset. |
| Movie S5 | Open field behavioral experiment of a mouse expressing ChR2 in PV neurons. Blue light stimulation (20 or 50 Hz for 1 s) results in defensive behavior, manifested as short arrest followed by the movement towards the corner. |
| Movie S6 | Open field behavioral experiment of a mouse expressing ChR2 in GAD2 neurons. Blue light stimulation (20 or 50 Hz for 1 s) results in orienting behavior, manifested as contralateral turning. |
| Movie S7 | Open field behavioral experiment of a mouse expressing ChR2 in GAD2 neurons. Blue light stimulation (20 or 50 Hz for 1 s) results in orienting behavior, manifested as contralateral whole-body drift. |
| Related to Figure 2 | |
| Movie S8 | Spatial map of a brain activity (the hemodynamic signals) obtained during and after the optogenetic stimulation of each cell-type. |
| Related to Figure 4 | |
| Movie S9 | 3D movie of all responsive brain areas (first part) and defensive circuit elements only (second part) during the early phase (0-2 s) upon CAMKII neurons stimulation. |
| Movie S10 | 3D movie of all responsive brain areas (first part) and defensive circuit elements only (second part) during the early phase (0-2 s) upon NTSR neurons stimulation. |
| Movie S11 | 3D movie of all responsive brain areas (first part) and defensive circuit elements only (second part) during the early phase (0-2 s) upon PV neurons stimulation. |
| Movie S12 | 3D movie of all responsive brain areas (first part) and defensive circuit elements only (second part) during the early phase (0-2 s) upon GAD2 neurons stimulation. |

| | |
|---|---|
| Movie S13 | 3D movie of all responsive brain areas (first part) and defensive circuit elements only (second part) during the late phase (3-8 s) upon CAMKII neurons stimulation. |
| Movie S14 | 3D movie of all responsive brain areas (first part) and defensive circuit elements only (second part) during the late phase (3-8 s) upon NTSR neurons stimulation. |
| Movie S15 | 3D movie of all responsive brain areas (first part) and defensive circuit elements only (second part) during the late phase (3-8 s) upon PV neurons stimulation. |
| Movie S16 | 3D movie of all responsive brain areas (first part) and defensive circuit elements only (second part) during the late phase (3-8 s) upon GAD2 neurons stimulation. |

Related to Figure 8

| | |
|---|---|
| Movie S17 | Inhibition of PPnT neurons facilitates habituation to repeated optogenetic stimulation of collicular NTSR neurons. |
| Movie S18 | Inhibition of PPnT neurons – control experiment. |

### Table S2 | Brain regions

| ABBREVIATION | FULL NAME |
|---|---|
| ACAd | Anterior cingulate area, dorsal part |
| ACAv | Anterior cingulate area, ventral part |
| AIa | Agranular insular area |
| AUDd | Dorsal auditory area |
| AUDp | Primary auditory area |
| AUDpo | Posterior auditory area |
| AUDv | Ventral auditory area |
| AON | Anterior olfactory nucleus |
| APr | Area prostriata |
| DP | Dorsal peduncular area |
| ECT | Ectorhinal area |
| GU | Gustatory areas |
| ILA | Infralimbic area |
| MOp | Primary motor area |
| MOs | Secondary motor area |
| NLOT | Nucleus of the lateral olfactory tract |
| OLF | Olfactory areas |
| ORB | Orbital area |
| OT | Olfactory tubercle |
| PERI | Perirhinal area |
| PIR | Piriform area |
| PL | Prelimbic area |
| RSPagl | Restrosplenial area, lateral agranular part |
| RSPd | Retrosplenial area, dorsal part |
| RSPv | Retrosplenial area, ventral part |
| SSp | Primary somatosensory area |
| SSs | Supplemental somatosensory area |
| TEa | Temporal association areas |
| TR | Postpiriform transition area |
| TT | Taenia tecta |
| VISC | Visceral area |
| VISa | Anterior visual area |
| VISal | Anterolateral visual area |
| VISam | Anteromedial visual area |
| VISl | Lateral visual area |
| VISli | Laterointermediate visual area |
| VISp | Primary visual cortex |
| VISpl | Posterolateral visual area |
| VISpm | Posteromedial visual area |
| VISpor | Postrhinal area |
| VISrl | Rostrolateral visual area |
| PA | Posterior amygdalar nucleus |
| PAA | Piriform-amygdalar area |
| MEA | Medial amygdalar nucleus |
| LA | Lateral amygdalar nucleus |

| | |
|---|---|
| IA | Intercalated amygdalar nucleus |
| COA | Cortical amygdalar area |
| AAA | Anterior amygdalar area |
| BLAd | Basolateral amygdalar nucleus, dorsal part |
| BLAv | Basolateral amygdalar nucleus, ventral part |
| BMA | Basomedial amygdalar nucleus |
| CEA | Central amygdalar nucleus |
| EP | Endopiriform nucleus |
| HATA | Hippocampo-amygdalar transition area |
| CA1d | CA1 subfield, dorsal part |
| CA1i | CA1 subfield, intermediate |
| CA1v | CA1 subfield, ventral |
| CA2d | CA2 subfield, dorsal part |
| CA2i | CA2 subfield, intermediate |
| CA2v | CA2 subfield, ventral |
| CA3d | CA3 subfield, dorsal part |
| CA3i | CA3 subfield, intermediate |
| CA3v | CA3 subfield, ventral |
| DGd | Dentate gyrus, dorsal part |
| DGi | Dentate gyrus, intermediate |
| DGv | Dentate gyrus, ventral |
| ENTl | Entorhinal area, lateral part |
| ENTm | Entorhinal area, medial part |
| PAR | Parasubiculum |
| POST | Postsubiculum |
| PRE | Presubiculum |
| ProS | Prosubiculum |
| SUB | Subiculum |
| ACB | Nucleus accumbens |
| BST | Bed nuclei of the stria terminalis |
| CLA | Claustrum |
| CPa | Caudoputamen, anterior ventral lateral |
| CPm | Caudoputamen, medial ventral medial |
| CPc | Caudoputamen, caudal ventral |
| FS | Fundus of striatum |
| GPe | Globus pallidus, external segment |
| GPi | Globus pallidus, internal segment |
| LSd | Lateral septal nucleus, dorsal |
| LSv | Lateral septal nucleus, ventral part |
| MS | Medial septal nucleus |
| MA | Magnocellular nucleus |
| NDB | Diagonal band nucleus |
| PAL | Pallidum |
| SF | Septofimbrial nucleus |
| SI | Substantia innominata |
| STR | Striatum unassigned |
| TRS | Triangular nucleus of septum |
| AD | Anterodorsal nucleus |

| | |
|---|---|
| AM | Anteromedial nucleus |
| AV | Anteroventral nucleus of thalamus |
| CL | Central lateral nucleus of the thalamus |
| CM | Central medial nucleus of the thalamus |
| Eth | Ethmoid nucleus of the thalamus |
| IAt | Interanteromedial nucleus of the thalamus |
| IGL | Intergeniculate leaflet of the lateral geniculate complex |
| IMD | Intermediodorsal nucleus of the thalamus |
| LD | Lateral dorsal nucleus of thalamus |
| LGd | Dorsal part of the lateral geniculate complex |
| LGv | Ventral part of the lateral geniculate complex |
| LPcl | Lateral posterior nucleus of the thalamus, caudal lateral |
| LPam | Lateral posterior nucleus of the thalamus, anterior medial |
| MD | Mediodorsal nucleus of thalamus |
| MG | Medial geniculate complex |
| PCN | Paracentral nucleus |
| PF | Parafascicular nucleus |
| PO | Posterior complex of the thalamus |
| PPnT | Posterior Paralaminar nuclei of the thalamus |
| PP | Peripeduncular nucleus |
| PR | Perireunensis nucleus |
| PT | Parataenial nucleus |
| PVT | Paraventricular nucleus of the thalamus |
| RE | Nucleus of reuniens |
| RH | Rhomboid nucleus |
| RT | Reticular nucleus of the thalamus |
| SMT | Submedial nucleus of the thalamus |
| SPA | Subparafascicular area |
| SPFm | Subparafascicular nucleus, magnocellular part |
| VAL | Ventral anterior-lateral complex of the thalamus |
| VM | Ventral medial nucleus of the thalamus |
| VPL | Ventral posterolateral nucleus of the thalamus |
| VPM | Ventral posteromedial nucleus of the thalamus |
| VPMpc | Ventral posteromedial nucleus of the thalamus, parvicellular part |
| Xi | Xiphoid thalamic nucleus |
| AHN | Anterior hypothalamic nucleus |
| AVP | Anteroventral preoptic nucleus |
| AVPV | Anteroventral periventricular nucleus |
| DMH | Dorsomedial nucleus of the hypothalamus |
| FF | Fields of Forel |
| LHA | Lateral hypothalamic area |
| LM | Lateral mammillary nucleus |
| LPO | Lateral preoptic area |
| ME | Median eminence |
| PON | Median preoptic nucleus |
| Mmm | Medial mammillary nuclei |
| PH | Posterior hypothalamic nucleus |
| PMH | Ventral premammillary nucleus |

| | |
|---|---|
| PS | Parastrial nucleus |
| STN | Subthalamic nucleus |
| PVH | Paraventricular hypothalamic nucleus |
| PVHd | Paraventricular hypothalamic nucleus, descending division |
| PeVH | Arcuate hypothalamic nucleus |
| PeF | Perifornical nucleus |
| RCH | Retrochiasmatic area |
| SO | Supraoptic nucleus |
| SUM | Supramammillary nucleus |
| TM | Tuberomammillary nucleus |
| TU | Tuberal nucleus |
| VL/VMPO | Ventrolateral and ventromedial preoptic nucleus |
| VMH | Ventromedial hypothalamic nucleus |
| ZI | Zona incerta |
| HBn | Medial habenula |
| RPF | Retroparafascicular nucleus |
| APN | Anterior pretectal nucleus |
| PTA | Posterior pretectal areas |
| NPC | Nucleus of the posterior commissure |
| PRC/SCO | Pre- and Sub-commissural organ |
| PPN | Pedunculopontine nucleus |
| CUN | Cuneiform nucleus |
| AT/VTN | Anterior and Ventral tegmental nucleus |
| CLI | Central linear nucleus raphe |
| DR | Dorsal nucleus raphe |
| IC | Inferior colliculus |
| IF | Interfascicular nucleus raphe |
| IPN | Interpeduncular nucleus |
| MRNal | Midbrain reticular nucleus, anterior lateral |
| MRNam | Midbrain reticular nucleus, anterior medial |
| MRNpl | Midbrain reticular nucleus, posterior lateral |
| MRNpm | Midbrain reticular nucleus, posterior medial |
| Tn | Lateral terminal nucleus of the accessory optic tract |
| NB | Nucleus of the brachium of the inferior colliculus |
| EW/MA3/ND | accesory oculomotor nuclei |
| PAGa | Periaqueductal gray, anterior part |
| PAGd | Periaqueductal gray, posterior dorsal |
| PAGv | Periaqueductal gray, posterior ventral |
| PAGl | Periaqueductal gray, medial lateral |
| PBG | Parabigeminal nucleus |
| Pa4/IV | Trochlear nucleus and Paratrochlear |
| RL | Rostral linear nucleus raphe |
| RN | Red nucleus |
| RR | Midbrain reticular nucleus, retrorubral area |
| SAG | Nucleus sagulum |
| SCd | Superior colliculus, deep layers, |
| Sci | Superior colliculus, intermediate layers |
| SCs | Superior colliculus, superficial layers |

| | |
|---|---|
| SN | Substantia nigra |
| VTA | Ventral tegmental area |
| AMB | Nucleus ambiguus |
| AP | Area postrema |
| CS | Superior central nucleus raphe |
| CU/GR | Cuneate and Gracile nucleus |
| DCO | Dorsal cochlear nucleus |
| DMX | Dorsal motor nucleus of the vagus nerve |
| DTN/PDTg | Dorsal and Posterodorsal tegmental nucleus |
| ECU | External cuneate nucleus |
| GRN | Gigantocellular reticular nucleus |
| I5/PC5 | Intertrigeminal and Parvicellular motor 5 nucleus |
| V | Motor nucleus of trigeminal |
| IO | Inferior olivary complex |
| IRN | Intermediate reticular nucleus |
| KF | Koelliker-Fuse subnucleus |
| LAC | Lateral vestibular nucleus |
| LC/B | Barrington's nucleus, Sublaterodorsal nucleus, subceruleus and Locus ceruleus |
| LDT | Laterodorsal tegmental nucleus |
| LIN | Linear nucleus of the medulla |
| LRN | Lateral reticular nucleus |
| MARN | Magnocellular reticular nucleus |
| MDRN | Medullary reticular nucleus |
| MV | Medial vestibular nucleus |
| NI | Nucleus incertus |
| NLL | Nucleus of the lateral lemniscus |
| NR/XII | Nucleus of Roller and Hypoglossal nucleus |
| NTB | Nucleus of the trapezoid body |
| NTS/PAS | Nucleus of the solitary tract and Parasolitary nucleus |
| P5/Acs5 | Peritrigeminal zone and Accessory trigeminal nucleus |
| PARN | Parvicellular reticular nucleus |
| PB | Parabrachial nucleus |
| PCG/SG | Pontine central gray and Supragenual nucleus |
| PG | Pontine gray |
| PGRN | Paragigantocellular reticular nucleus |
| POR | Superior olivary complex, periolivary region |
| PPY | Parapyramidal nucleus |
| PRNc | Pontine reticular nucleus, caudal |
| PRNr | Pontine reticular nucleus, rostral |
| PRP | Nucleus prepositus |
| PSV | Principal sensory nucleus of the trigeminal |
| Pa5 | Paratrigeminal nucleus |
| RM/RO | Nucleus raphe magnus and obscurus |
| RPA | Nucleus raphe pallidus |
| RPO | Nucleus raphe pontis |
| SOC | Superior olivary complex, medial part |
| SPIV | Spinal vestibular nucleus |

| | |
|---|---|
| SPVC | Spinal nucleus of the trigeminal, caudal part |
| SPVI | Spinal nucleus of the trigeminal, interpolar part |
| SPVO | Spinal nucleus of the trigeminal, oral part |
| SUT | Supratrigeminal nucleus |
| SUV | Superior vestibular nucleus |
| TRN | Tegmental reticular nucleus |
| VCO | Ventral cochlear nucleus |
| VII | Facial motor nucleus |
| x | Nucleus x |
| CB | Cerebellum unassigned |
| ANcr1 | Crus 1 |
| ANcr2 | Crus 2 |
| CENT2 | Lobule II |
| CENT3 | Lobule III |
| COPY | Copula pyramidis |
| DEC | Declive (VI) |
| DN | Dentate nucleus |
| FL | Flocculus |
| FOTU | Folium-tuber vermis (VII) |
| IP | Interposed nucleus |
| LING | Lingula (I) |
| NOD | Nodulus (X) |
| PFL | Paraflocculus |
| PRM | Paramedian lobule |
| PYR | Pyramus (VIII) |
| SIM | Simple lobule |
| UVU | Uvula (IX) |
| VeCB/ICB | Vestibulocerebellar and Infracerebellar nucleus |

#### Table S3 | Responsive areas - high-frequency stimulation

| | Early (0 - 2 s) | | | | Late (3 - 8 s) | | | |
|---|---|---|---|---|---|---|---|---|
| Brain Areas | CAMKII | NTSR | PV | GAD2 | CAMKII | NTSR | PV | GAD2 |
| ACAd | 1 | 1 | 1 | 0 | 1 | 0 | 1 | 0 |
| ACAv | 1 | 0 | 0 | -1 | 1 | 0 | 0 | -1 |
| Ala | 1 | 1 | 1 | 0 | -1 | -1 | 0 | 0 |
| AUDd | 1 | 0 | 1 | 0 | 0 | 0 | 0 | 0 |
| AUDp | 1 | 0 | 1 | 0 | 0 | 0 | 0 | 0 |
| AUDpo | 1 | 0 | 1 | 0 | 0 | 1 | 0 | 0 |
| AUDv | 1 | 0 | 1 | 0 | 0 | 0 | 1 | 0 |
| AON | 1 | 0 | 0 | 0 | 1 | 0 | 0 | 0 |
| APr | 1 | 0 | 0 | 0 | 1 | 0 | 0 | -1 |
| DP | 0 | 0 | 0 | 0 | 1 | -1 | -1 | 1 |
| ECT | 1 | 0 | 1 | 0 | 0 | 1 | 0 | 0 |
| GU | 1 | -1 | 1 | 0 | 0 | 0 | 0 | 0 |
| ILA | 1 | 0 | 1 | 0 | 0 | 0 | 0 | 0 |
| MOp | 1 | 1 | 0 | 0 | 0 | 0 | 0 | -1 |
| MOs | 1 | 0 | 1 | 0 | 0 | 1 | 1 | 0 |
| NLOT | 0 | 0 | 0 | 0 | 0 | 0 | 1 | 0 |
| OLF | 0 | 1 | 0 | 0 | 1 | 0 | -1 | 0 |
| ORB | 1 | 1 | 1 | 0 | 0 | 1 | 0 | 0 |
| OT | -1 | 0 | 1 | 0 | 1 | 0 | -1 | 0 |
| PERI | 1 | 0 | 1 | 0 | 0 | 0 | 0 | 0 |
| PIR | -1 | 1 | 0 | 0 | -1 | 0 | -1 | 0 |
| PL | 1 | 1 | 1 | 0 | 0 | 1 | 0 | 0 |
| RSPagl | 1 | 0 | 0 | 0 | 0 | 0 | 0 | 0 |
| RSPd | 0 | 0 | 0 | 0 | 0 | 0 | 0 | 0 |
| RSPv | 0 | 0 | -1 | 0 | 0 | -1 | 0 | 0 |
| SSp | 1 | 1 | 0 | 0 | 0 | 0 | 1 | 0 |
| SSs | 1 | 0 | 0 | 0 | 0 | 0 | 0 | 0 |
| TEa | 1 | 0 | 1 | 0 | 0 | 0 | 0 | 0 |
| TR | 0 | 0 | -1 | 0 | 1 | 0 | 0 | 0 |
| TT | 1 | 0 | 0 | 0 | 0 | 0 | -1 | 0 |
| VISC | 1 | 0 | 1 | 0 | 1 | 0 | 0 | 0 |
| VISa | 0 | 0 | 0 | 0 | 0 | 0 | 0 | 0 |
| VISal | 1 | 0 | 1 | 0 | 0 | 0 | 0 | 0 |
| VISam | 1 | 0 | 0 | 0 | 1 | 0 | 0 | 0 |
| VISl | 1 | 1 | 1 | 0 | 1 | 0 | 0 | 0 |
| VISli | 1 | 1 | 1 | 0 | 1 | 0 | 0 | 0 |
| VISp | 1 | 1 | 0 | 0 | 0 | 0 | 0 | 0 |
| VISpl | 1 | 1 | 1 | 0 | 1 | 1 | 0 | 1 |
| VISpm | 1 | 0 | 0 | 0 | 1 | 0 | 0 | 0 |
| VISpor | 1 | 1 | 1 | 0 | 1 | 1 | 1 | 0 |
| VISrl | 0 | 0 | 1 | 0 | 0 | 0 | 0 | 0 |
| PA | 0 | 1 | 0 | 0 | 1 | 0 | 0 | 0 |
| PAA | -1 | 1 | 0 | 0 | 1 | 0 | -1 | 0 |
| MEA | 0 | -1 | -1 | 0 | 0 | 0 | -1 | 0 |
| LA | 1 | 1 | 1 | 0 | 0 | 0 | 0 | 0 |
| IA | 0 | 0 | -1 | 0 | 0 | 0 | 0 | 0 |

| | | | | | | | | |
|---|---|---|---|---|---|---|---|---|
| COA | 0 | 1 | 0 | 0 | 1 | 1 | -1 | 0 |
| AAA | 0 | 0 | 0 | 0 | 0 | 0 | -1 | 0 |
| BLAd | 1 | 0 | 0 | 0 | 0 | 0 | 0 | 0 |
| BLAv | 0 | 0 | 0 | 0 | 1 | 0 | 0 | 0 |
| BMA | 0 | 1 | 0 | 0 | 0 | 0 | 0 | 0 |
| CEA | 1 | 1 | 0 | 0 | -1 | 0 | 0 | 0 |
| EP | 1 | 1 | 0 | 0 | 0 | 1 | 0 | 0 |
| HATA | 1 | 0 | 0 | 0 | 0 | 0 | 0 | 0 |
| CA1d | 1 | -1 | 0 | 0 | -1 | 1 | 1 | -1 |
| CA1i | 1 | 1 | 1 | 0 | 1 | 0 | 0 | 0 |
| CA1v | 1 | 0 | 1 | 0 | 1 | 1 | 0 | 0 |
| CA2d | 1 | -1 | 1 | 0 | 1 | -1 | 1 | 0 |
| CA2i | 1 | 1 | 1 | 0 | 1 | 0 | 0 | 0 |
| CA2v | 1 | 0 | 0 | 0 | 0 | 0 | 0 | 0 |
| CA3d | 1 | 0 | 0 | 0 | 1 | 1 | 0 | 0 |
| CA3i | 1 | 1 | 1 | 0 | 1 | -1 | 0 | 0 |
| CA3v | 1 | 0 | 0 | 0 | 0 | 0 | 0 | 0 |
| DGd | 1 | 0 | 1 | 1 | 1 | 0 | 0 | 0 |
| DGi | 1 | 0 | 0 | 0 | 0 | 0 | 0 | 0 |
| DGv | -1 | -1 | -1 | -1 | -1 | 0 | -1 | 0 |
| ENTl | 1 | 1 | 1 | 0 | 0 | 0 | 0 | 0 |
| ENTm | 1 | 1 | 0 | 0 | 0 | 0 | 0 | 0 |
| PAR | 0 | 0 | 0 | 0 | 0 | 0 | 1 | -1 |
| POST | 1 | 0 | 0 | 0 | 0 | 1 | 0 | -1 |
| PRE | 1 | 0 | -1 | -1 | 1 | -1 | -1 | -1 |
| ProS | 1 | 1 | 1 | 0 | -1 | 1 | 1 | -1 |
| SUB | 1 | 1 | 0 | 0 | 1 | -1 | 0 | 0 |
| ACB | 1 | 0 | 1 | 0 | 0 | 0 | 0 | 1 |
| BST | 1 | 0 | 1 | 0 | 0 | 0 | 1 | 0 |
| CLA | 1 | -1 | 1 | 0 | -1 | 0 | 0 | 0 |
| CPa | 1 | 1 | 1 | 0 | -1 | 0 | 1 | 0 |
| CPm | 1 | 0 | 1 | 0 | -1 | -1 | 1 | 0 |
| CPc | 1 | 1 | 1 | 0 | 1 | 0 | 1 | 0 |
| FS | -1 | -1 | -1 | 0 | 0 | 0 | 1 | 0 |
| GPe | 1 | 1 | 1 | 0 | -1 | 1 | 1 | 0 |
| GPi | 1 | 1 | 1 | 1 | 0 | 0 | 0 | 0 |
| LSd | 1 | 0 | 1 | 0 | 0 | 0 | 1 | 0 |
| LSv | 1 | 0 | 0 | 1 | 1 | 0 | 0 | 0 |
| MS | 1 | 0 | 1 | 0 | 0 | 0 | 0 | 0 |
| MA | 0 | 0 | 0 | 0 | 1 | 0 | -1 | 0 |
| NDB | 1 | 0 | 0 | 0 | 1 | 0 | -1 | 0 |
| PAL | 1 | 0 | 0 | 0 | 0 | 0 | -1 | 0 |
| SF | 1 | 0 | 0 | 0 | 0 | 1 | 0 | 0 |
| SI | 1 | 0 | 0 | 0 | 0 | 0 | 1 | 0 |
| STR | 1 | 1 | 0 | 0 | 0 | 0 | 1 | 0 |
| TRS | 0 | 0 | 0 | 0 | -1 | 0 | 0 | 0 |
| AD | 1 | 0 | 0 | 0 | 0 | 0 | 0 | 0 |
| AM | 1 | 1 | 1 | 0 | 0 | 0 | 1 | 0 |
| AV | 1 | 1 | 1 | 0 | 1 | 0 | -1 | 0 |
| CL | 0 | 1 | 0 | 0 | 0 | 0 | 1 | 0 |

| | | | | | | | | |
|---|---|---|---|---|---|---|---|---|
| CM | 1 | 0 | 0 | 0 | 1 | 0 | 0 | 0 |
| Eth | 0 | 1 | 0 | 0 | 0 | 0 | 0 | 0 |
| IAt | 1 | 0 | 1 | 0 | 0 | 1 | 1 | 0 |
| IGL | 1 | 1 | 1 | 1 | 1 | 1 | 0 | 1 |
| IMD | 0 | 0 | 0 | 0 | 0 | 0 | 0 | 0 |
| LD | 1 | 1 | 0 | 0 | 0 | 0 | 0 | 0 |
| LGd | 1 | 1 | 1 | 1 | 1 | 1 | 0 | 1 |
| LGv | 1 | 1 | 1 | 1 | 1 | 1 | 0 | 1 |
| LPcl | 1 | 1 | 0 | 0 | 0 | 0 | 0 | 0 |
| LPam | 1 | 1 | 0 | 0 | 1 | 0 | 1 | 0 |
| MD | 0 | 0 | 1 | 0 | 0 | 0 | 0 | 0 |
| MG | 1 | 1 | 1 | 1 | 1 | 1 | 0 | -1 |
| PCN | 1 | 0 | 0 | 0 | 0 | 0 | 0 | 0 |
| PF | 1 | 1 | 1 | 0 | 0 | 0 | 1 | 0 |
| PO | 1 | 1 | 0 | 0 | 1 | 0 | 0 | 0 |
| PPnT | 1 | 1 | 1 | 1 | 1 | 1 | 0 | 1 |
| PP | 1 | 1 | 1 | 0 | 1 | 0 | 0 | 0 |
| PR | 1 | 1 | 0 | 0 | 0 | 0 | 0 | 0 |
| PT | 1 | 0 | 1 | 0 | 0 | 0 | 0 | 0 |
| PVT | 1 | 1 | 0 | 0 | 1 | 0 | 0 | 0 |
| RE | 1 | 1 | 1 | 0 | 0 | 0 | 0 | 0 |
| RH | 0 | 0 | 0 | 0 | 1 | 0 | 0 | 0 |
| RT | 1 | 1 | 1 | 0 | 0 | 1 | 0 | 1 |
| SMT | 1 | 1 | 1 | 0 | 0 | 0 | 0 | 0 |
| SPA | 1 | 1 | 0 | 0 | 1 | 0 | 0 | 0 |
| SPFm | 1 | 1 | 0 | 0 | 0 | 0 | 0 | 0 |
| VAL | 1 | 1 | 1 | 0 | 1 | 0 | 0 | 0 |
| VM | 1 | 1 | 1 | 0 | 0 | 0 | 0 | 0 |
| VPL | 1 | 1 | 1 | 0 | 0 | 1 | 0 | 1 |
| VPM | 1 | 1 | 1 | 0 | 0 | 1 | 1 | 0 |
| VPMpc | 1 | 1 | 0 | 0 | 0 | 1 | 0 | -1 |
| Xi | 1 | 1 | 0 | 0 | 0 | 0 | 1 | 0 |
| AHN | 1 | 0 | 1 | 0 | 0 | 1 | 0 | 0 |
| AVP | 1 | 0 | 0 | 0 | 0 | 0 | 0 | 0 |
| AVPV | 0 | -1 | 0 | 0 | 1 | 1 | 0 | 0 |
| DMH | 0 | 0 | 0 | 0 | 1 | 1 | 0 | 0 |
| FF | 1 | 1 | 1 | 0 | 0 | 0 | 1 | 0 |
| LHA | 1 | 1 | 1 | 0 | 0 | 0 | 0 | 0 |
| LM | 1 | 0 | 0 | 0 | 0 | 0 | 0 | 0 |
| LPO | 1 | 0 | 0 | 1 | 0 | 0 | 0 | 1 |
| ME | 0 | 0 | 0 | 0 | 1 | 1 | 0 | 0 |
| PON | 1 | 0 | 1 | 0 | 0 | 0 | 0 | 1 |
| Mmm | 1 | 0 | 0 | -1 | 0 | 0 | 0 | 1 |
| PH | 1 | 1 | 0 | 0 | 0 | 0 | 0 | 0 |
| PMH | 0 | 0 | 0 | 0 | 1 | 1 | 0 | 0 |
| PS | 1 | 1 | 0 | 0 | 0 | 0 | 0 | 0 |
| STN | 1 | 1 | 1 | 0 | 1 | 0 | 0 | 0 |
| PVH | 1 | 1 | 0 | 0 | 0 | 0 | 0 | 0 |
| PVHd | 1 | 1 | 0 | 0 | 0 | 0 | 0 | 0 |
| PeVH | 1 | 0 | 0 | 0 | 1 | 1 | 0 | 0 |

| | | | | | | | | |
|---|---|---|---|---|---|---|---|---|
| PeF | 1 | 1 | 1 | 0 | 0 | 0 | 0 | 0 |
| RCH | 0 | 0 | 0 | 0 | 1 | 1 | 0 | 0 |
| SO | 0 | 0 | 0 | 0 | 1 | 0 | -1 | 0 |
| SUM | 1 | 0 | 0 | 0 | 0 | 0 | 0 | 1 |
| TM | -1 | -1 | -1 | 0 | -1 | 1 | -1 | 0 |
| TU | 0 | 0 | 0 | 0 | 1 | 0 | 0 | 0 |
| VL/VMPO | 0 | 0 | 0 | 0 | 1 | 1 | 1 | 0 |
| VMH | 0 | 0 | 0 | 0 | 1 | 0 | 1 | 0 |
| ZI | 1 | 1 | 1 | 1 | 1 | 1 | 1 | 1 |
| HBn | 1 | 1 | 0 | 0 | 0 | 0 | 0 | 0 |
| RPF | 1 | 1 | 1 | 0 | 0 | 0 | 1 | 0 |
| APN | 1 | 1 | 1 | 1 | -1 | 1 | 0 | 1 |
| PTA | 1 | 1 | 1 | 1 | 0 | 1 | 0 | 1 |
| NPC | 1 | 1 | 1 | 0 | 1 | 0 | 0 | 1 |
| PRC/SCO | 1 | 1 | 0 | 0 | 1 | 0 | 0 | 0 |
| PPN | 1 | 1 | 1 | 0 | 1 | 1 | 0 | 0 |
| CUN | 1 | 1 | 1 | 0 | 0 | 0 | 1 | 1 |
| AT/VTN | 1 | 1 | 1 | 0 | 0 | 0 | 0 | 1 |
| CLI | 1 | 0 | 1 | 0 | 0 | 1 | 1 | 0 |
| DR | 1 | 0 | 1 | 0 | 0 | 0 | 0 | 0 |
| IC | 0 | 1 | 1 | 1 | 0 | 0 | 1 | 1 |
| IF | 0 | 0 | 0 | 0 | 1 | 0 | 1 | 1 |
| IPN | -1 | 1 | -1 | -1 | 0 | 0 | -1 | -1 |
| MRNal | 1 | 1 | 1 | 0 | 1 | 1 | 1 | 1 |
| MRNam | 1 | 1 | 1 | 1 | 1 | 1 | 1 | 1 |
| MRNpl | 1 | 1 | 1 | 1 | 1 | 1 | 1 | -1 |
| MRNpm | 1 | 1 | 1 | 1 | 1 | 1 | 1 | 1 |
| Tn | 1 | 1 | 0 | 1 | 0 | 0 | 0 | 0 |
| NB | -1 | 0 | 0 | 0 | 1 | 1 | 0 | 1 |
| EW/MA3/ND | 1 | 1 | 1 | 0 | 0 | 0 | 0 | 0 |
| PAGa | 1 | 1 | 1 | 0 | 0 | 0 | 1 | 1 |
| PAGd | 1 | 1 | 1 | 1 | 1 | 1 | 1 | 1 |
| PAGv | 1 | 1 | 1 | 0 | 1 | 0 | 0 | 1 |
| PAGl | 1 | 1 | 1 | 1 | 1 | 1 | 1 | 1 |
| PBG | 0 | 0 | 0 | 0 | 1 | -1 | -1 | 0 |
| Pa4/IV | 1 | 0 | 1 | 0 | 1 | 0 | 0 | 0 |
| RL | 1 | 0 | 0 | 0 | 0 | 0 | 0 | 0 |
| RN | 1 | 1 | 1 | 1 | 0 | 1 | 1 | 0 |
| RR | 1 | 1 | 1 | 0 | 1 | 0 | 0 | 0 |
| SAG | 1 | 1 | 0 | 0 | 0 | 0 | 0 | 0 |
| SCd | 1 | 1 | 1 | 1 | 1 | 1 | 1 | 1 |
| SCi | 1 | 1 | 1 | 1 | 0 | 1 | 0 | 1 |
| SCs | 1 | 1 | 1 | 1 | 1 | 0 | 0 | 1 |
| SN | 1 | 1 | 0 | 0 | 0 | 0 | 0 | 0 |
| VTA | 1 | 1 | 0 | 0 | 0 | 0 | 0 | 0 |
| AMB | 0 | 0 | 0 | 0 | 0 | 0 | 0 | 0 |
| AP | -1 | -1 | -1 | -1 | -1 | 1 | -1 | 1 |
| CS | 1 | 1 | 1 | 0 | 0 | 0 | 1 | 1 |
| CU/GR | -1 | -1 | -1 | -1 | -1 | 1 | -1 | 1 |
| DCO | 0 | 1 | 0 | 0 | 1 | 0 | 1 | 0 |

| | | | | | | | | |
|---|---|---|---|---|---|---|---|---|
| DMX | 0 | 0 | 0 | 0 | 1 | 0 | -1 | 0 |
| DTN/PDTg | 1 | 1 | 1 | 0 | 0 | 1 | 0 | 1 |
| ECU | -1 | -1 | -1 | -1 | -1 | 1 | -1 | 1 |
| GRN | 1 | 0 | 1 | 0 | 1 | 0 | 0 | 1 |
| I5/PC5 | 0 | 0 | 0 | 0 | 1 | 0 | 0 | 0 |
| V | 1 | 0 | 0 | 0 | 1 | 0 | 0 | 0 |
| IO | 0 | 0 | 0 | 0 | 1 | 1 | -1 | 1 |
| IRN | 0 | 0 | 0 | 0 | 1 | 0 | 0 | 0 |
| KF | 1 | 0 | 0 | 0 | 1 | 0 | 0 | 0 |
| LAV | 1 | 0 | 0 | 0 | 1 | 0 | 0 | 0 |
| LC/B | 1 | 0 | 0 | 0 | 0 | 1 | 0 | 0 |
| LDT | 1 | 0 | 1 | 0 | 0 | 1 | 0 | 0 |
| LIN | 0 | 0 | 0 | 0 | 1 | 0 | 0 | 0 |
| LRN | 0 | 0 | 0 | 0 | 1 | 0 | 1 | 0 |
| MARN | 1 | 0 | 0 | 0 | 1 | 0 | 1 | 0 |
| MDRN | -1 | -1 | -1 | -1 | -1 | 1 | -1 | 1 |
| MV | 0 | 0 | 0 | 0 | 1 | 1 | 1 | 1 |
| NI | 1 | 1 | 1 | 0 | 0 | 0 | 0 | 1 |
| NLL | 0 | 0 | 0 | 0 | 0 | 0 | 0 | 0 |
| NR/XII | -1 | -1 | -1 | -1 | -1 | 1 | -1 | 1 |
| NTB | 0 | 0 | 0 | 0 | 1 | 0 | 0 | 0 |
| NTS/PAS | 0 | 0 | 0 | 0 | 1 | 0 | 0 | 0 |
| P5/Acs5 | 1 | 0 | 1 | 0 | 1 | 0 | 0 | 0 |
| PARN | 0 | 0 | 0 | 0 | 1 | 0 | 0 | 1 |
| PB | 1 | 1 | 1 | 0 | 0 | 0 | 0 | 0 |
| PCG/SG | 1 | 1 | 1 | 0 | 0 | 1 | 0 | 1 |
| PG | 0 | 0 | 0 | 0 | 1 | 0 | 0 | 0 |
| PGRN | 0 | 0 | 1 | 1 | 1 | 0 | 0 | 1 |
| POR | 0 | 0 | 0 | 0 | 1 | 0 | 0 | 0 |
| PPY | 0 | 0 | 0 | 0 | 1 | 1 | 0 | 0 |
| PRNc | 1 | 1 | 1 | 1 | 0 | 0 | 0 | 0 |
| PRNr | 1 | 1 | 1 | 1 | 0 | 1 | 1 | 0 |
| PRP | 0 | 0 | 0 | 0 | 1 | 0 | 1 | 1 |
| PSV | 1 | 0 | 1 | 0 | 0 | 0 | 0 | 0 |
| Pa5 | -1 | -1 | -1 | -1 | -1 | 1 | -1 | 1 |
| RM/RO | 0 | 0 | 1 | 0 | 1 | 0 | 0 | 0 |
| RPA | 0 | 0 | 0 | 0 | 1 | 0 | 0 | 1 |
| RPO | 1 | 1 | 1 | 0 | 0 | 0 | 0 | 0 |
| SOC | 0 | 1 | 0 | 0 | 1 | 0 | 0 | 0 |
| SPIV | 0 | 0 | 0 | 0 | 1 | 0 | 0 | 0 |
| SPVC | -1 | -1 | -1 | -1 | -1 | 1 | -1 | 1 |
| SPVI | 0 | 0 | 0 | 0 | 1 | 0 | 1 | 0 |
| SPVO | 0 | 0 | 0 | 0 | 1 | 0 | 1 | 0 |
| SUT | 1 | 0 | 0 | 0 | 1 | 0 | 0 | 0 |
| SUV | 1 | 0 | 0 | 0 | 1 | 0 | 0 | 0 |
| TRN | 1 | 1 | 1 | 0 | 0 | 0 | 0 | 0 |
| VCO | 1 | 1 | 0 | 0 | 1 | 0 | 0 | 0 |
| VII | 0 | 1 | 0 | 0 | 1 | 0 | 1 | 0 |
| x | 0 | 0 | 0 | 0 | 0 | 0 | 1 | 0 |
| CB | 0 | -1 | 1 | 0 | 1 | 0 | 1 | 0 |

| | | | | | | | | |
|---|---|---|---|---|---|---|---|---|
| ANcr1 | 0 | -1 | -1 | 0 | 1 | -1 | -1 | 0 |
| ANcr2 | 0 | 0 | 0 | 0 | 1 | 1 | 0 | 0 |
| CENT2 | 1 | 1 | 1 | 0 | 1 | 0 | 1 | 1 |
| CENT3 | 1 | 1 | 1 | 1 | 1 | 0 | 1 | 1 |
| COPY | 1 | 1 | 0 | 0 | 1 | 0 | 1 | 0 |
| DEC | 1 | 0 | 0 | 0 | 1 | 0 | 1 | 0 |
| DN | 1 | 0 | 0 | 0 | 0 | 0 | 0 | 0 |
| FL | 0 | 1 | 0 | 0 | 1 | 0 | -1 | 0 |
| FOTU | -1 | -1 | -1 | -1 | -1 | 1 | -1 | 1 |
| IP | 1 | 1 | 0 | 0 | 0 | 0 | 1 | 1 |
| LING | 0 | 0 | 0 | 0 | 1 | -1 | -1 | 1 |
| NOD | 0 | 0 | 0 | 0 | 1 | 1 | 1 | 0 |
| PFL | 0 | 1 | 0 | 0 | 1 | 1 | -1 | 0 |
| PRM | 1 | 1 | 0 | 0 | 0 | 0 | -1 | 0 |
| PYR | -1 | -1 | -1 | -1 | -1 | 1 | -1 | 1 |
| SIM | 0 | 0 | 0 | 0 | 0 | 0 | 0 | 0 |
| UVU | 0 | 0 | 0 | 0 | 1 | 0 | 0 | 0 |
| VeCB/ICB | 0 | 0 | 0 | 0 | 1 | -1 | 0 | 0 |

#### Table S4 | Responsive areas - low-frequency stimulation

| Brain Areas | Early (0 - 2 s) CAMKII | NTSR | PV | GAD2 | Late (3 - 8 s) CAMKII | NTSR | PV | GAD2 |
|---|---|---|---|---|---|---|---|---|
| ACAd | | 0 | 0 | 0 | | 0 | 0 | 0 |
| ACAv | | 1 | 0 | 0 | | 0 | 0 | 0 |
| Ala | | 0 | 0 | 0 | | 0 | 0 | 0 |
| AUDd | | 0 | 0 | 0 | | 0 | 0 | 0 |
| AUDp | | 0 | 0 | 0 | | 0 | 0 | 0 |
| AUDpo | | 0 | 0 | 0 | | 0 | 0 | 0 |
| AUDv | | 0 | 1 | 0 | | 0 | 1 | 0 |
| AON | | 1 | 0 | 0 | | 0 | 1 | 0 |
| APr | | 0 | 0 | 0 | | 1 | 1 | 0 |
| DP | | 1 | 0 | 0 | | 0 | 0 | 0 |
| ECT | | 1 | 1 | 0 | | 1 | 1 | 0 |
| GU | | 1 | 0 | 0 | | 1 | 0 | 0 |
| ILA | | 0 | 1 | 0 | | 0 | 0 | 0 |
| MOp | | 0 | 0 | 0 | | 0 | 0 | 0 |
| MOs | | 1 | 0 | 0 | | 0 | 0 | 0 |
| NLOT | | 1 | 1 | 0 | | 0 | 1 | 0 |
| OLF | | 1 | 0 | 0 | | 1 | 1 | 0 |
| ORB | | 0 | 0 | 0 | | 0 | 0 | 0 |
| OT | | 0 | 0 | 0 | | 1 | 0 | 0 |
| PERI | | 0 | 0 | 0 | | 0 | 0 | 0 |
| PIR | | 0 | 0 | 0 | | 1 | 0 | 0 |
| PL | | 1 | 0 | 0 | | 0 | 0 | 0 |
| RSPagl | | 0 | 0 | 0 | | 1 | 1 | 0 |
| RSPd | | 1 | 0 | 0 | | 0 | 1 | 0 |
| RSPv | | 0 | 0 | 0 | | 0 | 0 | 0 |
| SSp | | 1 | 1 | 0 | | 0 | 0 | 0 |
| SSs | | 1 | 0 | 0 | | 0 | 0 | 0 |
| TEa | | 1 | 0 | 0 | | 0 | 0 | 0 |
| TR | | 0 | 0 | 0 | | 1 | 0 | 0 |
| TT | | 0 | 0 | 0 | | 1 | 1 | 0 |
| VISC | | 1 | 0 | 0 | | 0 | 0 | 0 |
| VISa | | 1 | 0 | 0 | | 0 | 0 | 0 |
| VISal | | 1 | 1 | 0 | | 1 | 0 | 0 |
| VISam | | 1 | 0 | 0 | | 0 | 0 | 0 |
| VISl | | 1 | 0 | 0 | | 1 | 1 | 0 |
| VISli | | 0 | 0 | 0 | | 1 | 0 | 0 |
| VISp | | 1 | 0 | 0 | | 1 | 1 | 1 |
| VISpl | | 0 | 1 | 0 | | 1 | 1 | 0 |
| VISpm | | 1 | 0 | 0 | | 0 | 0 | 0 |
| VISpor | | 0 | 0 | 0 | | 1 | 0 | 0 |
| VISrl | | 1 | 0 | 0 | | 1 | 0 | 0 |
| PA | | 0 | 0 | 0 | | 1 | 0 | 0 |
| PAA | | 0 | 0 | 0 | | 1 | 0 | 0 |
| MEA | | 1 | 0 | 0 | | 1 | 0 | 0 |
| LA | | 1 | 0 | 0 | | 1 | 0 | 0 |
| IA | | 1 | 0 | 0 | | 1 | 0 | 0 |

| | | | | | | |
|---|---|---|---|---|---|---|
| COA | 0 | 0 | 0 | 1 | 1 | 0 |
| AAA | 1 | 0 | 0 | 1 | 1 | 0 |
| BLAd | 0 | 0 | 0 | 0 | 0 | 0 |
| BLAv | 0 | 0 | 0 | 0 | 0 | 0 |
| BMA | 0 | 0 | 0 | 1 | 0 | 0 |
| CEA | 0 | 0 | 0 | 0 | 0 | 0 |
| EP | 0 | 0 | 0 | 0 | 0 | 0 |
| HATA | 0 | 0 | 0 | 0 | 0 | 0 |
| CA1d | 1 | 0 | 0 | 1 | 1 | 0 |
| CA1i | 0 | 1 | 0 | 0 | 0 | 0 |
| CA1v | 1 | 1 | 0 | 0 | 0 | 1 |
| CA2d | 0 | 0 | 0 | 1 | 1 | 0 |
| CA2i | 0 | 1 | 0 | 0 | 0 | 0 |
| CA2v | 1 | 0 | 0 | 1 | 0 | 0 |
| CA3d | 0 | 0 | 0 | 1 | 0 | 0 |
| CA3i | 1 | 1 | 0 | 0 | 0 | 0 |
| CA3v | 0 | 0 | 0 | 1 | 0 | 0 |
| DGd | 0 | 0 | 0 | 0 | 0 | 0 |
| DGi | 0 | 0 | 0 | 0 | 1 | 0 |
| DGv | 1 | 0 | 0 | 0 | 0 | 0 |
| ENTl | 0 | 0 | 0 | 0 | 0 | 0 |
| ENTm | 1 | 0 | 0 | 0 | 0 | 0 |
| PAR | 0 | 0 | 0 | 0 | 0 | 0 |
| POST | 0 | 0 | 0 | 0 | 0 | 0 |
| PRE | 0 | 0 | 0 | 1 | 0 | 0 |
| ProS | 1 | 0 | 0 | 1 | 0 | 0 |
| SUB | 0 | 0 | 0 | 1 | 1 | 0 |
| ACB | 1 | 0 | 0 | 0 | 0 | 0 |
| BST | 1 | 0 | 0 | 0 | 0 | 0 |
| CLA | 0 | 0 | 0 | 0 | 0 | 0 |
| CPa | 0 | 1 | 0 | 1 | 0 | 0 |
| CPm | 0 | 1 | 0 | 0 | 0 | 0 |
| CPc | 1 | 1 | 0 | 1 | 0 | 0 |
| FS | 0 | 0 | 0 | 1 | 1 | 0 |
| GPe | 1 | 1 | 0 | 1 | 0 | 0 |
| GPi | 1 | 1 | 0 | 1 | 0 | 0 |
| LSd | 0 | 1 | 0 | 0 | 0 | 0 |
| LSv | 0 | 0 | 0 | 0 | 0 | 0 |
| MS | 0 | 0 | 0 | 0 | 0 | 0 |
| MA | 1 | 0 | 0 | 1 | 1 | 0 |
| NDB | 1 | 0 | 0 | 1 | 0 | 0 |
| PAL | 1 | 0 | 0 | 1 | 0 | 0 |
| SF | 0 | 0 | 0 | 1 | 0 | 0 |
| SI | 1 | 0 | 0 | 1 | 0 | 0 |
| STR | 1 | 0 | 0 | 1 | 0 | 0 |
| TRS | 1 | 0 | 0 | 1 | 0 | 0 |
| AD | 1 | 0 | 0 | 0 | 0 | 0 |
| AM | 1 | 1 | 0 | 1 | 0 | 0 |
| AV | 1 | 1 | 0 | 1 | 0 | 0 |
| CL | 1 | 0 | 0 | 1 | 0 | 0 |

| | | | | | | |
|---|---|---|---|---|---|---|
| CM | 1 | 0 | 0 | 1 | 0 | 0 |
| Eth | 1 | 0 | 0 | 1 | 0 | 0 |
| IAt | 1 | 0 | 0 | 0 | 0 | 0 |
| IGL | 1 | 0 | 0 | 0 | 0 | 0 |
| IMD | 1 | 0 | 0 | 1 | 0 | 0 |
| LD | 1 | 1 | 0 | 0 | 1 | 0 |
| LGd | 1 | 0 | 0 | 1 | 1 | 0 |
| LGv | 1 | 1 | 0 | 0 | 1 | 0 |
| LPcl | 1 | 1 | 0 | 1 | 0 | 0 |
| LPam | 1 | 0 | 0 | 1 | 0 | 0 |
| MD | 1 | 0 | 0 | 1 | 0 | 0 |
| MG | 1 | 1 | 0 | 1 | 0 | 0 |
| PCN | 1 | 0 | 0 | 0 | 0 | 0 |
| PF | 1 | 0 | 0 | 1 | 0 | 0 |
| PO | 1 | 1 | 0 | 0 | 0 | 0 |
| PPnT | 1 | 1 | 0 | 1 | 0 | 1 |
| PP | 1 | 0 | 0 | 1 | 1 | 0 |
| PR | 1 | 1 | 0 | 1 | 0 | 0 |
| PT | 1 | 1 | 0 | 1 | 0 | 0 |
| PVT | 1 | 0 | 0 | 1 | 0 | 0 |
| RE | 1 | 1 | 0 | 1 | 1 | 0 |
| RH | 0 | 0 | 0 | 1 | 0 | 0 |
| RT | 1 | 1 | 0 | 1 | 0 | 0 |
| SMT | 1 | 1 | 0 | 1 | 0 | 0 |
| SPA | 1 | 1 | 0 | 1 | 0 | 0 |
| SPFm | 1 | 1 | 0 | 1 | 0 | 0 |
| VAL | 1 | 1 | 0 | 0 | 0 | 0 |
| VM | 1 | 0 | 0 | 0 | 0 | 0 |
| VPL | 1 | 0 | 0 | 0 | 0 | 0 |
| VPM | 1 | 1 | 0 | 0 | 0 | 0 |
| VPMpc | 1 | 1 | 0 | 1 | 0 | 0 |
| Xi | 1 | 1 | 0 | 1 | 0 | 0 |
| AHN | 1 | 0 | 0 | 1 | 1 | 0 |
| AVP | 0 | 0 | 0 | 0 | 0 | 0 |
| AVPV | 0 | 0 | 0 | 1 | 0 | 0 |
| DMH | 0 | 0 | 0 | 1 | 0 | 0 |
| FF | 1 | 0 | 0 | 1 | 0 | 0 |
| LHA | 1 | 1 | 0 | 1 | 1 | 0 |
| LM | 1 | 0 | 0 | 1 | 1 | 0 |
| LPO | 1 | 0 | 0 | 1 | 1 | 0 |
| ME | 0 | 0 | 0 | 1 | 1 | 0 |
| PON | 0 | 0 | 0 | 1 | 1 | 0 |
| Mmm | 0 | 0 | 0 | 1 | 1 | 0 |
| PH | 1 | 0 | 0 | 1 | 1 | 0 |
| PMH | 0 | 0 | 0 | 1 | 1 | 0 |
| PS | 0 | 0 | 0 | 0 | 0 | 0 |
| STN | 1 | 1 | 0 | 1 | 0 | 0 |
| PVH | 1 | 1 | 0 | 1 | 0 | 0 |
| PVHd | 1 | 1 | 0 | 1 | 1 | 0 |
| PeVH | 0 | 0 | 0 | 1 | 1 | 0 |

| | | | | | | |
|---|---|---|---|---|---|---|
| PeF | 1 | 0 | 0 | 1 | 0 | 0 |
| RCH | 0 | 0 | 0 | 1 | 1 | 0 |
| SO | 1 | 0 | 0 | 1 | 0 | 0 |
| SUM | 0 | 0 | 0 | 0 | 0 | 0 |
| TM | 1 | 1 | 0 | 1 | 1 | 0 |
| TU | 0 | 0 | 0 | 1 | 1 | 0 |
| VL/VMPO | 0 | 0 | 0 | 1 | 0 | 0 |
| VMH | 1 | 0 | 0 | 1 | 1 | 0 |
| ZI | 1 | 1 | 1 | 1 | 0 | 0 |
| HBn | 1 | 1 | 0 | 1 | 0 | 0 |
| RPF | 1 | 1 | 0 | 1 | 0 | 0 |
| APN | 1 | 0 | 0 | 1 | 0 | 0 |
| PTA | 1 | 0 | 0 | 1 | 0 | 0 |
| NPC | 1 | 1 | 0 | 1 | 0 | 0 |
| PRC/SCO | 1 | 0 | 0 | 1 | 1 | 0 |
| PPN | 1 | 1 | 0 | 1 | 0 | 0 |
| CUN | 1 | 1 | 0 | 1 | 0 | 0 |
| AT/VTN | 1 | 1 | 0 | 0 | 0 | 0 |
| CLI | 1 | 0 | 0 | 0 | 1 | 0 |
| DR | 1 | 0 | 0 | 1 | 0 | 0 |
| IC | 1 | 1 | 0 | 1 | 0 | 0 |
| IF | 1 | 0 | 0 | 0 | 1 | 0 |
| IPN | 1 | 1 | 0 | 1 | 1 | 0 |
| MRNal | 1 | 1 | 0 | 1 | 0 | 0 |
| MRNam | 1 | 1 | 0 | 1 | 0 | 0 |
| MRNpl | 1 | 1 | 0 | 1 | 0 | 1 |
| MRNpm | 1 | 1 | 0 | 1 | 0 | 0 |
| Tn | 1 | 0 | 0 | 1 | 0 | 0 |
| NB | 0 | 0 | 0 | 1 | 1 | 0 |
| EW/MA3/ND | 1 | 1 | 0 | 1 | 0 | 0 |
| PAGa | 1 | 1 | 0 | 1 | 0 | 0 |
| PAGd | 1 | 1 | 1 | 1 | 0 | 1 |
| PAGv | 1 | 1 | 0 | 1 | 0 | 0 |
| PAGl | 1 | 0 | 0 | 1 | 0 | 0 |
| PBG | 1 | 0 | 0 | 1 | 1 | 0 |
| Pa4/IV | 1 | 1 | 0 | 0 | 0 | 0 |
| RL | 0 | 0 | 0 | 0 | 1 | 0 |
| RN | 1 | 0 | 0 | 1 | 0 | 0 |
| RR | 1 | 0 | 0 | 1 | 0 | 1 |
| SAG | 1 | 0 | 0 | 1 | 0 | 0 |
| SCd | 1 | 1 | 1 | 1 | 0 | 1 |
| SCi | 1 | 1 | 1 | 1 | 0 | 1 |
| SCs | 1 | 0 | 1 | 1 | 1 | 1 |
| SN | 1 | 0 | 0 | 1 | 0 | 0 |
| VTA | 1 | 0 | 0 | 1 | 0 | 0 |
| AMB | 0 | 0 | 0 | 1 | 0 | 0 |
| AP | 1 | 0 | 1 | 1 | 1 | 0 |
| CS | 1 | 1 | 0 | 0 | 0 | 0 |
| CU/GR | 1 | 0 | 1 | 1 | 1 | 0 |
| DCO | 1 | 0 | 0 | 1 | 0 | 0 |

| | | | | | | |
|---|---|---|---|---|---|---|
| DMX | 0 | 0 | 0 | 0 | 0 | 0 |
| DTN/PDTg | 1 | 1 | 0 | 1 | 0 | 0 |
| ECU | 1 | 0 | 1 | 1 | 1 | 0 |
| GRN | 1 | 0 | 0 | 1 | 0 | 0 |
| I5/PC5 | 0 | 0 | 0 | 1 | 0 | 0 |
| V | 0 | 0 | 0 | 1 | 0 | 0 |
| IO | 0 | 0 | 0 | 1 | 0 | 0 |
| IRN | 0 | 0 | 0 | 1 | 0 | 0 |
| KF | 0 | 0 | 0 | 1 | 0 | 0 |
| LAV | 0 | 0 | 0 | 1 | 0 | 0 |
| LC/B | 1 | 0 | 0 | 1 | 0 | 0 |
| LDT | 1 | 1 | 0 | 1 | 0 | 0 |
| LIN | 0 | 0 | 0 | 1 | 0 | 0 |
| LRN | 0 | 0 | 0 | 1 | 0 | 0 |
| MARN | 0 | 0 | 0 | 1 | 0 | 0 |
| MDRN | 1 | 0 | 1 | 1 | 1 | 0 |
| MV | 0 | 0 | 0 | 1 | 0 | 0 |
| NI | 1 | 0 | 0 | 1 | 0 | 0 |
| NLL | 0 | 0 | 0 | 1 | 1 | 0 |
| NR/XII | 1 | 0 | 1 | 1 | 1 | 0 |
| NTB | 0 | 0 | 0 | 1 | 0 | 0 |
| NTS/PAS | 0 | 0 | 0 | 1 | 0 | 0 |
| P5/Acs5 | 1 | 0 | 0 | 0 | 0 | 0 |
| PARN | 0 | 0 | 0 | 1 | 0 | 0 |
| PB | 1 | 0 | 0 | 1 | 0 | 0 |
| PCG/SG | 1 | 1 | 0 | 1 | 0 | 0 |
| PG | 0 | 0 | 0 | 1 | 0 | 0 |
| PGRN | 0 | 0 | 0 | 1 | 0 | 0 |
| POR | 0 | 0 | 0 | 1 | 0 | 0 |
| PPY | 0 | 0 | 0 | 1 | 0 | 0 |
| PRNc | 1 | 0 | 0 | 1 | 0 | 0 |
| PRNr | 1 | 1 | 0 | 1 | 0 | 0 |
| PRP | 1 | 1 | 0 | 1 | 1 | 0 |
| PSV | 1 | 0 | 0 | 0 | 1 | 0 |
| Pa5 | 1 | 0 | 1 | 1 | 1 | 0 |
| RM/RO | 1 | 0 | 0 | 1 | 1 | 0 |
| RPA | 0 | 0 | 0 | 1 | 1 | 0 |
| RPO | 1 | 0 | 0 | 1 | 1 | 0 |
| SOC | 0 | 0 | 0 | 1 | 0 | 0 |
| SPIV | 0 | 0 | 0 | 1 | 0 | 0 |
| SPVC | 1 | 0 | 1 | 1 | 1 | 0 |
| SPVI | 0 | 0 | 0 | 1 | 0 | 0 |
| SPVO | 0 | 0 | 0 | 1 | 0 | 0 |
| SUT | 1 | 0 | 0 | 1 | 0 | 0 |
| SUV | 1 | 0 | 0 | 1 | 0 | 0 |
| TRN | 1 | 1 | 0 | 1 | 0 | 0 |
| VCO | 0 | 0 | 0 | 1 | 1 | 0 |
| VII | 0 | 0 | 0 | 1 | 0 | 0 |
| x | 0 | 0 | 0 | 1 | 0 | 0 |
| CB | 1 | 0 | 0 | 1 | 1 | 0 |

| | | | | | | |
|---|---|---|---|---|---|---|
| ANcr1 | 1 | 0 | 0 | 1 | 0 | 0 |
| ANcr2 | 0 | 0 | 0 | 1 | 1 | 0 |
| CENT2 | 1 | 1 | 0 | 1 | 0 | 0 |
| CENT3 | 1 | 1 | 0 | 1 | 0 | 0 |
| COPY | 1 | 0 | 0 | 1 | 0 | 0 |
| DEC | 1 | 0 | 0 | 1 | 0 | 0 |
| DN | 1 | 0 | 0 | 1 | 1 | 0 |
| FL | 1 | 0 | 0 | 1 | 1 | 0 |
| FOTU | 1 | 0 | 1 | 1 | 1 | 0 |
| IP | 0 | 0 | 0 | 1 | 0 | 0 |
| LING | 0 | 0 | 0 | 1 | 1 | 0 |
| NOD | 0 | 0 | 0 | 1 | 1 | 0 |
| PFL | 1 | 1 | 0 | 1 | 1 | 0 |
| PRM | 1 | 0 | 0 | 1 | 1 | 0 |
| PYR | 1 | 0 | 1 | 1 | 1 | 0 |
| SIM | 1 | 0 | 0 | 1 | 0 | 0 |
| UVU | 1 | 1 | 0 | 1 | 1 | 0 |
| VeCB/ICB | 1 | 0 | 0 | 1 | 0 | 0 |
