## Supplementary figures and images for "Brain-wide mapping of neural activity mediating collicular-dependent behaviors"

### Movie S8

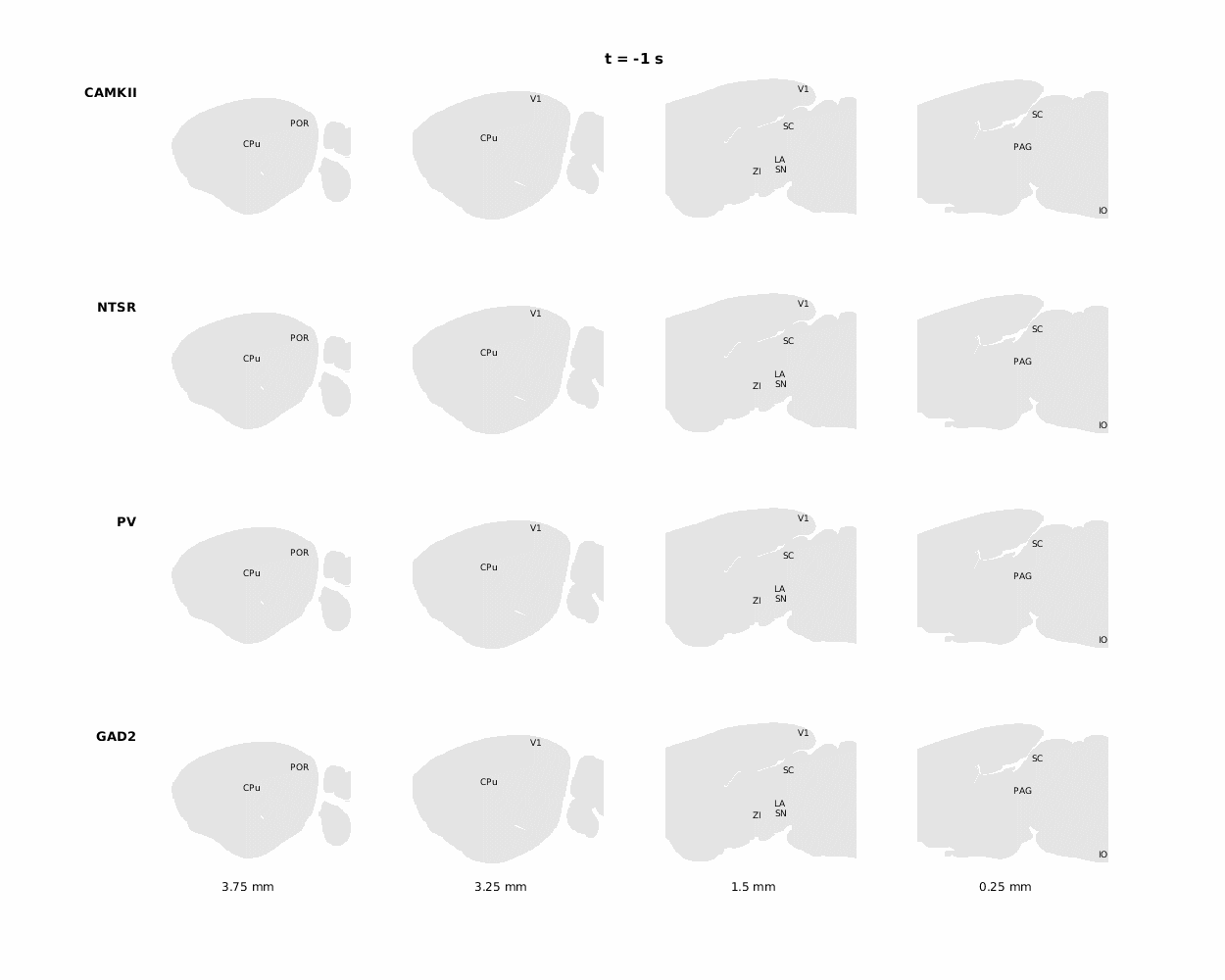
